# Dextran sodium sulfate-induced colitis alters the proportion and composition of replicating gut bacteria

**DOI:** 10.1101/2024.02.01.578403

**Authors:** Eve T. Beauchemin, Claire Hunter, Corinne F. Maurice

**Affiliations:** Department of Microbiology & Immunology, Faculty of Medicine and Health Sciences, McGill University, Montreal, Quebec, Canada; Department of Public Health and Primary Care, School of Clinical Medicine, University of Cambridge, Cambridge, England; McGill Centre for Microbiome Research, Montreal, Quebec, Canada

## Abstract

The bacteria living in the human gut are essential for host health. Though the composition and metabolism of these bacteria is well described in both healthy hosts and those with intestinal disease, less is known about the activity of the gut bacteria prior to, and during, disease development – especially regarding gut bacterial replication. Here, we use a recently developed single-cell technique alongside existing metagenomics-based tools to identify, track, and quantify the replicating gut bacteria and their replication dynamics in the dextran sodium sulfate mouse model of colitis. We show that the proportion of replicating gut bacteria decreases when mice have the highest levels of inflammation and returns to baseline levels as mice begin recovering. We additionally report significant alterations in the composition of the total replicating gut bacterial community during colitis development. On the taxa level, we observe significant changes in the abundance of taxa such as the mucus-degrading *Akkermansia muciniphila* and the poorly described *Erysipelatoclostridium* genus. We further demonstrate that many taxa exhibit variable replication rates during colitis, including *A. muciniphila*. Lastly, we show that colitis development is positively correlated with increases in the presence and abundance of bacteria predicted to be fast replicators, suggesting that taxa with the potential to replicate quickly may have an advantage during intestinal inflammation. These data support the need for additional research using activity-based approaches to further characterize the gut bacterial response to intestinal inflammation and its consequences for both the host and the gut microbial community at large.

**Importance:** It is well known that the bacteria living inside the gut are important for human health. Indeed, the type of bacteria which are present and their metabolism is different in healthy people versus those with intestinal disease. However, less is known about how these gut bacteria are replicating, especially as someone begins to develop intestinal disease. This is especially important as it is thought that the active gut bacteria may be more relevant to health. Here, we begin addressing this gap by using several complementary approaches to characterize the replicating gut bacteria in a mouse model of intestinal inflammation. We reveal which gut bacteria are replicating, and how quickly, as mice develop and recover from inflammation. This work can serve as a model for future research to identify how the active gut bacteria may be impacting health, or why these particular bacteria tend to thrive during intestinal inflammation.

## Introduction

Inflammatory bowel disease (IBD) is an umbrella term for several chronic inflammatory intestinal diseases, including Crohn’s disease (CD) and ulcerative colitis (UC).^1^ The incidence of IBD has been increasing in both high-income and low- and middle-income countries over the past 10 years and shows no signs of slowing down.^2^ Though the exact etiology of IBD is unknown, evidence suggests that IBD results from a combination of genetic defects in the immune system and environmental triggers, which results in an aberrant host immune response to the gut microbiota.^3,4^ Indeed, this disease cannot occur without the presence of the gut bacteria.^5^ As such, these bacteria may play a crucial role in the manifestation and/or perpetuation of IBD.

Several studies have characterized the composition of the colonic or fecal bacteria in patients with IBD versus those without. Patients with IBD generally exhibit decreased bacterial diversity, increased abundances of aerotolerant and facultative anaerobic bacteria (such as members of the *Enterobacteriaceae* family), and decreased abundances of beneficial obligate anaerobes.^6^ It remains unclear what roles the gut bacteria play in IBD, and whether the observed diversity changes are a cause or consequence of disease. One emerging hypothesis is that the immune dysregulation apparent in IBD patients could initially alter the gut environment, favouring the survival and growth of bacterial taxa with pro-inflammatory properties, which further contribute to inflammation.^1,7,8^ This self-perpetuating cycle is broken primarily through the use of medication, such as corticosteroid drugs and 5-aminosalicylic acid, also known as mesalamine.^7,9^

Studies which have evaluated changes in gut bacterial transcripts, proteins, and metabolites have shown more consistent alterations than changes in gut bacterial composition in IBD patients, supporting the notion that gut bacterial metabolism plays a larger role in perpetuating inflammation than changes in community structure.^3,10^ For example, in IBD there is an increased abundance of bacterial genes encoding for, and proteins involved in, oxidative stress and DNA replication;^11,12^ increased levels of bacterial proteins related to sulfur metabolism and hydrogen sulfide production;^12^ increased transcriptional activity of specific bacteria, including *Clostridia* species and the mucin-degrader *Ruminococcus gnavus;*^6^ and significant alterations in bacterial metabolites, such as decreased levels of anti-inflammatory short-chain fatty acids like butyrate, and increased levels of primary bile acids.^6^

One fundamental aspect of bacterial activity which remains understudied in the gut is bacterial replication,^13^ specifically in the context of inflammation. Pioneering work from Olm et al. (2019) showed that bacterial replication rates were one of the best predictors of the development of necrotizing enterocolitis (NEC), an inflammatory intestinal disease in pre-term infants. Indeed, the authors reported a significant increase in gut bacterial replication rates, especially members of the *Enterobacteriaceae* family, preceding the development of NEC.^14^ In the same year, Riglar et al. (2019) tested their newly developed a synthetic gene oscillator, the repressilator 2.0, which reported on the growth dynamics of a transformed *E. coli* strain in a mouse model of dextran sodium sulfate (DSS)-induced colitis. The authors found that during active inflammation, there was increased variability in the growth rates of the transformed *E. coli*.^15^

In a more recent publication, Joseph et al. (2022) used an updated version of their replication rate measurement tool, termed Co-PTR, to identify taxa with altered replication rates in healthy individuals versus those with IBD.^16^ They found that gut bacterial replication rates exhibited large interindividual variability but seemed relatively stable within healthy individuals. They also reported that replication rates of specific taxa were associated with CD (*Subdoligranulum* species) or UC (*Roseburia intestinalis*, *Ruminiclostridium* species, and *Subdoligranum* species), whereas the relative abundances of these same taxa were not associated with their respective disease. Notably, they highlight that at least one other study also reported *R. intestinalis* as having increased replication rates in individuals with CD and UC, emphasizing the relevance of this taxon in IBD.

Collectively, these studies demonstrate how gut bacterial replication provides information about gut bacterial activity that complements relative abundance measurements, contributing to a better understanding of which taxa will thrive during intestinal inflammation. This could allow us to characterize how gut bacteria contribute to inflammatory conditions and shape the composition and function of the post-inflammatory gut microbiota.

Commonly used tools to study bacterial replication in community contexts rely on whole genome shotgun sequencing (WGS) data. Though these tools are informative and quantitative, they can be prohibitively expensive to implement and are data-analysis intensive.^17^ As such, there is a need for more physiologically-based techniques,^18^ which are more affordable and easily implementable, to complement existing WGS-focused techniques.

Most studies to date which evaluated gut bacterial activity or replication in IBD and other inflammatory conditions were cross-sectional and did not focus on disease development and progression. This is partly because it is challenging to identify and track individuals who will later develop IBD, despite new efforts in this direction.^19–21^ We hypothesize that tracking gut bacterial replication dynamics during the development and recovery from colitis will better inform us about the bacteria most relevant to intestinal inflammation.

Here, we use a combination of bacterial physiology- and bioinformatics-based techniques to capture changes in the composition and replication dynamics of gut bacteria during the development and recovery from chemically induced colitis in mice. The dextran sodium sulfate (DSS) colitis model is one of the most widely used rodent models of intestinal inflammation, reproducibly recapitulating the colonic damage and mucosal recovery of UC,^22^ despite its limitations.^22–27^ Importantly, this model allows the precise timing of colitis onset and development. We thus tracked the proportion and identity of actively replicating gut bacterial cells, quantified their replication rates when possible, and estimated the replication potential of the entire bacterial community over time.

As mice develop colitis, there are decreases in the proportions of replicating gut bacteria, which revert to baseline values as the mice recover. We also report significant changes in the composition of the replicating community of gut bacteria, as well as significant changes in the abundance of specific taxa, such as the mucus-degrader *Akkermansia muciniphila* and the poorly described *Erysipelatoclostridium* genus. Whole genome shotgun sequencing approaches reveal dynamic replication rates for many taxa throughout colitis development, including *A. muciniphila*. Lastly, we observed that the development of colitis coincided with the preferential retention in the gut of bacterial genomes with faster predicted doubling times. Our data support the need for additional functional approaches to better understand the gut bacterial dynamics underlying the progression of intestinal inflammation and the resulting altered microbial community.

## Results

### DSS reproducibly causes colitis and changes in fecal bacterial composition

We first confirmed that administration of 2% DSS for 5 days in the drinking water of the mice in our facility led to reproducible colitis. To do so, we monitored weight loss, blood in stool, and levels of fecal lipocalin-2 (LCN-2), a commonly used and sensitive marker of intestinal inflammation,^23^ before, during, and after administration of DSS (**Figure 1A; Supplementary Figure 1**). We incorporated these three parameters into a modified scoring metric of colitis as described by Kim et al. (2012) based on the disease activity index (DAI) (**Supplementary Table 1, Supplementary Table 2**).^28^ In this way, we defined the different health states of the mice during colitis development: baseline, pre-symptomatic, symptomatic, and recovery,^29^ as detailed in the Methods.

**Figure 1.**
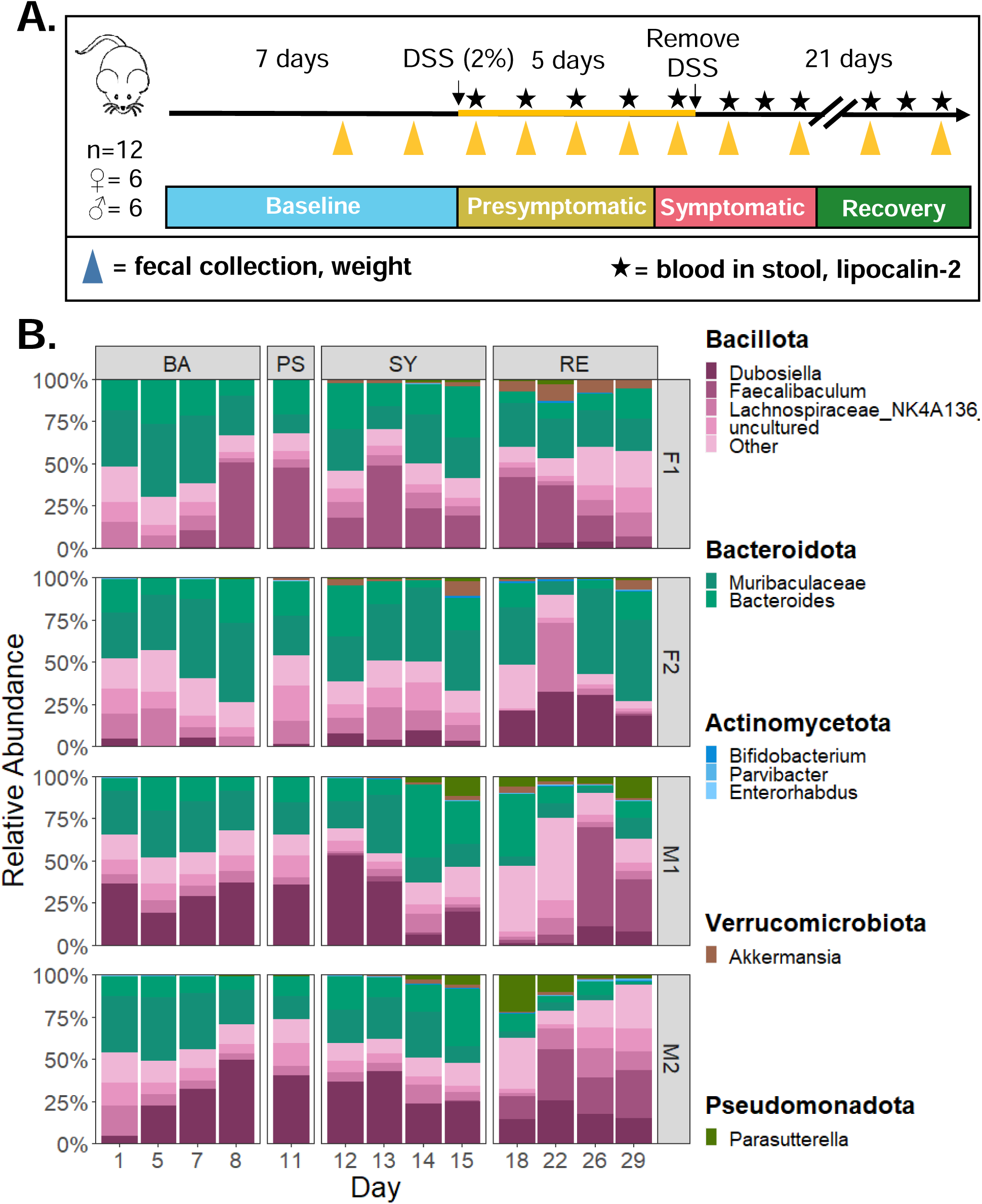
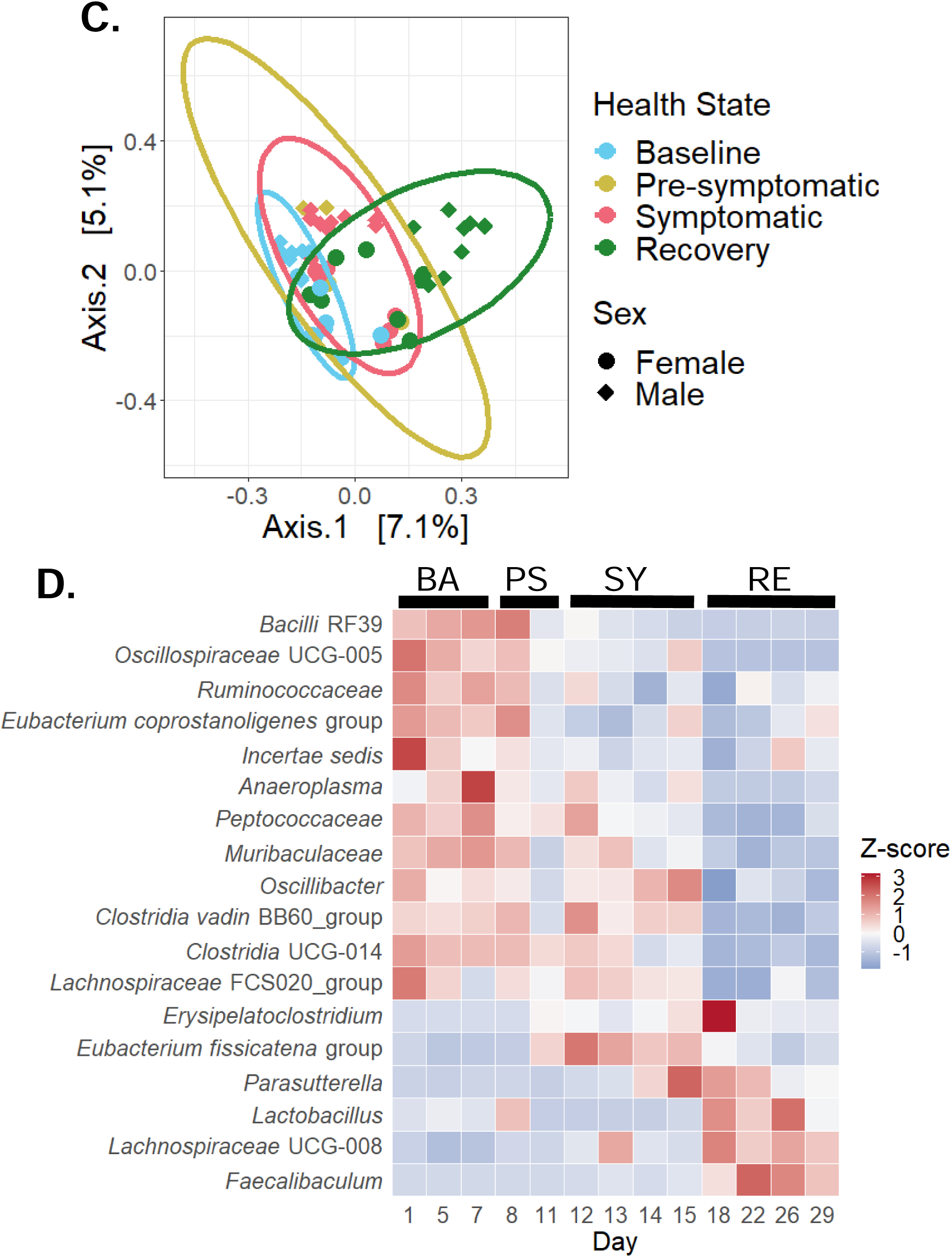
Experimental timeline and changes in gut bacterial community during DSS-colitis. (A) Timeline for each mouse experiment. After a 2-week acclimatization period, the feces of four cages of mice (two cages per sex, with 3 mice per cage) were collected over 7 days. Then, dextran sodium sulfate (DSS) was administered in the drinking water of the mice at a concentration of 2% (w/v) for 5 days, after which normal water was returned. Mouse feces continued to be collected throughout. Once DSS administration began, and thereafter, mouse health was monitored via measurements of blood in stool and lipocalin-2. Based on mouse health parameters, the phases of this study were split into four health states: baseline, pre-symptomatic, symptomatic, and recovery. (B) Taxonomic bar plots at the genus level of the fecal gut bacteria from each mouse during each health state (C) Beta diversity (Jaccard index) of the fecal gut bacterial communities from all cages, grouped per health state. D) Heat maps of standardized relative abundances of taxa from all cages of mice which were differentially abundant as compared to the baseline state. BA = Baseline; PS = Pre-symptomatic; SY = Symptomatic; RE = Recovery.

Amplicon sequencing of the 16S V4-5 region was conducted on bacterial DNA extracted from mouse feces collected throughout the experiment, as described in the Methods. Following this, QIIME2 was used to taxonomically identify the bacteria present in each sample (**Figure 1B**).^30^ The bacterial communities of these mice included the genera *Muribaculaceae* (F1: 25.37% ± 9.36; F2: 34.20% ± 12.42; M1: 17.62% ± 9.30; M2: 17.93% ± 12.80) and *Bacteroides* (F1: 17.34% ± 7.50; F2: 14.08% ± 8.37; M1: 15.68% ± 11.89; M2: 12.04% ± 7.91) of the Bacteroidota phylum; along with *Dubosiella* (F1: 0.64% ± 1.11; F2: 10.15% ± 10.96; M1: 21.55% ± 15.63; M2: 26.41% ± 12.61), *Lachnospiraceae* NK4136 group (F1: 7.58% ± 3.88; F2: 11.72% ± 10.09; M1: 5.01% ± 2.92; M2: 8.05% ± 5.10), and *Faecalibaculum* (F1: 24.01% ± 18.33; F2: 6.66 * 10^-3 % ± 2.40 * 10^-2; M1: 7.85% ± 1.72; M2: 7.09% ± 1.17) of the Bacillota phylum (**Figure 1B**).

When combining the data from all cages, we report no significant differences in the alpha diversity between the different health states for any of the metrics we assessed (**Supplementary Figure 2A**). Nevertheless, we saw a trend towards significance for the Shannon index, supported by a significant Friedman’s test (p = 0.044) but non-significant post-hoc test (Wilcoxon signed-rank test; p = 0.375 for all comparisons) (**Supplementary Table 3**).

At the beta diversity level, we report significant differences in the bacterial communities between baseline and both the symptomatic and recovery states based on the Jaccard index (PERMANOVA, q = 0.039; **Figure 1C; Supplementary Table 4**). Despite no significant differences between the health states for the other calculated beta diversity metrics (**Supplementary Table 4**), we observed the same pattern of deviation from baseline during the symptomatic state and partial return to near-baseline values during recovery (**Supplementary Figure 2B-D**).

When grouping the bacterial diversity data by sex, significant differences were observed in the beta-diversity values between health states (**Supplementary Figure 3**). For female mice (cages F1 and F2), this included significant differences in the unweighted Unifrac distance between the baseline and symptomatic health states (PERMANOVA, q = 0.027), and between the recovery and both the pre-symptomatic (PERMANOVA, q = 0.042) and the symptomatic (PERMANOVA, q = 0.027) health states (**Supplementary Table 5iv**). There were no significant differences for the other beta diversity metrics (**Supplementary Table 5i-iii**). For male mice (cages M1 and M2), there were significant differences in the unweighted Unifrac distance (UU), Jaccard index (JC), and Bray-Curtis dissimilarity (BC) between the baseline and symptomatic (PERMANOVA, UU: q = 0.02 ; JC: q = 0.036; BC: q = 0.039) and recovery (PERMANOVA, UU: q = 0.018; JC: q = 0.012; BC: q = 0.033) health states, and between the recovery and pre-symptomatic (PERMANOVA, UU: q = 0.0315; JC: q = 0.036; BC: q = 0.039) and symptomatic (PERMANOVA, UU: q = 0.02; JC: q = 0.015; BC: q = 0.033) health states (**Supplementary Table 6i, ii, iv**). There were no significant differences for the weighted Unifrac distance (**Supplementary Table 6iii**). These results suggest possible sex-specific differences in gut bacterial communities during the development of colitis.

We next sought to determine whether there were any changes in the abundance of specific taxa during the progression of colitis, using ANCOM-BC2 to conduct differential abundance analysis.^31^ We compared the abundances of each taxon at a genus level between baseline and the other health states (see Methods). We identified 3, 9, and 17 taxa as differentially abundant in the pre-symptomatic, symptomatic, and recovery states, respectively (**Supplementary Figure 4**).

We noted a consistent increase in the abundance of *Erysipelatoclostridium* and a *Eubacterium fissicatena* group member in all health states as compared to the baseline, as well as the increased abundance of *Parasutterella* and *Faecalibaculum* during the symptomatic and recovery states (**Figure 1D; Supplementary Figure 4**). A *Lactobacillus* member consistently appeared at significantly higher abundances during baseline relative to the pre-symptomatic and symptomatic health states; whereas UCG-005 (from the *Oscillospiraceace* family), a *Ruminococcaceae* family member, a *Eubacterium coprostanoligenes* group member, and RF39 (from the *Bacilli* class) all had higher relative abundances during baseline when compared to the symptomatic and recovery health states (**Figure 1D; Supplementary Figure 4**).

### Colitis alters the replicating and whole communities of gut bacteria

Next, we wanted to assess the proportions and identity of the replicating gut bacteria during the development of, and recovery from, DSS colitis. As such, a second independent mouse experiment with the same design was conducted, wherein the recently optimized EdU-click technique was used to label newly synthesized bacterial DNA.^13^ EdU-click was combined with flow cytometry, as well as fluorescence-activated cell sorting and 16S rRNA amplicon sequencing (FACSeq), to quantify the proportions of actively replicating cells in fresh mouse feces and to taxonomically identify these bacteria (see Methods and **Figure 2A**), respectively. The health status of the mice was monitored and health states were determined as described earlier (**Supplementary Figure 5**).

**Figure 2.**
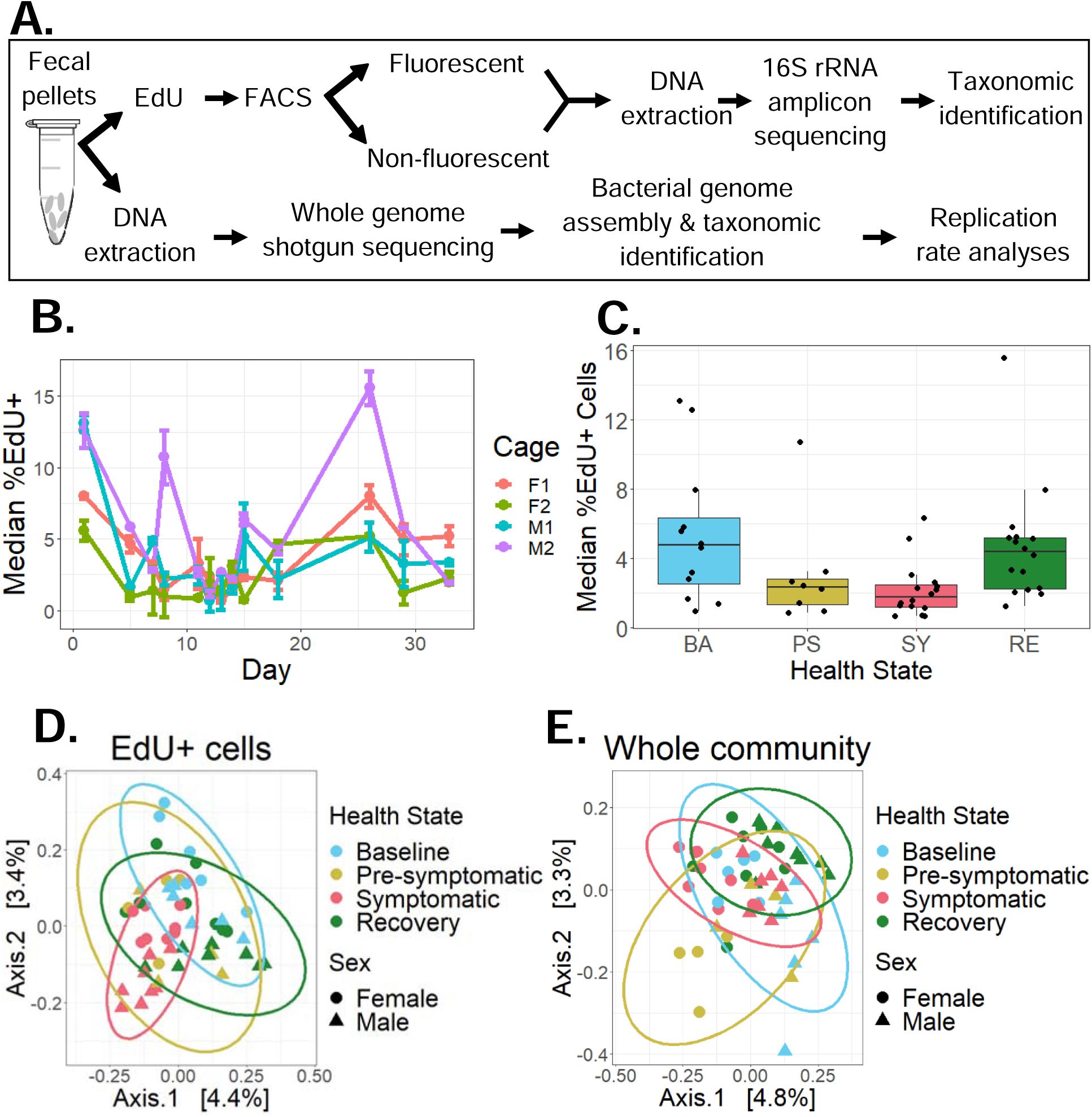
Experimental workflow and changes in the proportion and composition of replication gut bacteria during DSS colitis. (A) Workflow for each mouse experiment. Whenever fecal pellets were collected, half were immediately used for EdU-click and FACSeq; the other half were frozen for later DNA extraction, whole genome shotgun sequencing, and subsequent analyses. (B-C) Median proportion of replicating (%EdU^+^) cells, represented as a line graph showing per-cage dynamics over time (B) and as boxplots, grouping all cages per health state (C). (D-E) Beta diversity (Jaccard indices) of the replicating (EdU^+^) (D) and whole community (Whole) (E) of bacterial cells for all cages, grouped by health state

The proportion of replicating fecal bacterial cells, denoted as %EdU^+^ cells, changed as the mice developed and recovered from DSS-induced colitis (**Figure 2B**). Specifically, there was a notable but non-significant decrease in the median proportion of replicating cells from 4.75% ± 3.73 during baseline to 2.33% ± 1.35 during the pre-symptomatic state and 1.79% ± 0.94 during the symptomatic state, with a return to near-baseline values during the recovery period (4.40% ± 1.89) (**Figure 2C**). Notably, the variability in the median proportion of replicating cells between cages decreased during the pre-symptomatic (1.35) and symptomatic (0.94) time periods as compared to the baseline (3.73) and recovery periods (1.89) (**Supplementary Figure 6A,B**), demonstrating that the proportion of replicating cells was most similar between cages during active colitis.

We next wanted to determine if certain taxa preferentially replicate as mice develop colitis and recover. To accomplish this, we conducted 16S rRNA amplicon sequencing on the V4-5 region from DNA extracted from the replicating cells (EdU^+^) and the whole community (Whole) of bacteria originating from the same fecal sample (**Figure 2A**).

When considering the alpha diversity metrics, there were no significant differences in any metric between the different health states for either the EdU^+^ fraction or the Whole community (**Supplementary Figure 7; Supplementary Table 8**). The lack of statistically significant differences is likely due to both inter-cage variability and low numbers of cages (n = 4 cages).

We then compared the fecal bacterial communities of these mice across the different health states using beta diversity metrics, for both the EdU^+^ and Whole community of sorted bacteria. We saw that there was grouping and separation between the different health states in both fractions (**Figure 2D, E**). However, these differences were only statistically significant with the Jaccard index for the EdU^+^ fraction, between the symptomatic state and both the baseline (PERMANOVA, q = 0.012) and the pre-symptomatic (PERMANOVA, q = 0.012) health states (**Supplementary Table 9i**). As with the alpha diversity measurements, the lack of other statistically significant differences (**Supplementary Figure 8; Supplementary Table 9ii-iv**) is likely due to both inter-cage variability as well as low numbers of cages (n = 4 cages). When the two sexes were considered separately, only the Whole community of the male mice (cages M1 and M2) showed a statistical difference in any alpha diversity metric, being the weighted Unifrac distance, between the pre-symptomatic and recovery states (PERMANOVA, q = 0.042) (**Supplementary Figure 9P; Supplementary Table 14iii**). No other beta diversity metrics were significant (**Supplementary Figure 9A-O; Supplementary Table 11, Supplementary 12, Supplementary 13, Supplementary 14i, ii, iv**)

We then sought to identify taxa which were differentially abundant in either the EdU^+^ or the Whole fraction in the different health states relative to the baseline. As before, we first taxonomically identified the bacteria present in our samples using QIIME2 (**Figure 3A, B**).

**Figure 3.**
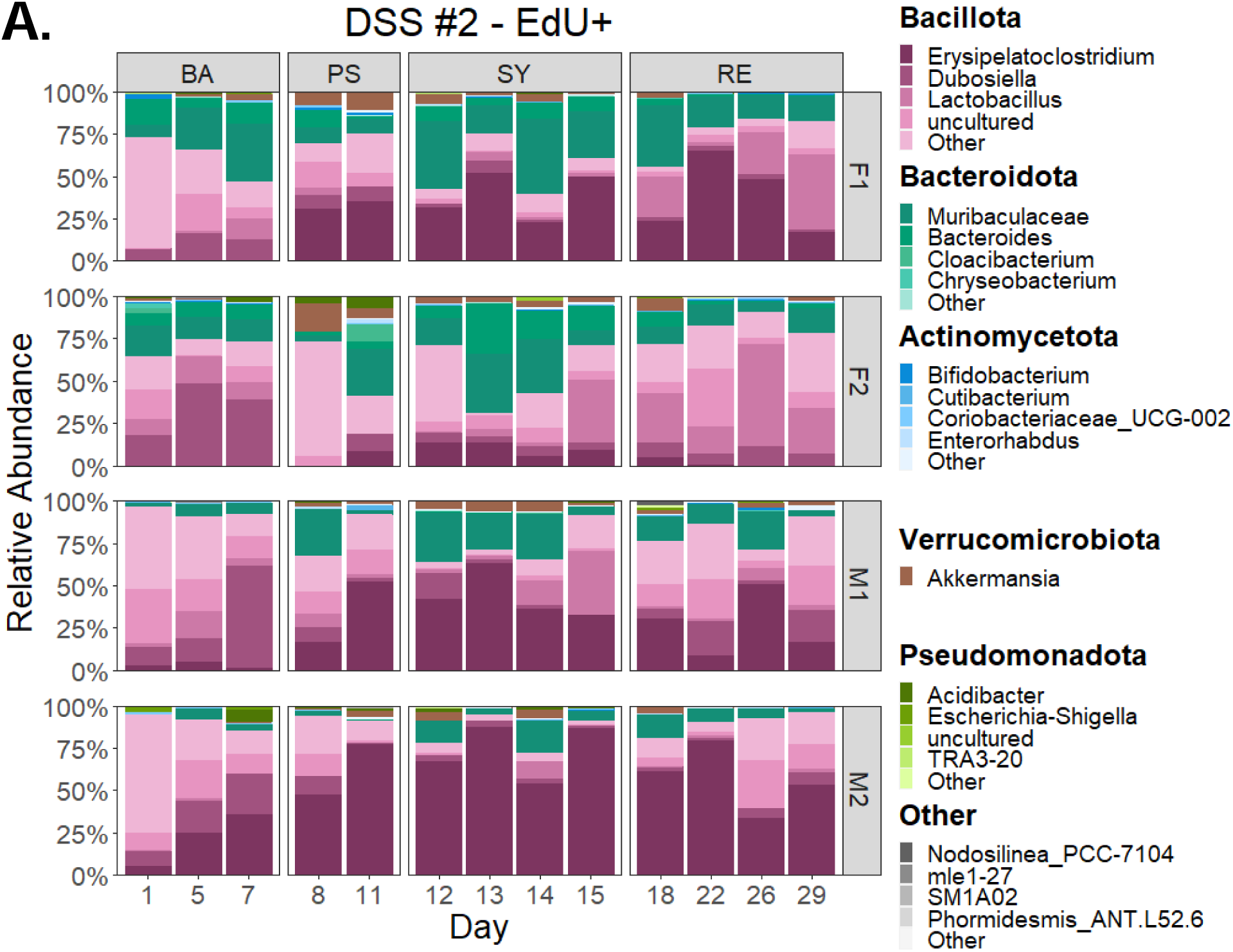
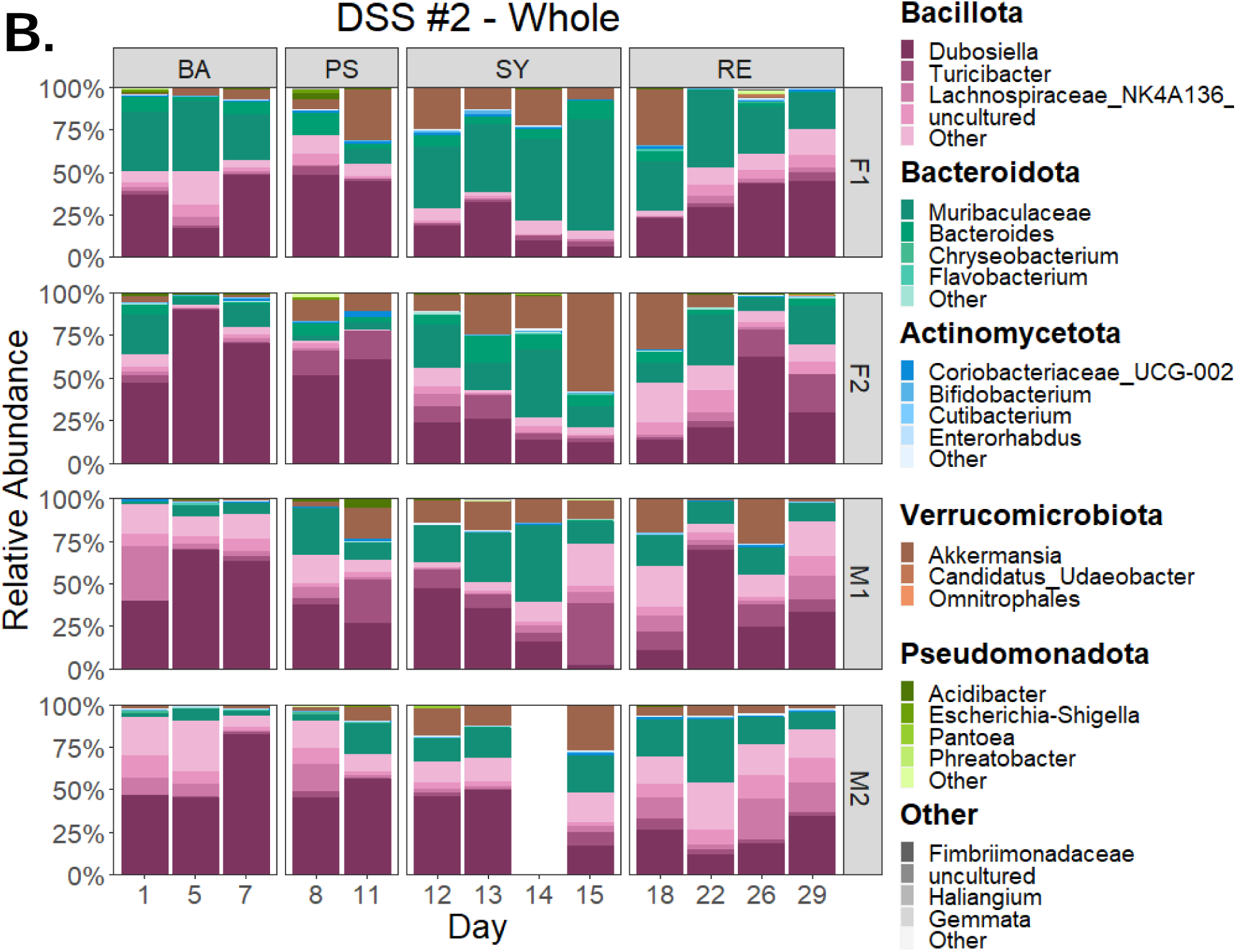
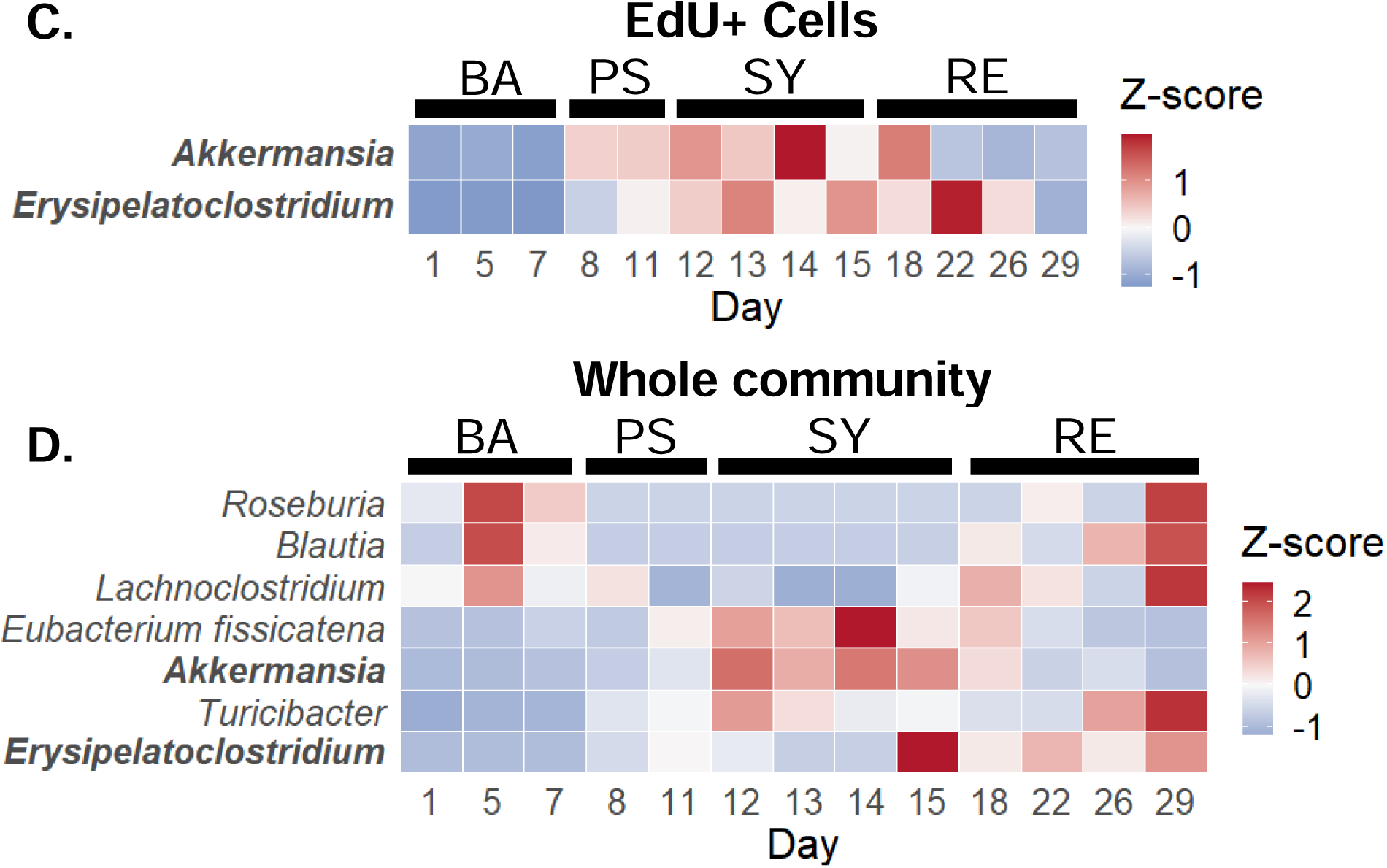
Taxonomy and differentially abundant taxa during DSS colitis. (A-B) Taxonomic bar plots of the fecal gut bacteria for each cage during each health state for the replicating (EdU^+^) (A) and whole community (Whole) (B) of bacterial cells. (C-D) Heat maps of the standardized abundance of taxa determined as differentially abundant as compared to the baseline state for the replicating (EdU^+^) cells (C) and the whole community (Whole) (D). BA = baseline; PS = pre-symptomatic; SY = symptomatic; RE = recovery.

In the EdU^+^ fraction, the most abundant genera included *Erysipelatoclostridium* (mean 28.76% ± 26.65), *Muribaculaceae* (mean 15.12% ± 11.19), *Lactobacillus* (mean 8.33% ± 13.05), *Lachnospiraceae* NK4A136 group (mean 6.38% ± 11.73), and *Dubosiella* (mean 9.23% ± 11.53) (**Figure 3A, B**).

In the Whole fraction, the most abundant genera included *Muribaculaceae* (mean 20.22% ± 14.59), *Dubosiella* (mean 36.09% ± 20.71), *Akkermansia* (mean 10.69% ± 11.58), *Turicibacter* (mean 5.50 % ± 7.40), and *Lachnospiraceae* NK4A136 group (4.39% ± 6.38) (**Figure 3A, B**).

We then used ANCOM-BC2 to conduct differential abundance analysis at the genus level. In the replicating (EdU^+^) fraction, *Akkermansia* significantly increased its abundance during the pre-symptomatic and symptomatic time points compared to the baseline state, whereas in the Whole fraction, this taxon only increased during the symptomatic state (**Supplementary Figure 10**). In contrast, *Erysipelatoclostridium* was significantly increased in both the EdU^+^ fraction and the Whole fraction in all health states compared to the baseline state (**Supplementary Figure 10**). In the Whole fraction during the symptomatic state, there were also transient increases in a *Eubacterium fissicatena* group member and *Turicibacter*, and decreases in *Lachnoclostridum*, *Blautia*, and *Roseburia*, with no corresponding changes in the EdU^+^ fraction (**Supplementary Figure 10A-D**).

The changes in the proportion of replicating cells (EdU^+^) of both *Akkermansia* and *Erysipelatoclostridium* species mirrored their changes in total abundance (Whole), suggesting the active and successful replication of these taxa, leading to increased total biomass (**Figure 3C, D; Supplementary 10G**). This increase was transient for *Akkermansia* but seemed to persist for the *Erysipelatoclostridium* taxon.

### Altered replication dynamics of fecal bacteria during colitis development

As a complementary method to EdU-click, we also used whole genome shotgun (WGS) sequencing to quantify the replication dynamics of fecal bacteria on a subset of samples. Bacterial genomes were assembled from WGS data into metagenome-assembled genomes (MAGs) as described in the Methods. WGS sequencing was performed on fecal samples from mice in cages F2 and M2 (see **Supplementary Table 15** for WGS statistics), as these mice had more severe signs of inflammation compared to their biological replicate cages (**Supplementary Figure 5**).

The taxonomic composition and differentially abundant taxa of the MAGs, as well as their beta diversity, are summarized in **Figure 4** and **Supplementary Figure 11**, respectively. None of the beta diversity measurements showed significant differences in the fecal bacterial composition (**Supplementary Figure 11**), likely due to the limited number of genomes for which taxonomy could be determined (37 MAGs for cage F2; 43 MAGs for cage M2). Nevertheless, differential abundance analysis showed a significant increase in the relative abundance of an *Akkermansia muciniphila* MAG, and a significant decrease in the relative abundance of two MAGs from the *Lachnospiraceae* family, a 14-2 species and a COE1 species, during all health states as compared to the baseline state (**Figure 4C-E**). There was also a significant increase in the relative abundance of a *Duncaniella* MAG during the symptomatic state (**Figure 4D**).

**Figure 4.**
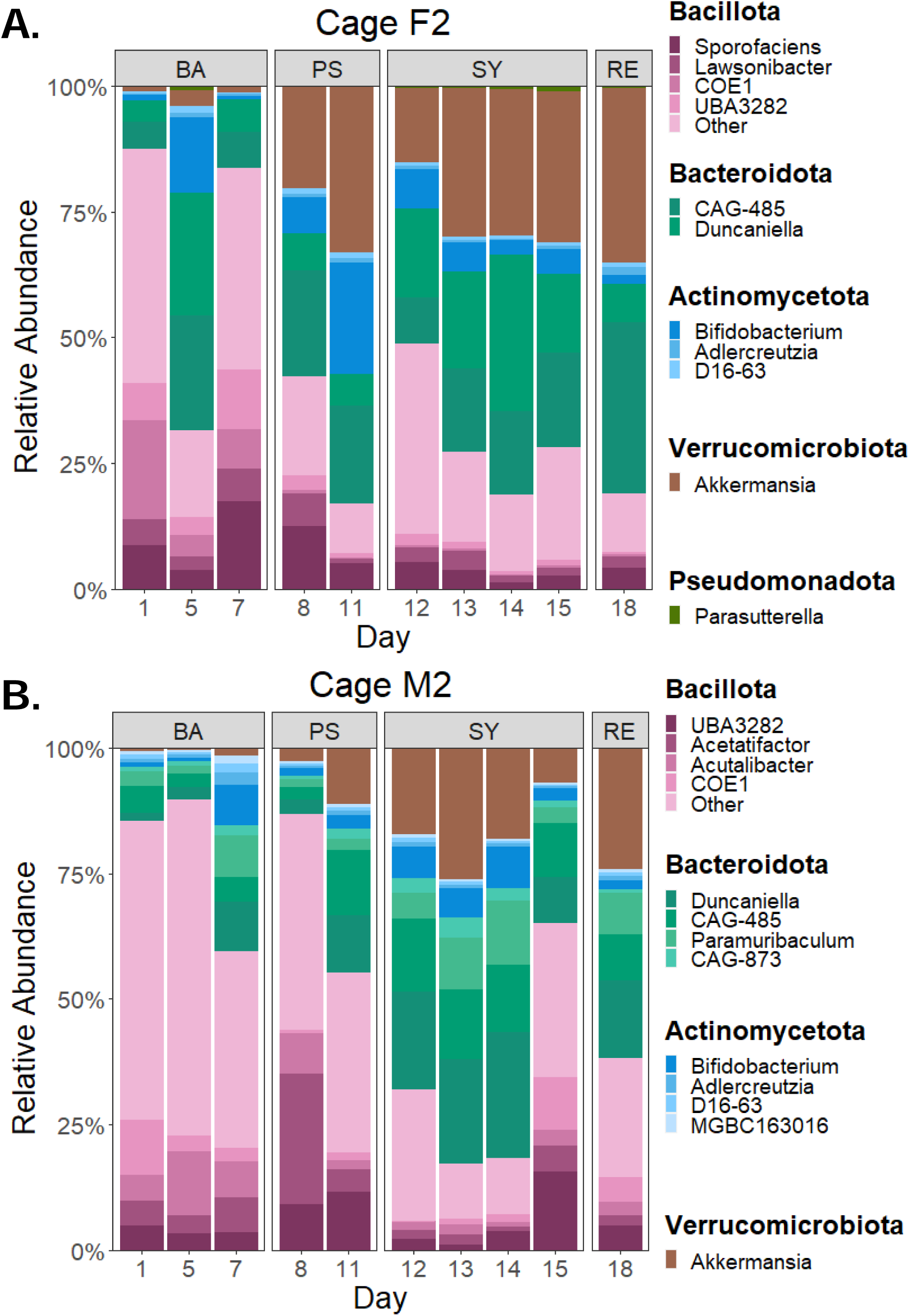
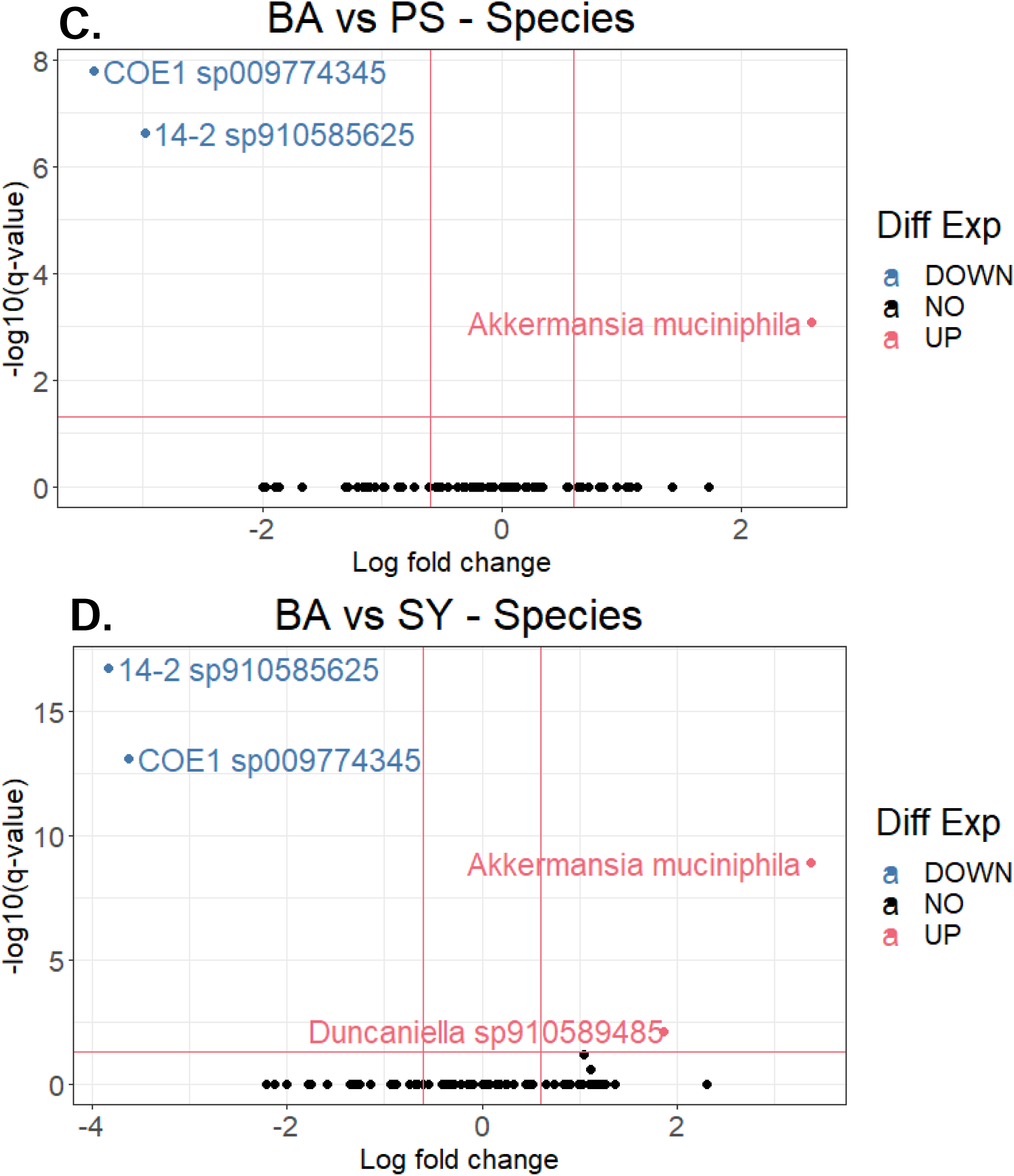
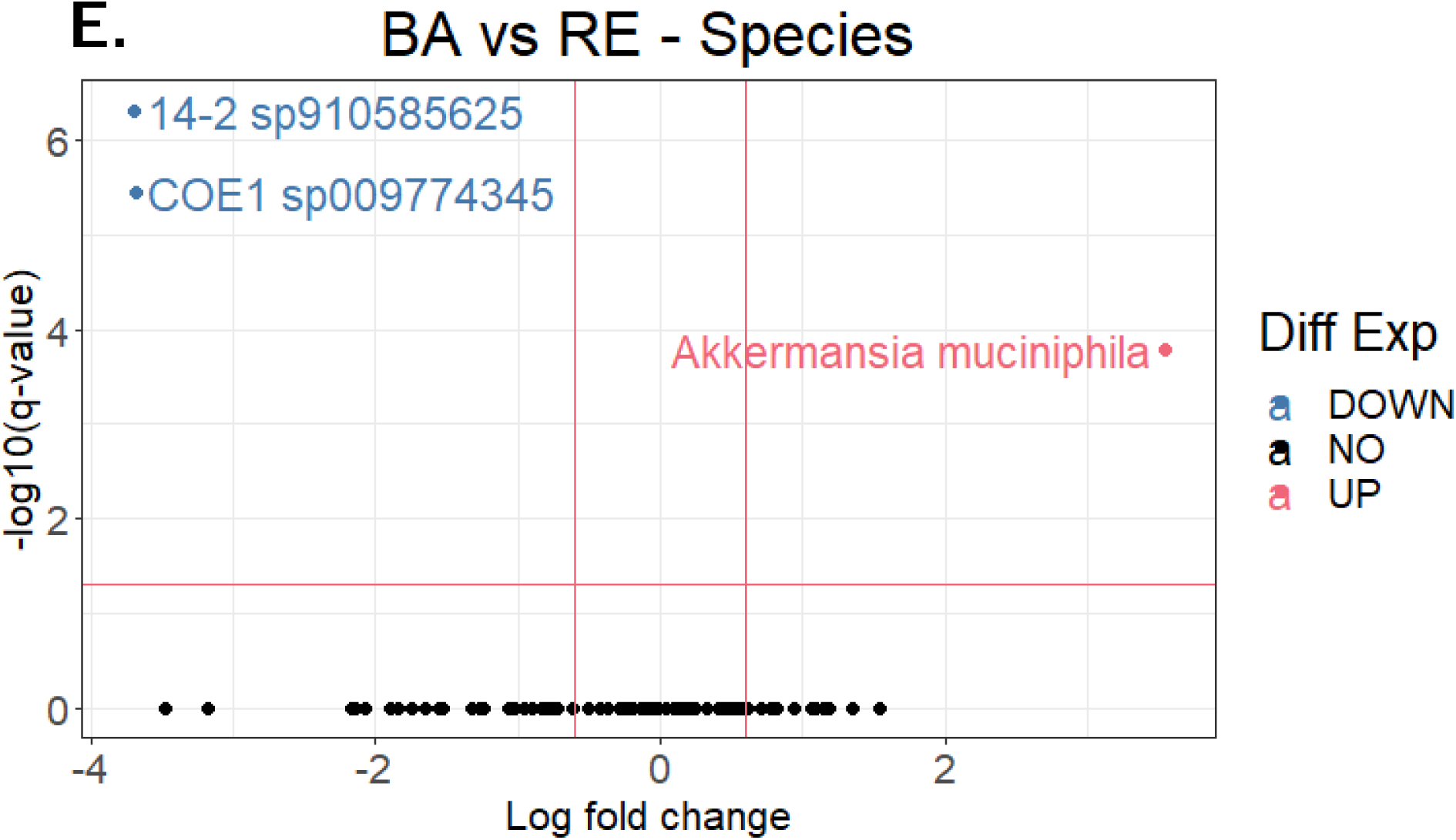
Relative abundance and differentially abundant taxa from whole genome shotgun sequencing (WGS) data. (A-B) Taxonomic bar plots at the genus level for the metagenome-assembled genomes. (C) Volcano plots of metagenome-assembled genomes determined to be differentially abundant from the baseline. Note that genomes 14-2 & COE1 are both from the Lachnospiraceae family. BA = baseline; PS = pre-symptomatic; SY = symptomatic; RE = recovery.

Next, we wanted to quantify the replication dynamics of these MAGs using two distinct methods: the peak-to-trough (PTR) method, which estimates instantaneous replication rates,^32^ and the codon usage bias (CUB) method, which estimates the maximal replication potential or minimal doubling time (DT).^33–35^

To calculate the replication rates during the development of DSS colitis for as many quality genomes as possible, we used the PTR-based tool GRiD^36^, which works well with medium quality MAGs at low coverage (**Supplementary Table 15**). GRiD is specialized in using genomes with a coverage of as little as 0.2x and as many as 90 scaffolds/Mbp, and for genomes with at least 1x coverage, as little as 50% completion and as much as 200 scaffolds/Mbp.^36^ The other main PTR-based tools, iRep,^37^ DEMIC,^38^ and CoPTR,^16^ all have requirements or assumptions which are not appropriate for the quality, coverage, or temporal nature of our data. The differences between these tools have been thoroughly evaluated elsewhere and will not be covered here.^16,35,39^

From cage F2, we were able to calculate PTR values using GRiD across all time points for 32 MAGs (corresponding to 53% of all MAGs from cage F2) (**Supplementary Table 15, Supplementary Table 16**); from cage M2, we calculated PTR values across all time points for 43 MAGs (corresponding to 65% of all MAGs from cage M2) (**Supplementary Table 15, Supplementary Table 17**). Of these MAGs, we highlighted the replication rate dynamics of the following four: *Akkermansia muciniphila* (**Figure 5A**), a *Duncaniella* species (**Figure 5B**), *Bifidobacterium globosum* (**Figure 5C**), and a *Muribaculaceae* species (**Figure 5D**). We selected these taxa based on whether they met all or most of the following criteria: they were differentially abundant between the health states, had the highest median replication rates during the symptomatic period, had replication rate measurements for all experimental days, were one of the most abundant taxa overall or during the symptomatic period, and had replication rate measurements estimated in both mouse cages.

**Figure 5.**
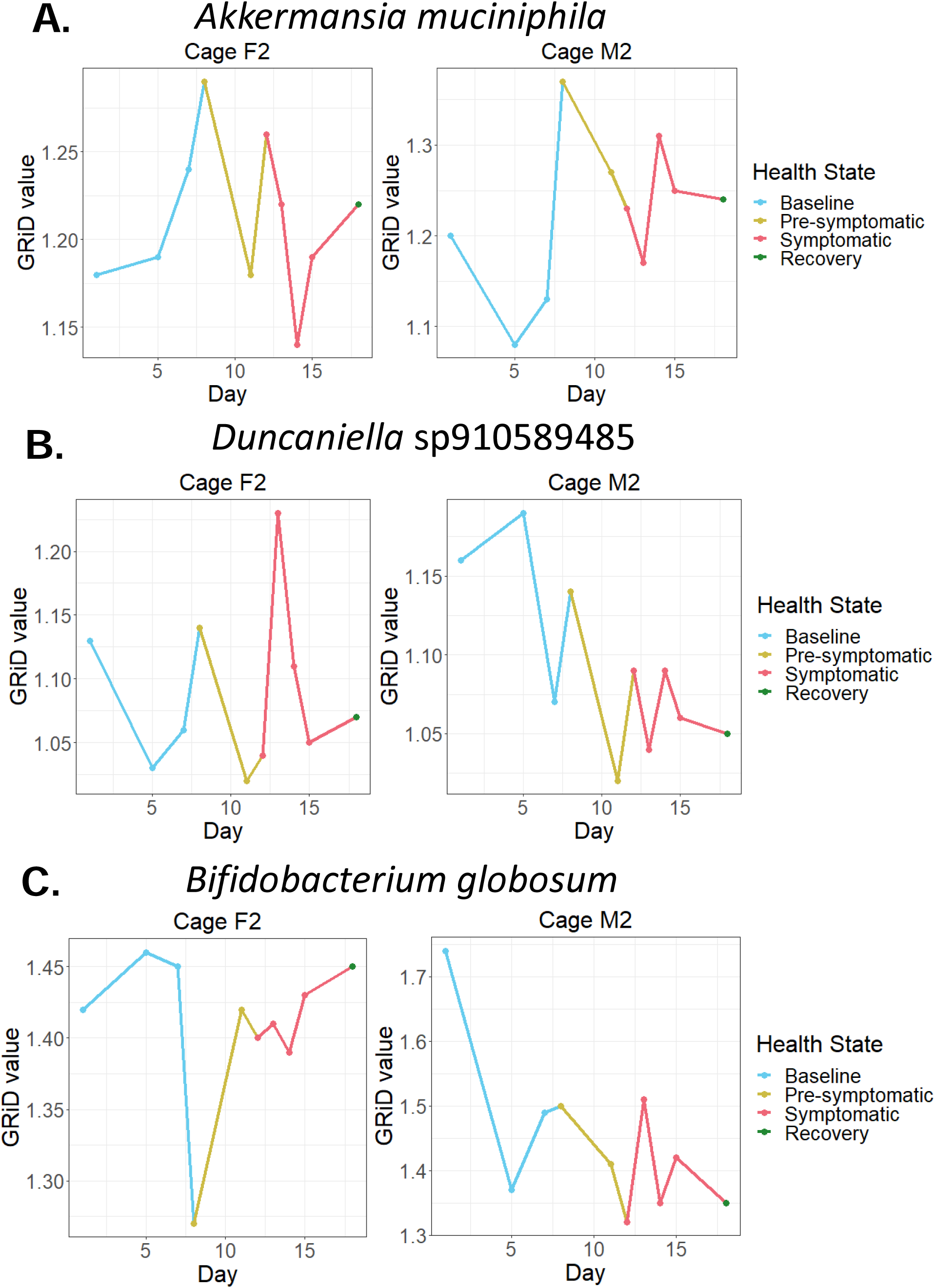
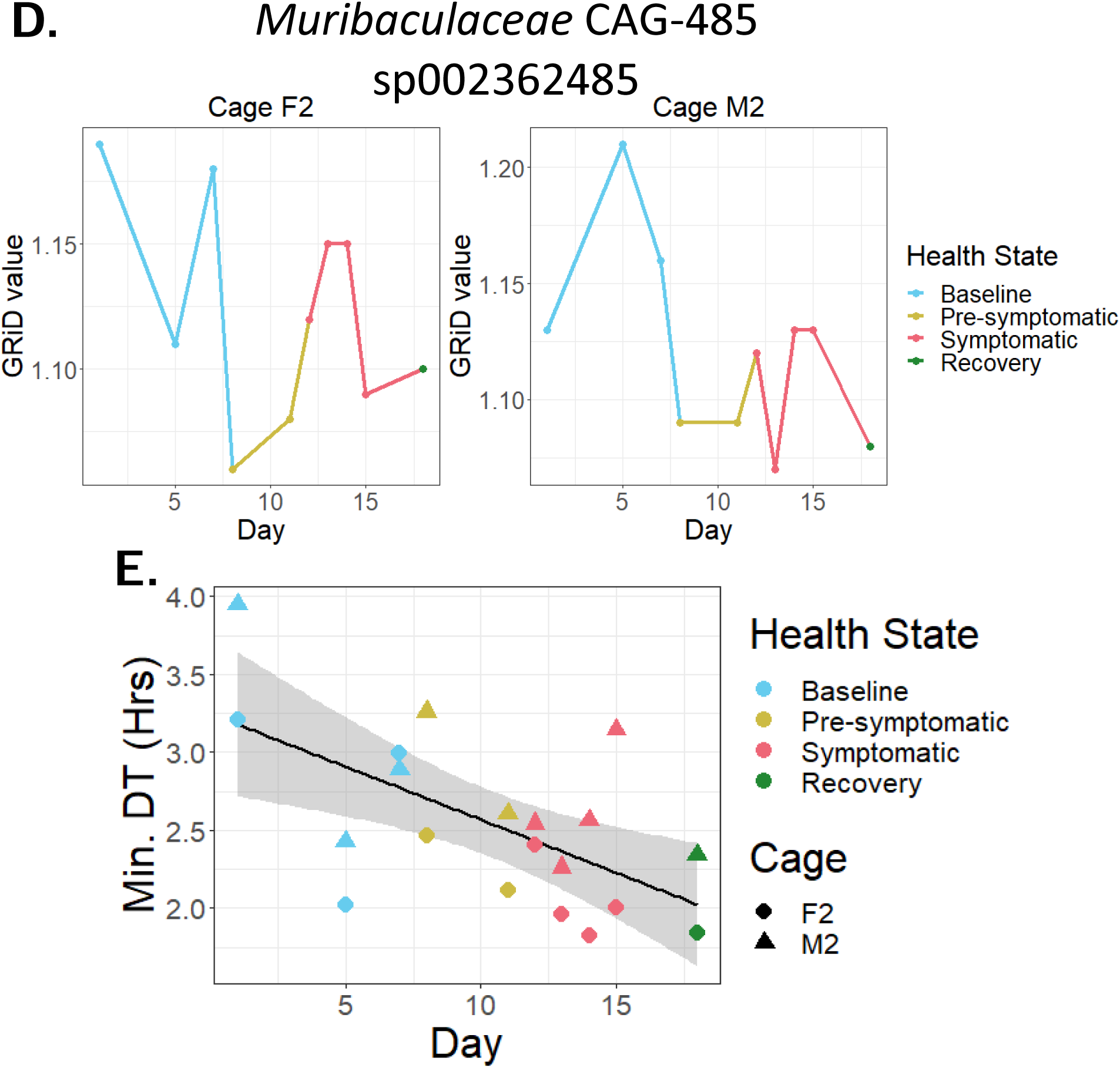
Replication dynamics of gut bacteria during DSS colitis. (A-D) Replication rates of representative taxa as calculated by GRiD, coloured by health state. (E) Correlation between health states and minimal doubling times (DTs) for both cages, as calculated by gRodon.

These taxa appear to have cage-specific dynamics. For instance, in cage F2, the *A. muciniphila* MAG had no correlation with the health status of the mice; whereas, in cage M2, the *A. muciniphila* MAG had consistently higher replication rates once the mice developed colitis relative to the baseline state (**Figure 5A**).

For the *Duncaniella* species, this MAG had its highest peak of replication rates during the symptomatic period in cage F2; while the same MAG had the highest replication rates during the baseline period relative to other periods in cage M2 (**Figure 5B**).

For *B. globosum*, this MAG had relatively steady replication rates in cage F2, except for a rapid and transient decrease on the first day of the pre-symptomatic period. This same MAG had decreased replication rates throughout the experiment starting after the first baseline day in cage M2 (**Figure 5C**).

Finally, for the *Muribaculaceae* species, this MAG had its lowest replication rates during the pre-symptomatic state in cage F2, whereas in cage M2 its replication rates were highest during the baseline state and were consistently lower afterwards (**Figure 5D**).

We further determined no correlation between bacterial replication rates and the relative abundance of their corresponding taxa in the whole community (Pearson r = -0.14756; r^2 = 0.02177; **Supplementary Figure 12**).^16^

To glean more information about the replication abilities of the taxa we identified, we next used gRodon, a CUB-based tool, to calculate the minimal doubling times (DTs) for each MAG (**Supplementary Figure 13**).^34^ This method provides information on the replication potential of bacteria, as opposed to their instantaneous replication rates. Previous studies have shown that measurements of replication potential using CUB-based tools can more accurately predict the replication behaviour of certain taxa, especially slower growing ones in complex communities.^39^

The median minimal DTs for all phyla in both cages was between 1.39 and 3.46 hours (**Supplementary Figure 13A; Supplementary Table 16, Supplementary Table 17**), supporting the notion that gut bacteria are fast replicators.^33^ Within the Bacillota phylum, there was a larger range (F2: 0.89-7.09 hrs; M2: 0.63-7.79 hrs) and more variability (F2: median absolute deviation (MAD) 1.67 hrs; M2: MAD 1.48 hrs) in the minimal DTs as compared to the other phyla with more than one genome present (F2: Actinomycetota range: 0.98-2.33 hrs, MAD: 0.74 hrs; Bacteroidota range: 1.44-2.59 hrs, MAD: 0.66 hrs; M2: Actinomycetota range: 1.04-2.72 hrs, MAD: 0.64 hrs; Bacteroidota range: 1.50-2.53 hrs, MAD: 0.55 hrs) (**Supplementary Table 18, Supplementary Table 19**). This range and variability specifically occurred within the Clostridia class (F2: range: 0.89 -7.09 hrs; MAD: 1.69 hrs; M2: range: 1.15-7.79 hrs; MAD: 1.40 hrs), which were comprised mostly of genomes from the *Lachnospiraceae* family (F2: 15 genomes: range: 1.53-7.00 hrs; MAD: 0.57 hrs; M2: 30 genomes: range: 1.64-7.79 hrs; MAD: 1.28 hrs) (**Supplementary Figure 13B,C; Supplementary Table 18, Supplementary Table 19**).

To see whether there were any changes in the replication potential of all bacteria detected over time, rather than individual taxa, we next used gRodon’s metagenome mode to calculate the minimal DTs of the whole community of gut bacteria during the development of colitis.^35^ There was a negative but non-significant (p > 0.05) slope when plotting the minimal DTs of both cages of mice as a function of the health state, while accounting for cage and day as random effects (see Methods) (**Figure 5E, Supplementary Figure 14; Supplementary Table 20**). However, this trend was close to significant (p = 0.0743) when the minimal DTs were plotted as a function of fecal LCN-2 levels, a sensitive marker of colitis (**Supplementary Figure 15; Supplementary Table 21**)^40^. Both data indicate that the colitic environment could select for bacteria with the capacity to replicate quickly.

As fast replication could be a strategy for bacterial taxa to persist in a perturbed environment,^41^ we also sought to describe the link between MAGs with the potential to replicate quickly (i.e., with low minimal DTs estimated by gRodon) and those with increased replication rates and/or increased relative abundances during the symptomatic period.

When combining information from all of the MAGs and both cages together, we report no correlation between the minimal DT of a MAG and its median replication rate during the symptomatic period (r = 0.049; **Supplementary Figure 16**). In contrast, there seemed to be a weak negative correlation between the minimal DT of a MAG and its median relative abundance during the symptomatic period (r = -0.329), suggesting that bacteria with the potential to replicate quickly tend to be at higher abundances during active colitis (**Supplementary Figure 17**). However, our analyses are limited by the rare high-abundance bacterial taxa and the prevalent low-abundance taxa.

## Discussion

This study sought to determine the replication dynamics of gut bacteria during the development of, and recovery from, intestinal inflammation in a mouse model of chemically induced colitis. We found that the development of colitis resulted in decreased proportions of replicating gut bacterial cells, as well as significant changes in the composition of the replicating gut bacterial community. At the genus level, we report significantly increased abundances of replicating *Akkermansia* and *Erysipelatoclostridium* taxa, which preceded increases in their relative abundances in the community. Replication rates of *A. muciniphilia, Bifidobacterium globosum,* a *Muribaculaceae* species, and a *Duncaniella* species also changed during colitis. Lastly, we saw that the progression of colitis coincided with an overall decrease in the predicted minimal doubling time of the whole bacterial community. Collectively, our data suggest that the composition of the mouse fecal bacterial community shifts towards taxa with the potential to replicate faster as intestinal inflammation progresses.

### DSS colitis results in broad-level and taxon-specific changes in gut bacterial communities

While the role of gut microbial communities in intestinal inflammation is increasingly studied, most studies remain cross-sectional. Thus, the dynamic alterations of the gut microbiota during colitis progression remain less characterized.^19,20,29,42–50^ By monitoring murine fecal microbial communities in DSS colitis, we aimed to determine some of these underlying microbial dynamics of colitis. While we do not report significant decreases in alpha-diversity, unlike other DSS studies^48,50–52^, our beta-diversity analyses suggest that these gut bacterial communities may have reached an alternative stable state distinct from baseline.^29,43,48,49,53^ These dynamics were especially marked in male mice, possibly explaining the more reproducible and severe phenotype typically observed for males in this colitis model.

Our longitudinal approach allowed us to determine that differentially abundant gut bacterial taxa could be categorized into 4 broad response groups, according to their dynamics during colitis progression (**Supplementary Figure 4**). Taxa in Group 1 – which include RF39, *Oscillibacter,* a *Ruminococcaceae* member, and a *Clostridium vadin* BB60 group member – had decreased abundances during the recovery period, suggesting that they likely cannot survive a colitic environment. In contrast, Group 2 taxa, including *Erysipelatoclostridium* and a *Eubacterium fissicatena* group member, had increased abundances during colitis development and onwards, suggesting that these taxa expanded when the host environment was perturbed, and were able to maintain their increased abundances even as the host recovered. Other response groups include taxa which decreased temporarily below the limit of detection during colitis but which returned during the recovery period (Group 3; *Lactobacillus*); and those which only expanded during recovery, possibly taking over a niche made available by the loss of other taxa (Group 4; *Lachnospiraceace* UCG-008).

The limited existing literature on the above highlighted taxa suggests some possible explanations for their dynamics. For instance, RF39 of Group 1, which has been shown to decrease in abundance in mouse models of colitis,^54–65^ has also been described as having a reduced genome and as being metabolically dependent on other bacteria for some amino acids and vitamins.^66,67^ Our data support the hypothesis that auxotrophic bacteria such as RF39 are less likely to survive harsh colitic environments, as has been suggested to occur in IBD patients.^68,69^ In contrast, Group 2 bacterial taxa *Erysipelatoclostridium* and *E. fissicatena* persistently increase after colitis development, as reported elsewhere.^70–87^ While the mechanisms behind this increase are unclear, our data align with the literature while also contributing a finer time scale regarding their changes in abundance during colitis.

At least one other DSS colitis study has seen the growth patterns of Group 3 member *Lactobacillus* reported here.^88^ One putative explanation for such behaviour is that, since *Lactobacillus* species colonize the forestomach of mice, these taxa can be washed into the cecum and colon – especially during the diarrhea commonly seen in DSS colitis.^22–24,49,89^ If a favourable niche in these distal intestinal locations becomes available during a perturbation, *Lactobacillus* may be able to establish there; whereas under homeostatic conditions, this would be less likely to occur.^90^ Furthermore, similarly to the earlier discussed taxon RF39, the metabolic dependence of most *Lactobacillus* species likely precludes their ability to thrive in an inflamed environment.^90^ After the inflammation, however, it may be possible that *Lactobacillus* from the forestomach establish in the colon.

For the Group 4 member *Lachnospiraceae* UCG-008, little is known about its metabolism; or indeed, the metabolism of most *Lachnospiraceae* species,^91^ despite including many SCFA-producers and metabolizers of complex plant-derived carbohydrates.^92^ *Lachnospiraceae* UCG-008 is likely a butyrate producer, since the *Lachnospiraceae* family is known to contain many butyrate-producing members.^91–94^ There is discrepancy between different studies as to whether *Lachnospiraceae* members are increased or decreased in various intestinal perturbations, and whether this is beneficial or detrimental to the host.^91,94–99^ It has been suggested that the impact of *Lachnospiraceae* members on the host is likely species or strain specific.^91,94,96^ As such, the overall role of *Lachnospiraceae* UCG-008 in colitis remains unclear. One speculation as to how this taxon thrives during colitis recovery is that it could create endospores.^100^ Upon the return to a more favourable environment, such as when the mice begin recovering from colitis, these endospores may allow *Lachnospiraceae* UCG-008 to rapidly take advantage of newly opened niches in the gut.

### Decreased proportions of replicating gut bacteria during colitis

To determine if the aforementioned patterns of altered abundances reflected changes in gut bacterial replication, we quantified and taxonomically identified the replicating gut bacteria using EdU-click in an independent study. We observed a decreased proportion of replicating bacteria during the pre-symptomatic and symptomatic stages, suggesting that the inflammatory environment is detrimental for the replication of most taxa.

Importantly, we noted that the replicating gut bacterial community during the symptomatic state was markedly different from other health states. Narrowing down to a per-taxon basis, we report significant differences in the abundance of *Akkermansia* and *Erysipelatoclostridium* in both the replicating fraction and in the whole bacterial community.

Since *Akkermansia* began increasing its number of replicating cells during the pre-symptomatic state, and then subsequently increased its abundance in the whole fraction during the symptomatic state, this suggests that its increased number of replicating cells successfully led to an increase in its biomass. Once the mice began to recover, however, the abundance of this taxon quickly decreased back to baseline values.

The impact of *Akkermansia* in DSS colitis is unclear, as it is associated with health in humans and is depleted in individuals with IBD,^101^ whereas it is commonly seen to increase during DSS colitis in mice.^29,102–104^ In addition, while some studies have found that mice with higher levels of *Akkermansia* exhibit less severe colitis,^105^ supporting a protective role for this taxon, other studies have shown the reverse^.103^ More recent studies highlight that the impact of *Akkermansia* on host health could be strain specific^106^ or depend on the microbial and immunological status of the host.^107^ As a mucus degrader, *Akkermansia* could worsen inflammation,^108^ rendering the host intestinal epithelium more accessible to microbial translocation. Alternatively, its mucus metabolism could stimulate increased mucus secretion by the host,^109^ as some *Bacteroides* species have been shown to do.^110^ An emerging hypothesis for the transient increase in *Akkermansia* during DSS colitis is that a DSS-dependent increase in mucus permeability^89^ allows this taxon greater access to its preferred carbon source.^111^ *Akkermansia* is also likely to be somewhat resistant to the increased oxidative environment of colitis, given that there are higher levels of oxygen present at the mucus layer where this taxon resides.^112^ The combination of its resistance to oxidative stress and an increased access to its preferred energy source creates a perfect environment for *Akkermansia* to thrive, at least until host mucosal recovery begins.^113^

The *Erysipelatoclostridium* taxon was increased at all time points relative to the baseline state, in both the replicating and the whole bacterial community. Despite a limited number of studies noting the contrary^99,114–116^, these results are in accordance with the behaviour of *Erysipelatoclostridium* reported in prior studies during intestinal disturbance. Indeed, this genus and one of its species, namely *Clostridium ramosum* (also known as *E. ramosum* or *Thomasclavelia ramosum*),^117–119^ have been associated with intestinal inflammation or injury in humans,^46,48,73,120–127^ increasing in abundance in mouse models of colitis^71–82^ and infection^128^, and being positively correlated with fecal water content (i.e., diarrhea), a common colitis symptom^2,22–24,129,130^. The high replication activity of *Erysipelatoclostridium* we report demonstrates that it is a particularly active taxon during colitis, resulting in a possibly outsized impact on the microbial environment and the host.

What allows *Erysipelatoclostridium* to benefit from perturbations can only be speculated, since its metabolism is not well described,^70^ despite being a dominant gut Clostridial taxon.^131^ The best described member of this genus is *C. ramosum*, which is reported to produce low amounts of acetate and possibly butyrate, and can significantly increase the number and function of anti-inflammatory Treg cells.^132,133^ These capabilities point towards the potential anti-inflammatory role of this taxon, without explaining its continual increase in abundance in perturbed intestinal conditions.

As more thoroughly discussed by Beauchemin et al.,^13^ not all bacterial taxa can be labelled with EdU, and labeled bacteria likely do not equally incorporate EdU into their DNA. As such, this study can only report on replicating taxa amenable to EdU-click, and likely does not capture all replicating taxa. As one way to address this limitation, we also used metagenomics-based bioinformatic tools to calculate bacterial replication dynamics on a subset of our data, as discussed below.

### Taxon-specific changes in gut bacterial replication rates and increased abundances of fast replicating gut bacteria during colitis

Stool samples from ten non-consecutive sampling times between days 1-18 in one cage of female mice (F2) and one cage of male mice (M2) underwent shotgun sequencing. This allowed us to quantify the replication dynamics of several metagenome-assembled genomes (MAGs) during colitis development, using both PTR-based and CUB-based approaches. For the most part, the oscillatory replication rates for each taxon in each cage went back to near-baseline values, with no clear link to the host health states – with the exception of *A. muciniphila*. In agreement with our EdU-click and differential abundance analyses, the *A. muciniphila* genome seen in both cages replicated most rapidly once DSS was administered, after which its replication rates decreased back to near baseline values as colitis developed.

We also observed cage-specific replication rate dynamics of the same gut taxa. For instance, the *Duncaniella*, *Bifidobacterium globosum,* and *Muribaculaceae* CAG-485 genomes in cage M2 had decreased replication rates as colitis developed, whereas in cage F2 these same taxa had variable replication rates. These differences suggest individual-specific gut bacterial replication dynamics, as reported in previous human cohorts^16^, in addition to possible sex- and cage-specific responses to inflammation. Since *Duncaniella, Muribaculaceae* CAG-485, and *B. globosum* are not as well described as *A. muciniphila*, the possible explanations underlying their replication rate patterns remain to be determined.

We additionally report a lack of correlation between gut bacterial replication rates and their associated relative abundances, as noted by other studies^16,32,37,134^. Indeed, relative abundance measurements account for the culmination of bacterial growth and death, whereas replication rates only account for the former, emphasizing that both approaches measure distinct aspects of the gut bacterial community.^16,32,37^

We next used an independent metric based on genome-specific CUB bias, using gRodon, to determine possible links between gut bacterial replication and the response of specific taxa during inflammation. When calculating the replication potential of the whole gut bacterial community over time, we noted slightly lower minimal doubling times (i.e., increased maximal replication rates) as colitis developed or as levels of LCN-2 increased. This suggests that faster replicating gut bacteria may be preferentially selected for in an inflammatory environment.

These findings were further corroborated by the negative correlation between the minimal doubling time (DT) of a MAG and its median relative abundance during the symptomatic period, suggesting that taxa with the ability to replicate quickly are present at higher abundances during colitis. This is in line with previous research which suggests that faster replicating bacteria can adapt more quickly to changing environments,^33,135,136^ that gut bacteria during IBD flares may be replicating more than during remission,^137^ and that fast replication is one strategy gut bacteria can use to maintain or increase their biomass when cell turnover and intestinal sloughing are high ^41^ as in DSS-induced colitis.^22,23,138,139^

We further report a lack of correlation between gut bacterial minimal doubling times and replication rates during the symptomatic period. This suggests that replication potential and replication rates are also measurements of fundamentally different processes, although this remains to be validated.

It should be noted that our WGS analysis has limitations. We could only assemble approximately 35-36% of the sequences obtained into MAGs, of which only 53-65% could be both taxonomically characterized and have their replication rates estimated using GRiD (**Supplementary Table 15**). This can be partly attributed to the relatively shallow sequencing (median 12,775,255 reads; range 8,448,754 – 19,756,090 reads), resulting in the low coverage of many genomes for most time points. Despite this, we believe that this study provides a foundational representation of the replication dynamics of the most abundant gut taxa during DSS colitis development, and argue that bacterial replication rates should be measured in ongoing microbiome workflows where WGS data is acquired. We have created and shared on GitHub (see Data Accessibility) our workflow for genome assembly and binning, as well as subsequent MAG classification and bacterial replication measurements, to help researchers proceed with such analyses.

### Conclusions and Future Perspectives

Overall, this work demonstrates the utility of tracking various facets of gut bacterial replication alongside measurements of cell abundance and taxonomy during the development of intestinal inflammation. We provide further evidence for the relevance of some taxa, such as *Akkermansia*, and describe new gut bacterial replication dynamics, such as those for *Erysipelatoclostridium*, in a DSS mouse model of colitis.

These taxa should be further studied for their relevance to host health.^64^ For instance, determining the shared metabolic characteristics of gut bacteria which persist, and/or thrive, in the inflamed gut could reveal how these taxa or their metabolism may be linked to the development of an inflammatory environment. Such taxa or metabolic activities could act as biomarkers, wherein changes in their presence could indicate the onset of a flare before overt symptoms manifest. This information could further help predict how a particular gut bacterial community will respond to inflammation, depending on the composition and activity of the community members.^140^

Simplified *in vitro or in vivo* systems (e.g., artificial guts, gut-on-a-chip, or bacterial consortia like the OMM12 community^141^) will be particularly useful, since the complexity of the native gut microbiota and the host environment severely limit our ability to disentangle the myriad microbe-microbe and host-microbe interactions occurring during colitis development. Such approaches could, for example, allow for the direct comparison of transcripts or metabolites produced by thriving and non-thriving gut taxa in an inflammatory intestinal environment.

Evaluating gut bacterial replication dynamics provides insight into bacterial activity which cannot be captured from abundance measurements alone. For instance, quantifying bacterial replication rates could be beneficial when following colonization dynamics in early life, to improve our understanding of how microbial activity is linked to the known succession dynamics observed therein.^33,37^ Characterizing bacterial replication dynamics could also be useful when determining whether members of a probiotic cocktail are actively replicating in the gut after administration,^142^ or how the introduction of a pathogen affects bacterial community replication patterns. On a conceptual level, describing various aspects of gut bacterial replication could help uncover the role of r- and K-selection in the gut. Such research will contribute to increasing our knowledge in a variety of ecosystems on this fundamental, yet poorly described, characteristic of complex microbial communities.

## Methods

### Animals

Equal numbers of 6–8-week-old specific-pathogen free (SPF) male and female wild-type C57BL/6 mice (Jackson Laboratories) were kept at the Goodman Cancer Research Center Animal Facility at McGill University, in accordance with the McGill Ethics Research Board (animal ethics protocol MCGL-7999). For each experiment, there were six mice of each sex, and mice were housed three per cage according to sex, with a total of four cages per experiment.

Before administration of dextran sodium sulfate (DSS), mice were allowed to acclimate to the animal facility for two weeks. On the third week, fecal pellets were collected from the mice on three non-consecutive days. Fecal pellets were collected per cage by putting all mice from a single cage into a new, sterile cage, empty of corncob bedding and nesting material, with no access to food or water. Feces were collected soon after defecation using ethanol-cleaned tweezers reserved for this purpose. Feces from mice of the same cage were put into the same 1.5 mL microcentrifuge tube. This procedure lasted no longer than one hour, after which mice were placed back into their respective cages. Immediately after collection, the fecal pellets were either processed in an anaerobic chamber or frozen at -80°C for later DNA extraction, as detailed below.

The week following acclimatization, mice were administered 2% DSS (molecular weight: 36-50 kDA; MP Biomedicals) in the drinking water for 5 days, and fecal pellets were collected daily except on weekends. Afterwards, the mice were returned to the facility water line, and fecal pellets continued to be collected daily, except on weekends, for another 6 days, after which fecal pellets were collected twice weekly for an additional two to three weeks.

Mouse health parameters, including body weight, blood in stool, and levels of fecal lipocalin-2, were monitored throughout the experiment. Body weight was measured per mouse daily except on weekends. Blood in stool was detected using the Hemoccult Sensa kit (Beckman Coulter). Levels of fecal lipocalin-2 (LCN-2) were quantified using the mouse lipocalin 2 DuoSet enzyme-linked immunosorbent assay (ELISA) (R&D). Fecal samples were prepared for the ELISA using the method of Chassaing et al. (2012)^40^ and diluted 10-fold to 10,000-fold depending on the concentration of LCN-2 in the sample. The ELISA was performed according to the manufacturer’s instructions.

The health states of the mice were determined based on body weight, blood in stool, and fecal lipocalin-2 levels, as detailed in **Supplementary Table 1**. Baseline days were defined as those before DSS administration, during which the mice had no clinical symptoms of colitis. Pre-symptomatic days were defined as those during DSS administration where the mice had minimal symptoms of colitis, including minor weight loss and increased levels of fecal LCN-2 up to 100 ng per gram of feces (ng/g). Symptomatic days were defined as those during or after DSS administration where the mice had weight loss of 1-10%, increased levels of fecal LCN-2 exceeding 100 ng/g, and/or the presence of blood in their stool. Recovery days were defined as the days after DSS administration cessation and during which the mice began to gain weight and no longer had blood in their stool, though high levels (≥100 ng/g) of fecal LCN-2 still remained (**Figure 1**).

We randomly selected fecal samples collected from each cage of mice, rather than tracking each individual mouse, due to our need for large quantities of fecal samples to conduct our experiments. The microbial composition of the mice in a single cage is thought to be very similar due to mouse coprophagic behaviour,^143–145^ and as such, fecal samples taken on a per-cage basis are expected to be broadly representative of all the mice in a single cage.

This experimental setup was conducted independently two times, as detailed in the main text.

### Isolation of bacteria from mouse feces, EdU-click, and flow cytometry

Freshly collected fecal samples from mice were transferred to an anaerobic chamber (Coy Laboratory Products; 5% H2, 20% CO2, 75% N2) within one hour of collection. Isolation of bacteria from mouse feces, EdU labeling, EdU-click, flow cytometry, and cell sorting were performed as described previously.^13^ EdU-positive (EdU^+^) and EdU-negative (EdU^-^) cells and the whole community were sorted based on their level of fluorescence in the appropriate channel and quantified as described previously.^13^

### DNA extraction and 16S rRNA amplicon sequencing and analysis

Bacterial DNA was extracted from sorted cells and mouse fecal pellets using the AllPrep PowerFecal DNA/RNA kit (Qiagen) as previously described.^13^ Extracted DNA underwent library preparation and 16S rRNA gene sequencing of the V4-V5 hypervariable region with 515F/926R primers using the Illumina MiSeq PE250 system at the Université de Québec à Montréal (UQAM) Center of Excellence in Research on Orphan Diseases – Foundation Courtois (CERMO-FC) genomics platform.

Sequenced DNA was analyzed using QIIME2^30^ version 2022.2 as previously described.^13^ Sequences were rarified to the depth of the sample with the fewest reads before conducting alpha and beta diversity analyses. Taxonomic classification and relative abundance measurements were conducted on the quality-controlled, filtered sequences using QIIME2’s machine-learning program trained on the 16S V4-V5 regions from our data set, using the Silva 132 database on 99% operational taxonomic units. Reads present in the sheath fluid but absent in the Whole community samples were identified as contaminants and removed.

Further analysis was performed using phyloseq (version 1.44.0)^146^ in R (version 4.3.0) running in RStudio (2023.03.1). Beta diversity measurements were calculated on weighted or unweighted UniFrac distances, Jaccard indices, or Bray-Curtis dissimilarities using rbiom (version 1.0.3).^147^ Repeated measures, multiple comparisons PERMANOVAs were calculated with 999 permutations to test for significance using the pairwiseAdonis package (version 0.4.1),^148^ with multiple comparisons being corrected using the Benjamini-Hochberg (BH) false discovery rate (FDR) correction. Differential abundance analyses were performed on the combined data from all cages, using ANCOM-BC2 (version 2.1.4)^31^, and correcting for cage origin and the repeated measures nature of the data, as described in *Statistical tests* and shown in the code available on GitHub (see *Data availability*).

### Whole genome shotgun sequencing and analysis

For fecal samples from two cages (F2 and M2, corresponding to a cage of female mice and a cage of male mice, respectively) from the second experiment, DNA extracted from the feces collected from these cages underwent library preparation and whole genome shotgun (WGS) sequencing using the Illumina Miseq PE150 system at SeqCenter (Pittsburgh, PA). The resulting bacterial genomes were assessed for their quality using FastQC (version 0.11.9)^149^ and the resulting FastQC files were visualized using MultiQC (version 1.12).^150^ Sequences were then co-assembled per cage across all sampling days using Megahit (version 1.2.9).^151^ Bowtie2 (version 2.4.4)^152^ was used to remove reads of host origin and to assess the extent to which assembled contigs represented the original, quality-controlled reads. Binning of bacterial genomes from the co-assemblies was performed using CONCOCT (version 1.1.0),^153^ MaxBin2 (version 2.2.7),^154^ and MetaBAT 2 (version 2.14),^155^ followed by DASTool (version 1.1.4),^156^ the latter of which identifies the best bins from the three aforementioned binning tools. Bin quality was assessed using CheckM (version 1.2.0).^157^ Only bins meeting the minimum MIMAGS standards of medium quality genomes^158^ went on to be taxonomically identified using GTDB-Tk (version 2.1.0).^159^ The relative abundance of each bin was calculated based on the length of each genome and its coverage (see code under *Data availability*).

### Quantification of replication rate dynamics

Bins meeting the minimum MIMAGS standards of medium quality (≥50% complete, <10% contamination)^158^ were assessed for their potential maximal replication rates using gRodon2 (version 2.3.0).^35^ The maximal potential replication rate for the entire bacterial community each day was calculated from the prokka (version 1.14.6)^160^ protein annotation output of per-cage, per-day assemblies as specified in the gRodon2 vignette (http://microbialgamut.com/gRodon-vignette), using metagenome mode V2 and with n_le = 1000 and all other parameters as default.

Bins meeting the minimum standards for GRiD (>=50% completeness, <=90 scaffolds/Mbp, >=x0.2 coverage) were assessed with this tool using default parameters. An index was created for each bin using Bowtie2, and each sample (i.e., each day) per cage was mapped to each appropriate index (i.e., each bin) that came from that cage. The resulting SAM files from mapping were used as input for GRiD, using the “grid single” mode.

### Correlations

To correlate gut bacterial replication rates with their relative abundances, we used the approach of Joseph TA et al. (2020).^16^ Briefly, we first log2-transformed all relative abundance and replication rate (GRiD) values. Then, we standardized the log2-transformed relative abundance and GRiD values per taxon using the scale() function in R. The standardized, log2-transformed values for abundance and GRiD were then correlated using Pearson correlation. We proceeded with the same approach to correlate gut bacterial replication potentials with their corresponding median replication rates during the symptomatic period, and to correlate gut bacterial replication potential with their corresponding median relative abundances during the symptomatic period.

### Statistical tests

Statistical testing was performed according to the non-parametric, repeated measures (i.e., longitudinal), and multiple comparisons nature of the study design as necessary, as described in the results and/or figure legends, where appropriate.

Comparisons of the proportion of EdU^+^ cells (%EdU^+^ cells) between health states were performed by first calculating the median %EdU^+^ cells per cage per health state, as the median is more robust to outliers than the mean. Then, the median %EdU^+^ cells per health state, considering all four cages in each health state, were compared to each other using the Friedman test, accounting for cage as a random intercept effect.

Comparisons of alpha diversity metric values (Shannon index, Simpson evenness, and observed features) between health states was performed by first calculating the median alpha diversity value per cage per health state. Then, the median alpha diversity values per health state, considering all four cages in each health state, were compared to each other using the Friedman test, accounting for cage as a random variable. Where appropriate, the Wilcoxon sign-rank post-hoc test was conducted, wherein the baseline group was compared to all of the other health states, with the p-value adjusted for multiple comparisons using the Bonferroni correction.

PERMANOVA comparisons of the beta diversity between different health states of mice were conducted using pairwiseAdonis (version 0.4.1).^148^ These comparisons accounted for the repeated measures design of the study as described in the available code and as first described here: https://thebiobucket.blogspot.com/2011/04/repeat-measure-adonis-lately-i-had-to.html. Multiple comparisons were conducted using the pairwiseAdonis package in R. When data from the various cages were compiled together, cage origin was considered as a random intercept effect.

ANCOM-BC2 (version 2.1.4)^31^ was used to detect differentially abundant bacteria between health states as described in the main text and by the creators of the ANCOM-BC2 package. All health states were compared to the baseline state, using Dunnett’s test. When data from all cages of mice were compiled together, cage origin was considered as a random intercept effect.

To determine to what extent the health state of the host could explain the minimal doubling time of the whole community of bacteria, we fit a linear mixed-effects model to our data using the lme4 (version 1.1-34)^161^ and lmerTest (version 3.1-3)^162^ packages in R, with health state and minimal doubling time as fixed effects, and cage and sampling day as random intercept effects.

## Data availability

Bacterial 16S rRNA gene sequencing data from all experimental arms are available in the NCBI SRA database under project number PRJNA1040262 and PRJNA1040873, respectively. Whole genome shotgun sequencing data from the second experiment is available in the NCBI SRA database under project number PRJNA1040965. Code related to the analysis of these data have been deposited in GitHub under the account at https://github.com/evetb, repository DSS_Manuscript. Raw flow cytometry data have been deposited in FigShare (https://figshare.com/projects/DSS_Manuscript/193004).

## Declarations

### Authors’ contributions

ETB designed, performed, analyzed, and interpreted experiments. CH aided with extracting DNA and supernatant for lipocalin-2 assays for the first mouse experiment. CFM helped develop the project, obtained funding, and helped design experiments and interpret the data. ETB and CFM edited the manuscript and approved the final draft.

