## Supplementary Figures for "Dextran sodium sulfate-induced colitis alters the proportion and composition of replicating gut bacteria"

**A.**

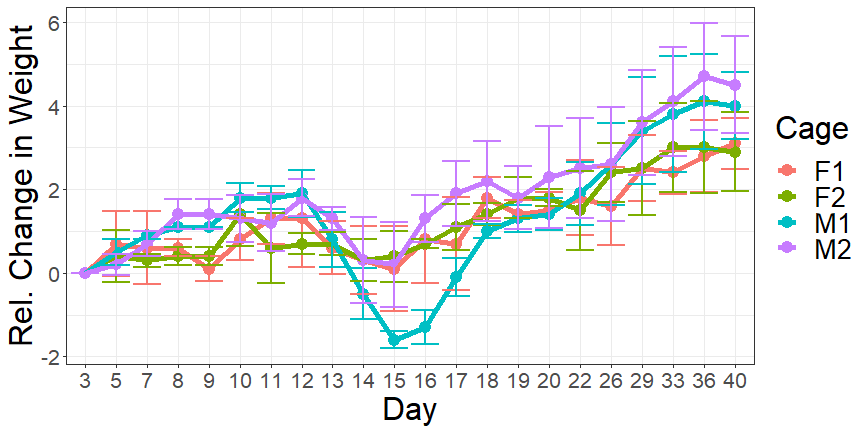

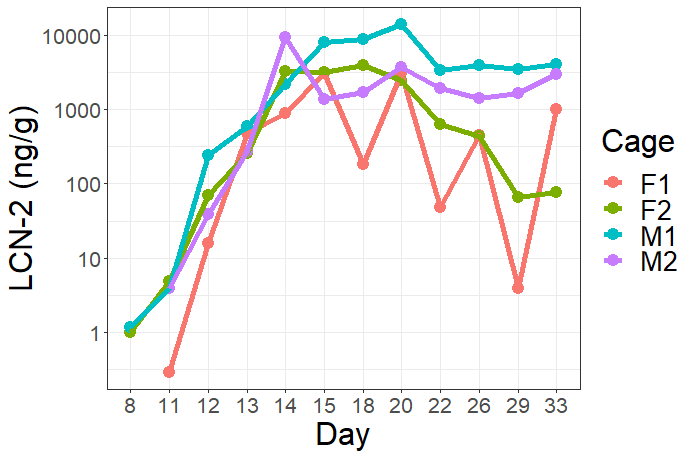

**C.**

**B.**

| **Date** | **Day** | **Health State** | **Cage** | | | |
| --- | --- | --- | --- | --- | --- | --- |
|  |  |  | F1 | F2 | M1 | M2 |
| 22 Oct 2020 | 8 | Baseline | - | - | - | - |
| 25 Oct 2020 | 11 | Pre-symptomatic | - | - | - | - |
| 26 Oct 2020 | 12 | Pre-symptomatic | - | - | + | + |
| 27 Oct 2020 | 13 | Symptomatic | + | + | + | + |
| 28 Oct 2020 | 14 | Symptomatic | + | + | + | - |
| 29 Oct 2020 | 15 | Symptomatic | - | - | + | - |
| 01 Nov 2020 | 18 | Recovery | - | - | - | - |
| 03 Nov 2020 | 20 | Recovery | - | - | - | - |
| 05 Nov 2020 | 22 | Recovery | - | - | - | - |

**Supplementary Figure 1.** Mouse health parameters during first DSS colitis experiment.

Three main indicators of colitis were followed in four cages of mice before, during, and after the administration of DSS. (A) Relative change in mouse weight, relative to their starting weight at the beginning of the experiment. (B) Levels of lipocalin-2 (LCN-2) reported as nanograms per gram of feces (ng/g). (C) Presence (+) or absence (-) of blood in the feces of mice in each cage, as determined by hemoccult analysis

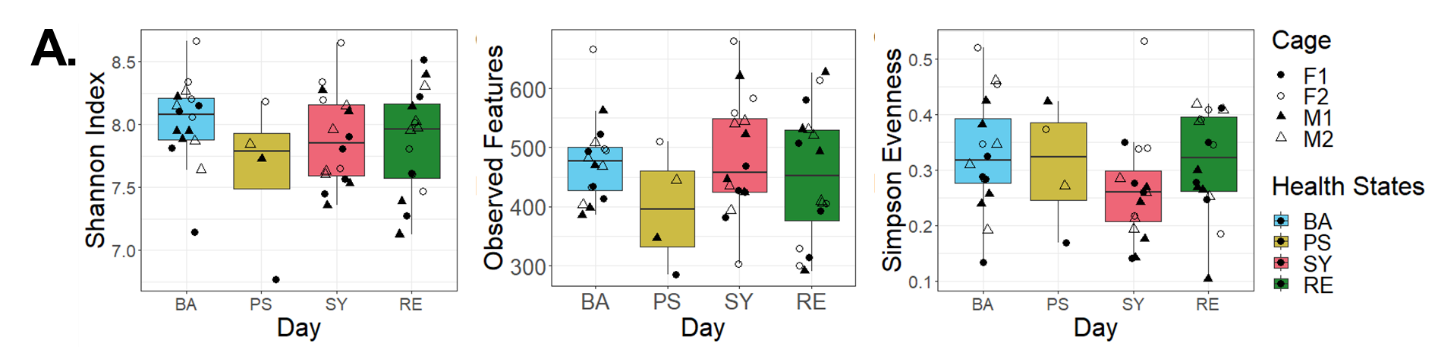

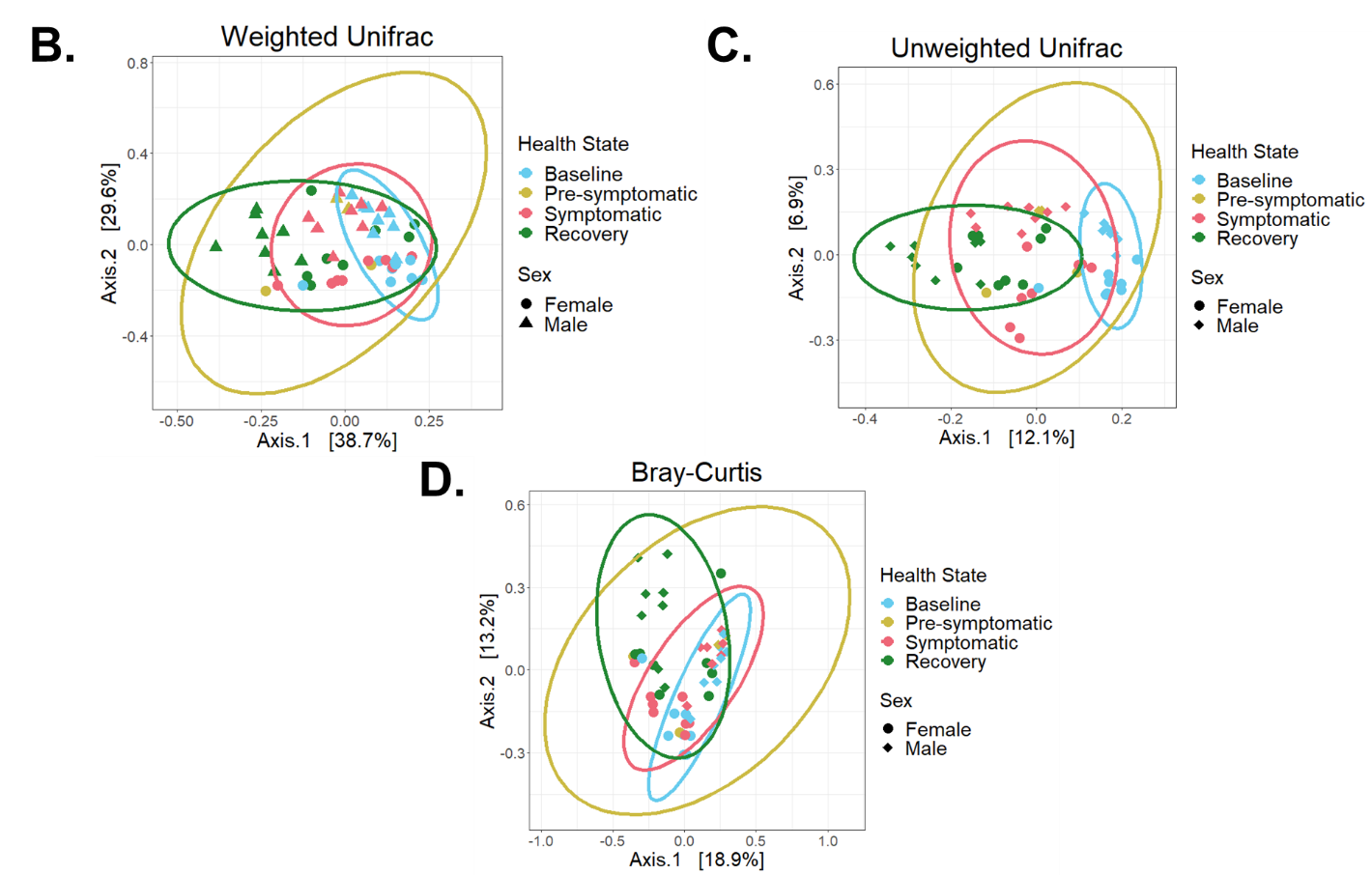

**Supplementary Figure 2.** Alpha and beta diversity statistics for first DSS mouse experiment.

(A) Alpha diversity for all cages combined – Shannon index (left), observed features (middle), and Simpson evenness (right). (B-D) Beta diversity for all cages combined. Colors of dots and ellipses/grouping determined by health state. (B) Weighted Unifrac distance (WU), (C) Unweighted Unifrac distance (UU), and (D) Bray-Curtis dissimilarity (BC).

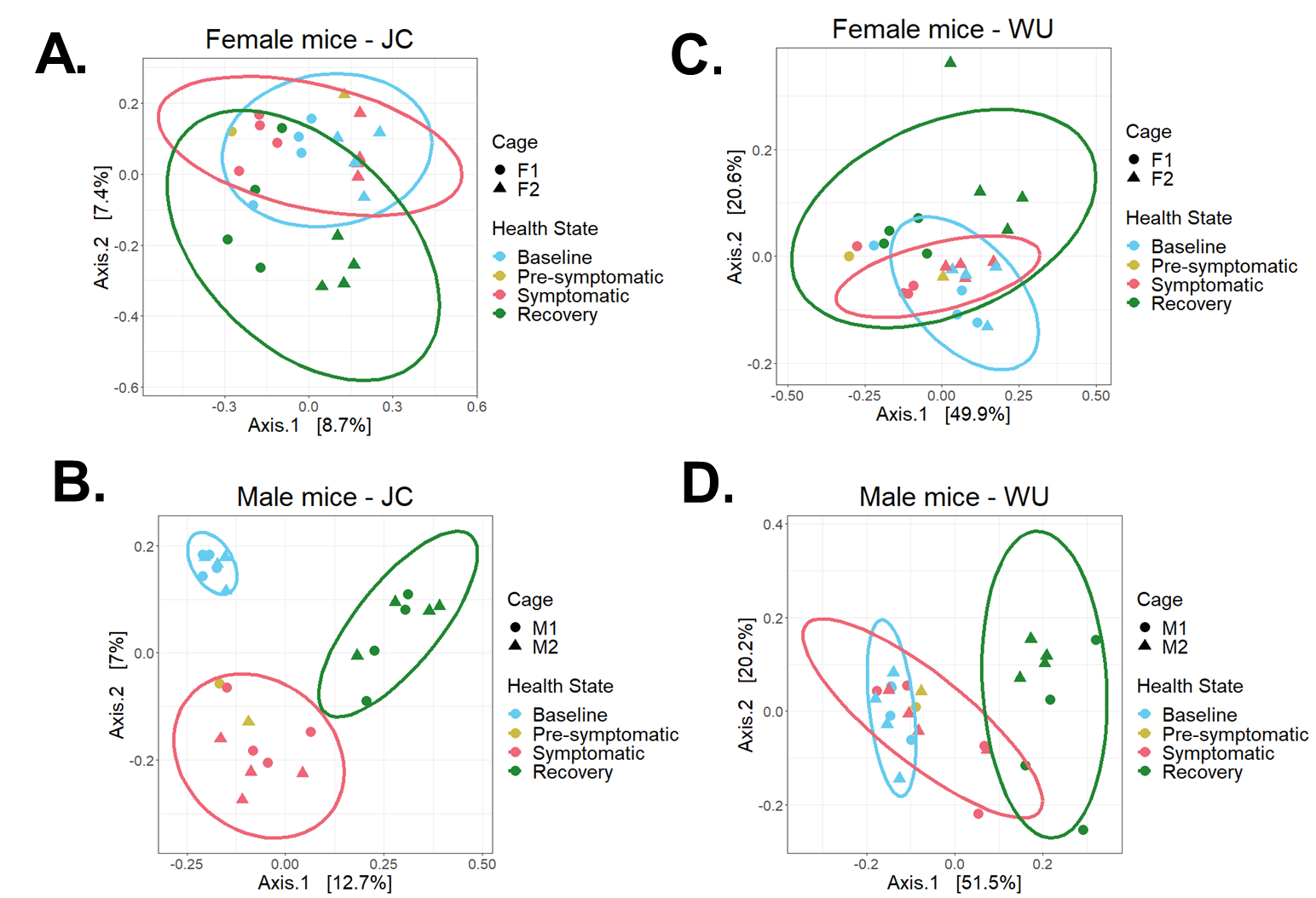

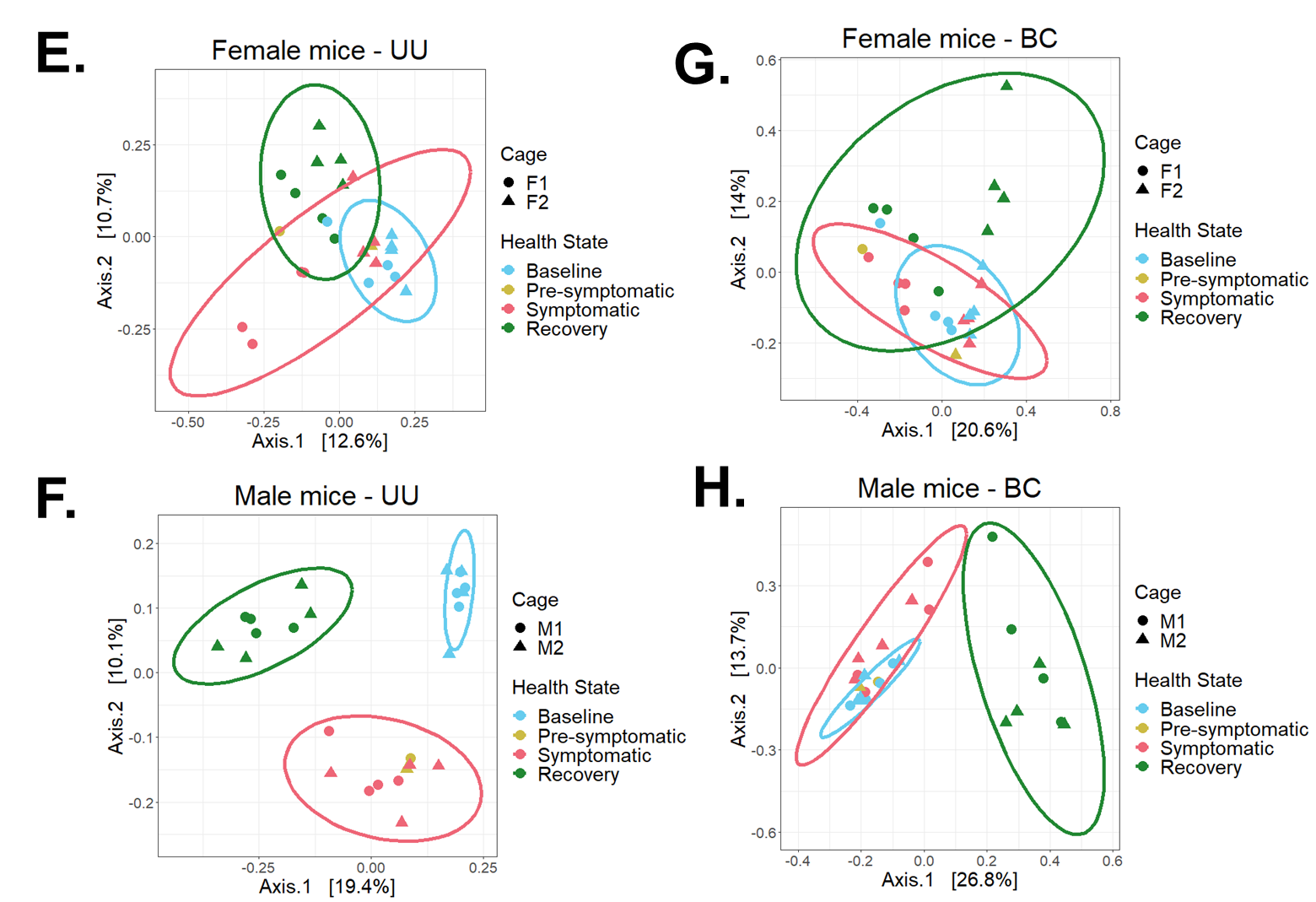

**Supplementary Figure 3.** Beta diversity statistics per sex, grouped by health state.

Beta diversity statistics per sex, grouped by health state, for the first DSS experiment. (A-B) Jaccard index (JC) (C-D) Weighted Unifrac distance (WU). (E-F) Unweighted Unifrac distance (UU) (G-H) Bray-Curtis dissimilarity (BC)

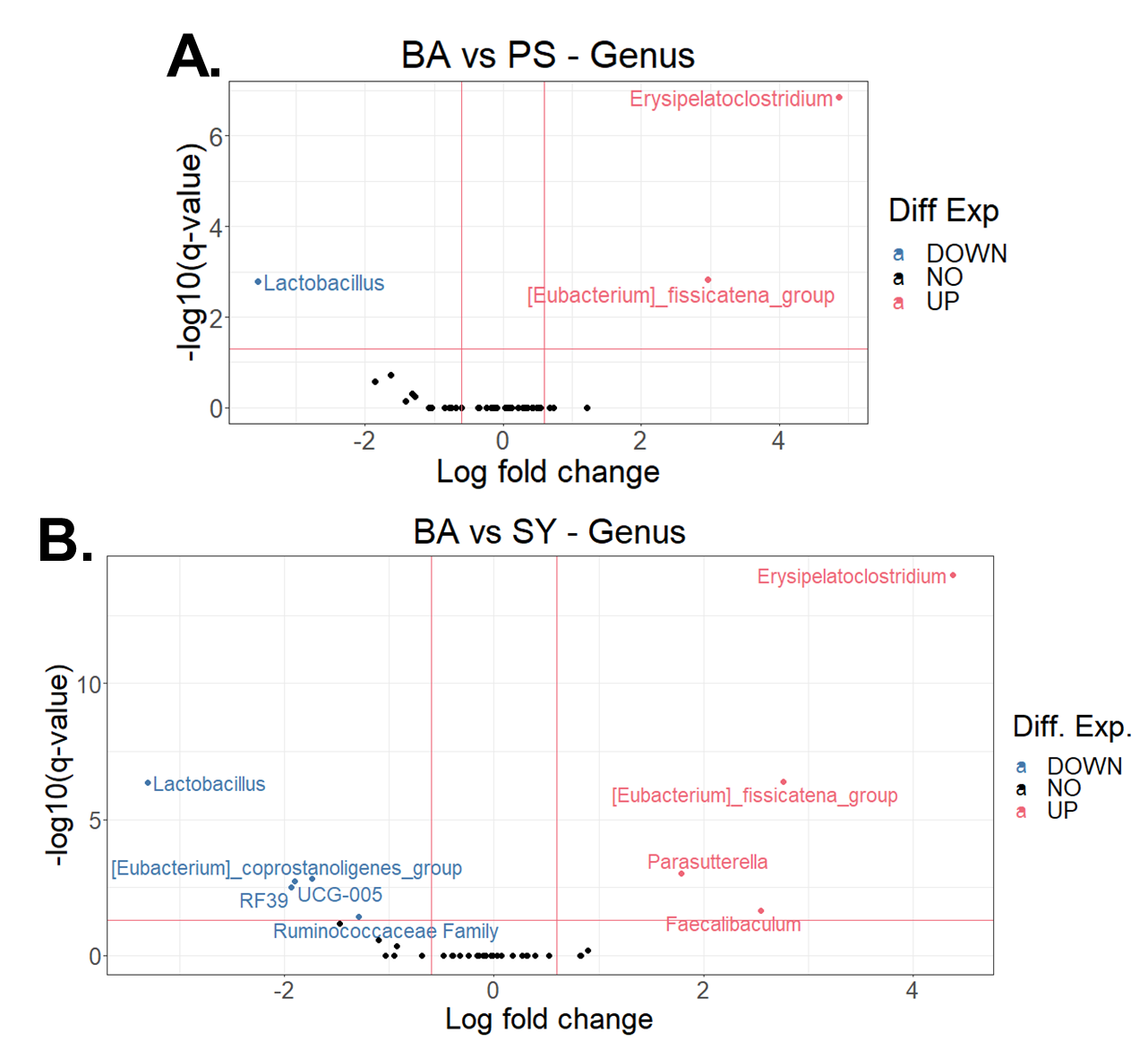

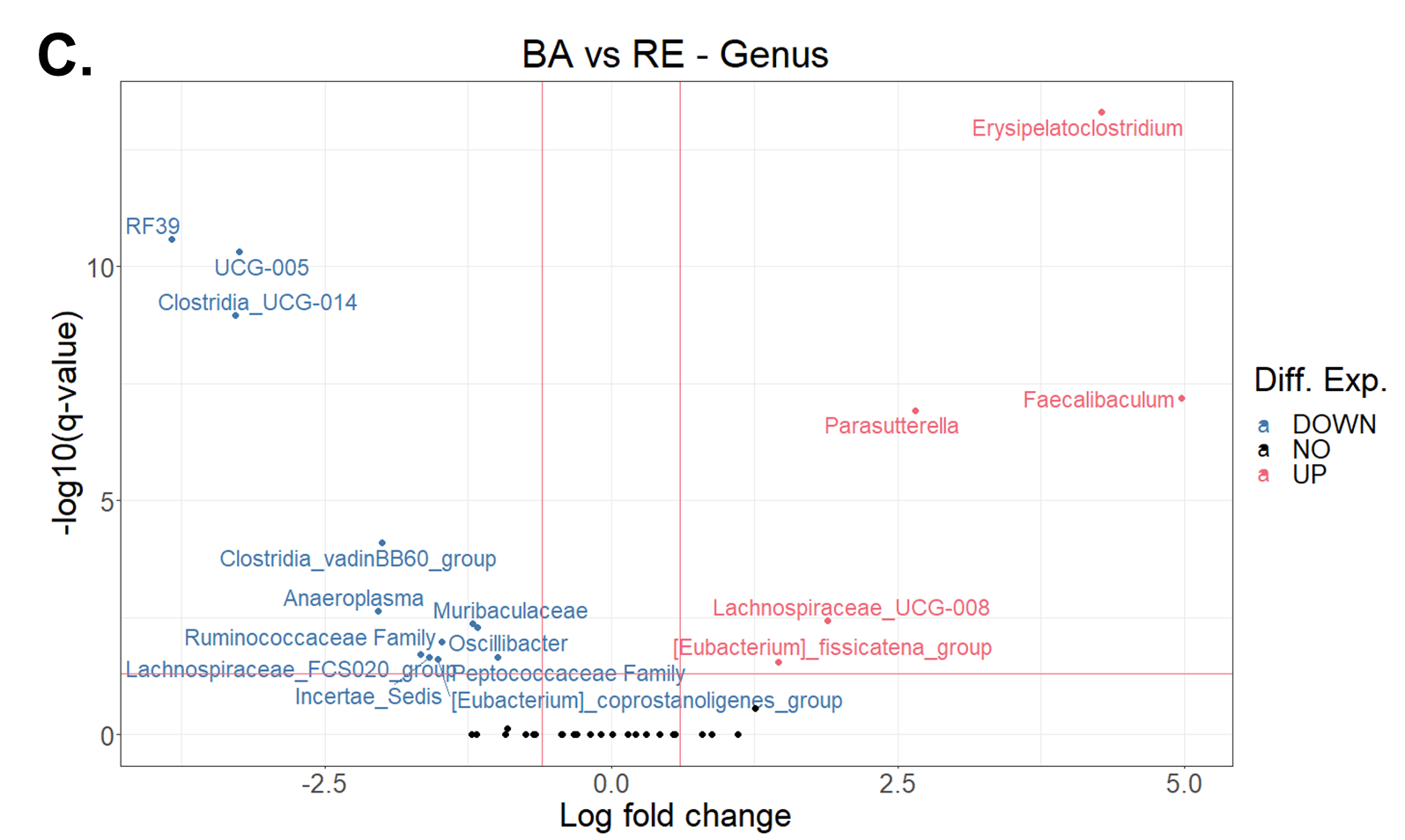

**Supplementary Figure 4**. Taxa marked as differentially abundant as compared to the baseline state during DSS colitis.

Taxa marked as differentially abundant as compared to the baseline state, during the first DSS colitis experiment. (A-C) Volcano plots showing the differentially abundant taxa at the genus level, when comparing from the baseline state (BA) to the (A) pre-symptomatic (PS), (B) symptomatic (SY), and (C) recovery (RE) state. Taxa with increased abundances (Diff Exp = UP) during the health state of interest (pre-symptomatic, symptomatic, recovery) versus baseline are in red/pink on the upper right of each plot, whereas taxa with increased abundances in the baseline state (Diff Exp = DOWN) as compared to the health state of interest are in blue, on the upper left of each plot.

**A.**

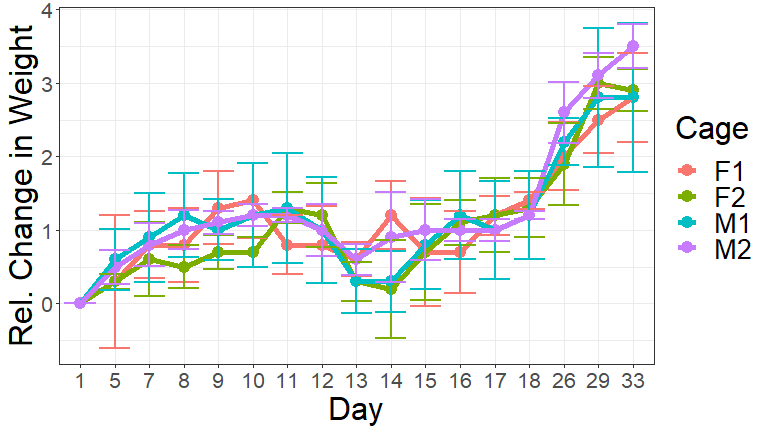

**B.**

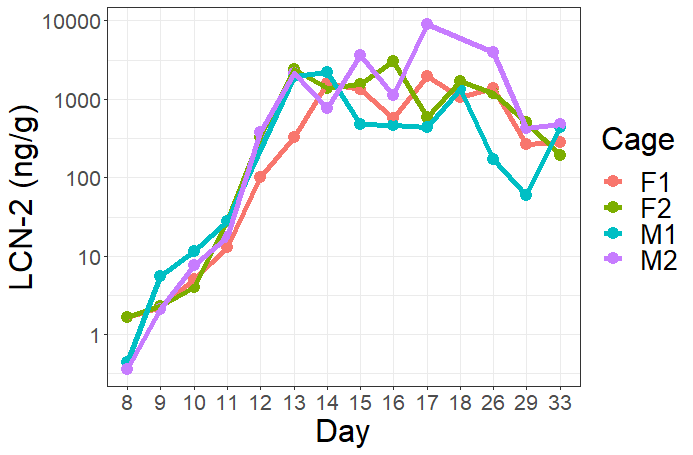

**C.**

| **Date** | **Cage** | | | | **Day** | **Health State** |
| --- | --- | --- | --- | --- | --- | --- |
|  | F1 | F2 | M1 | M2 |  |  |
| 22 April 2021 | - | - | - | - | 8 | Pre-symptomatic |
| 23 April 2021 | - | - | -/+ | - | 9 | Pre-symptomatic |
| 24 April 2021 | - | - | - | - | 10 | Pre-symptomatic |
| 25 April 2021 | - | ~ | - | ~ | 11 | Pre-symptomatic |
| 26 April 2021 | - | +/- | +/- | + | 12 | Symptomatic |
| 27 April 2021 | ~ | - | + | + | 13 | Symptomatic |
| 28 April 2021 | - | - | - | - | 14 | Symptomatic |
| 29 April 2021 | - | - | - | - | 15 | Symptomatic |
| 30 April 2021 | - | - | - | - | 16 | Recovery |

**Supplementary Figure 5**. Mouse health parameters during second DSS colitis experiment.

Three main indicators of colitis were followed in four cages of mice before, during, and after the administration of DSS. (A) Relative change in mouse weight, relative to their starting weight at the beginning of the experiment. (B) Levels of lipocalin-2 (LCN-2) reported as nanograms per gram of feces (ng/g). (C) Presence (+) or absence (-) of blood in the feces of mice in each cage, as determined by hemoccult analysis. When some fecal samples had a positive signal and others had a negative signal for presence of blood in stool, the symbol “-/+” or “+/-“ is used. For indeterminate results, the tilde symbol “~” is used.

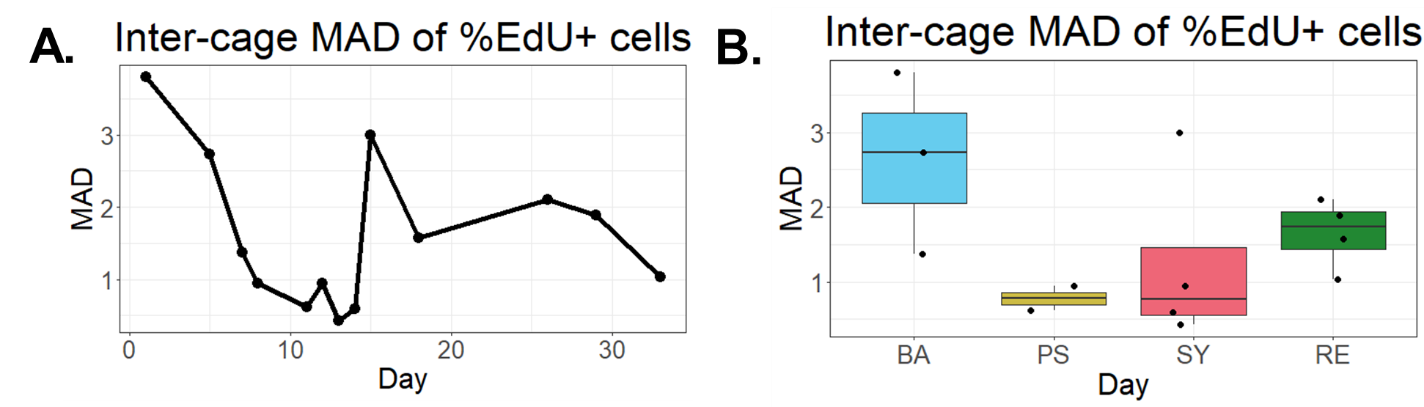

**Supplementary Figure 6**. Median absolute deviation (MAD) of replicating (EdU^+^) cells.

(A-B) MAD of the proportion of replicating (EdU^+^) cells between cages over time in each health state, visualized as a line plot (A) and as boxplots with per-cage MAD values grouped per health state. BA = baseline, PS = pre-symptomatic, SY = symptomatic, RE = recovery.

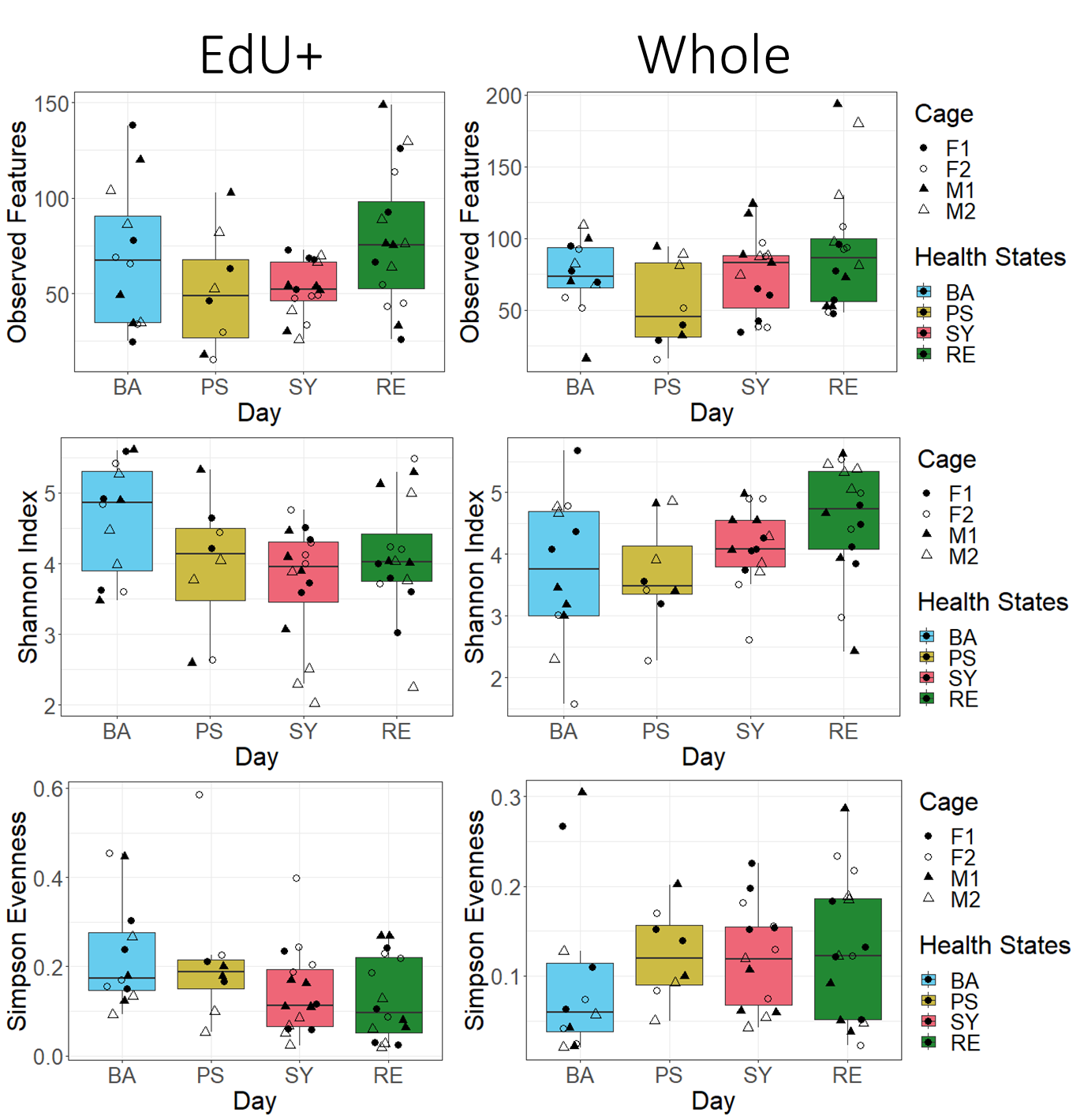

**Supplementary Figure 7**. Alpha diversity metrics during DSS colitis

Observed features, Shannon index, and Simpson evenness for data from all cages combined during the second DSS mouse experiment, for both the replicating (EdU^+^) cells and the whole community (Whole) of bacterial cells. BA = baseline, PS = pre-symptomatic, SY = symptomatic, RE = recovery.

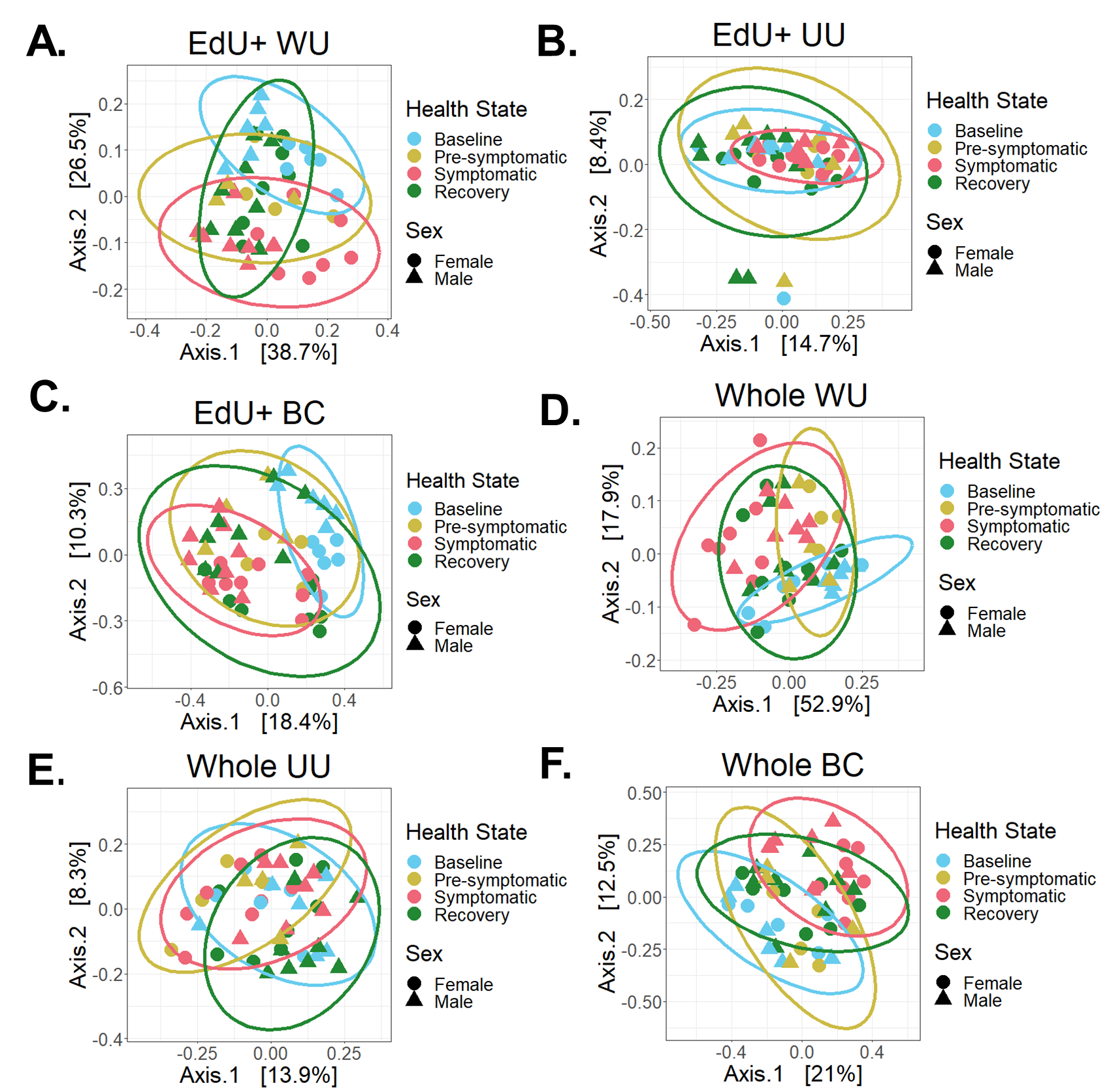

**Supplementary Figure 8**. Beta diversity of the replicating (EdU^+^) and whole community (Whole) of bacterial cells.

(A-C) Beta diversity for the replicating bacterial cells, grouped by health state. (A) Weighted Unifrac distance (WU) (B) Unweighted Unifrac distance (UU) (C) Bray-Curtis dissimilarity (BC). (D-F) Beta diversity for the whole community of bacterial cells, grouped by health state. (D) Weighted Unfirac distance (WU) (E) Unweighted Unifrac distance (UU) (F) Bray-Curtis dissimilarity (BC).

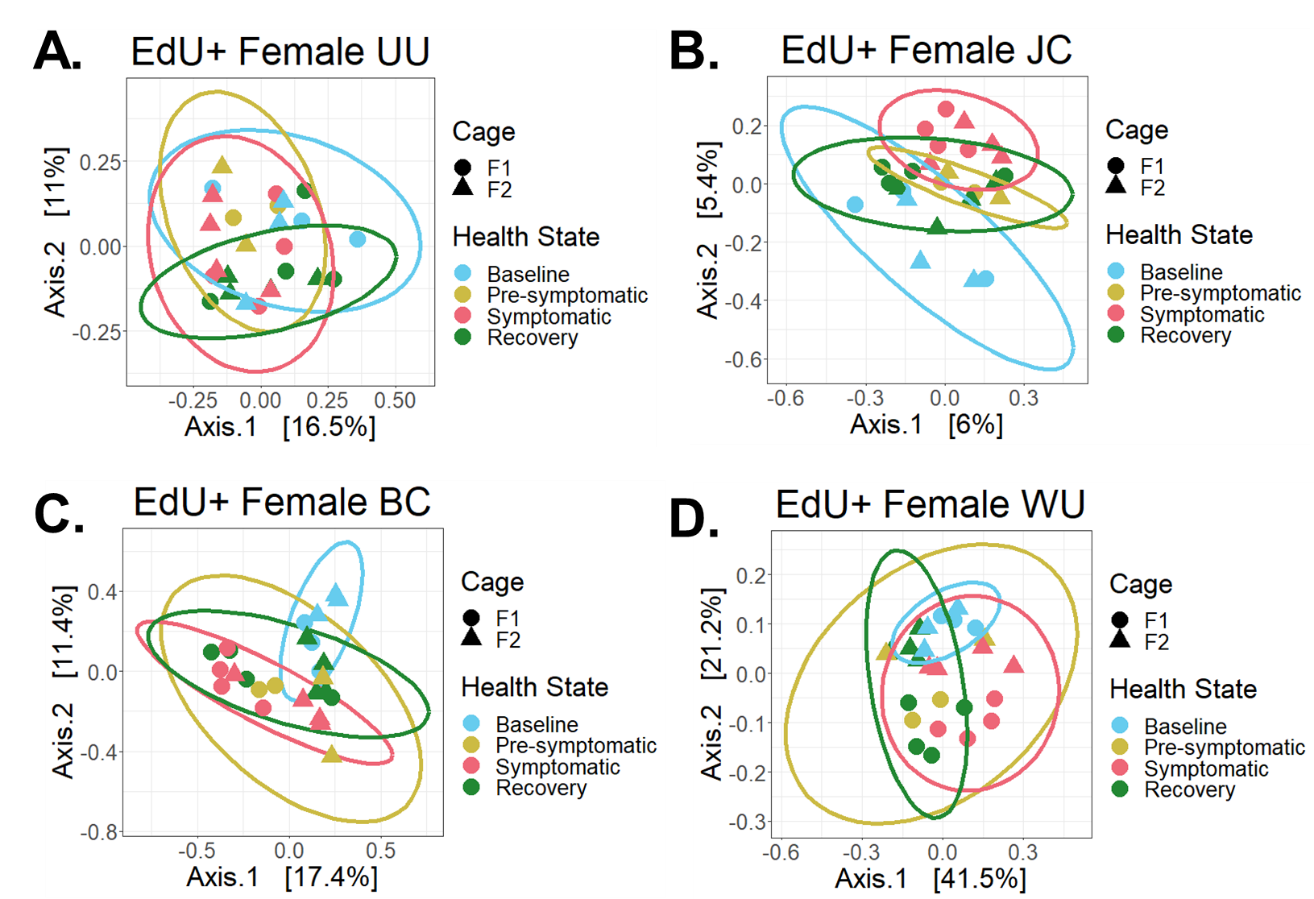

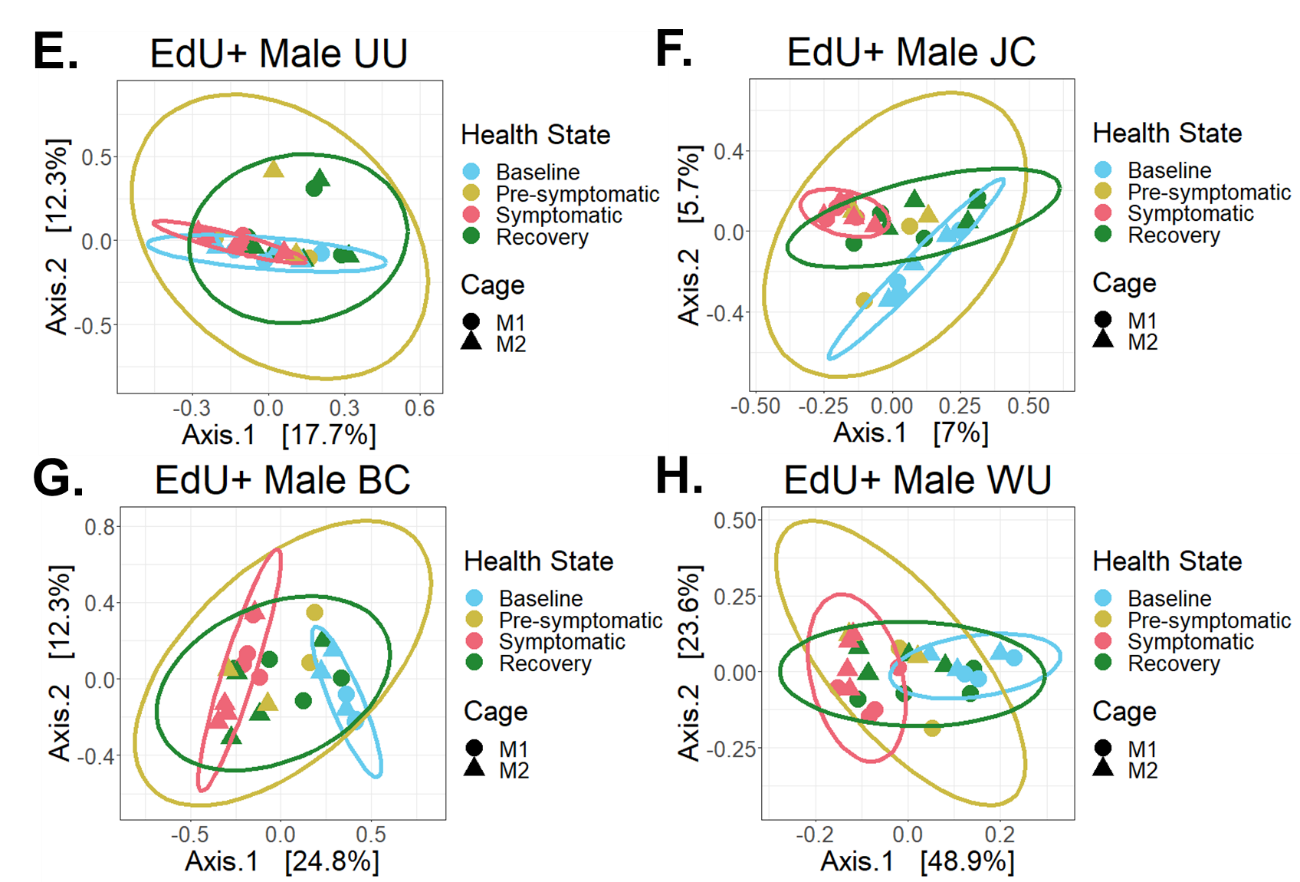

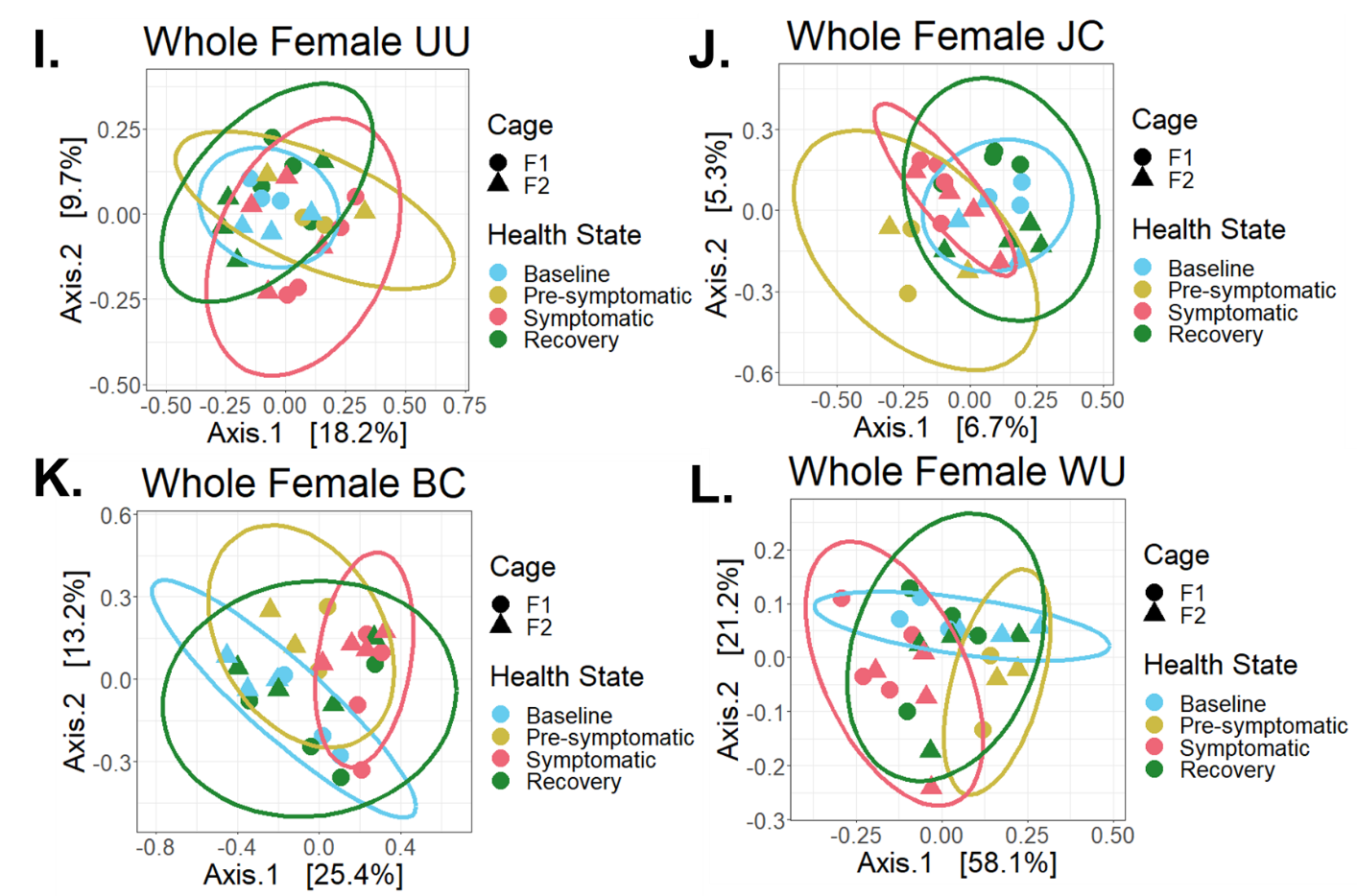

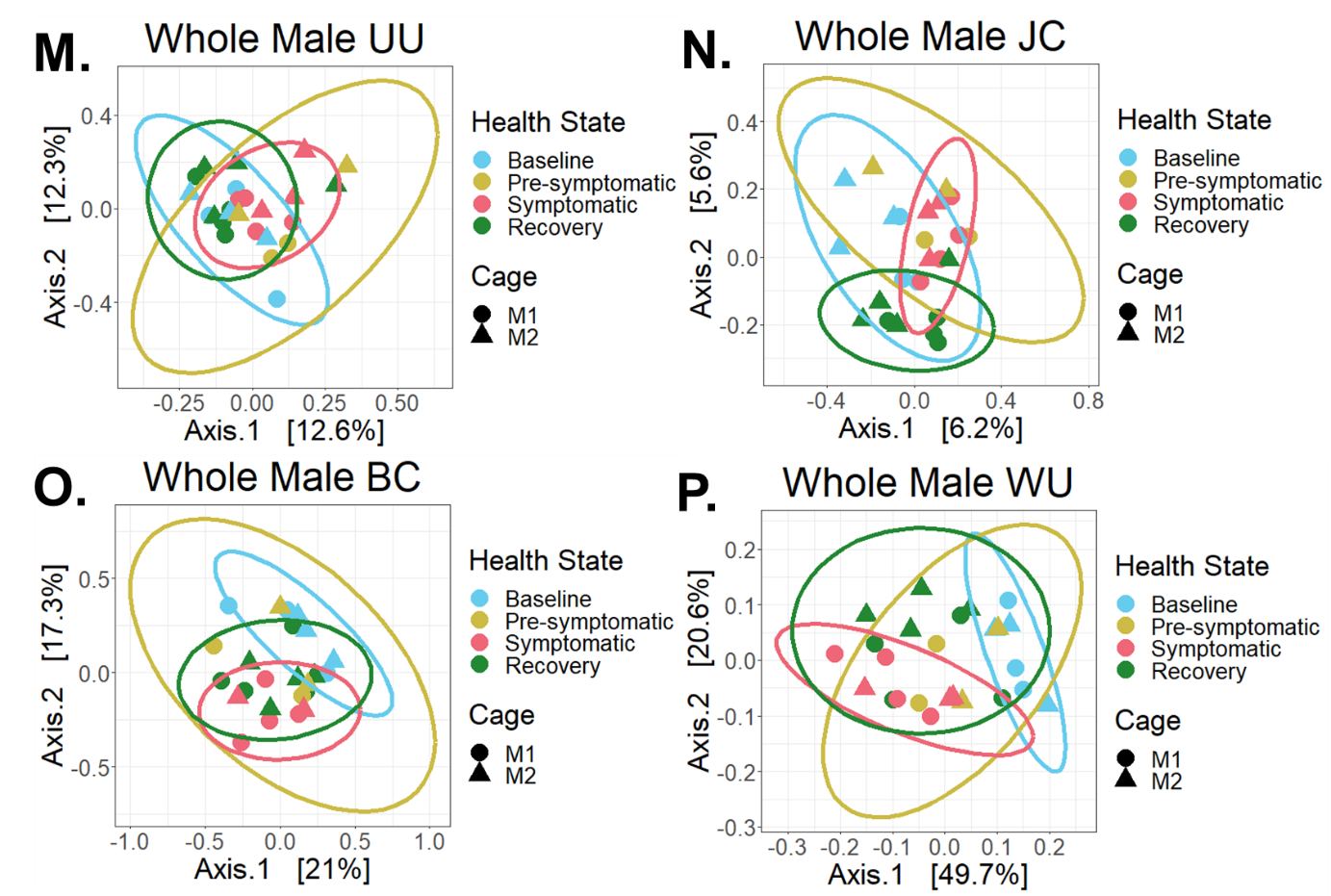

**Supplementary Figure 9**. Beta diversity statistics grouped per sex for each sorted fraction.

(A-D) Beta diversity statistics for the replicating (EdU^+^) bacteria of both cages of Female mice. (A) Unweighted Unifrac distance (UU) (B) Jaccard index (JC) (C) Bray-Curtis dissimilarity (BC) (D) Weighted Unifrac distance (WU). (E-H) Beta diversity statistics for the replicating (EdU^+^) bacteria of both cages of Male mice. (E) Unweighted Unifrac distance (UU) (F) Jaccard index (JC) (G) Bray-Curtis dissimilarity (BC) (H) Weighted Unifrac distance (WU). (I-L) Beta diversity statistics for the whole community of bacteria of both cages of Female mice. (I) Unweighted Unifrac distance (UU) (J) Jaccard index (JC) (K) Bray-Curtis dissimilarity (BC) (L) Weighted Unifrac distance (WU). (M-P) Beta diversity statistics for the whole community of bacteria of both cages of Male mice. (M) Unweighted Unifrac distance (UU) (N) Jaccard index (JC) (O) Bray-Curtis dissimilarity (BC) (P) Weighted Unifrac distance (WU).

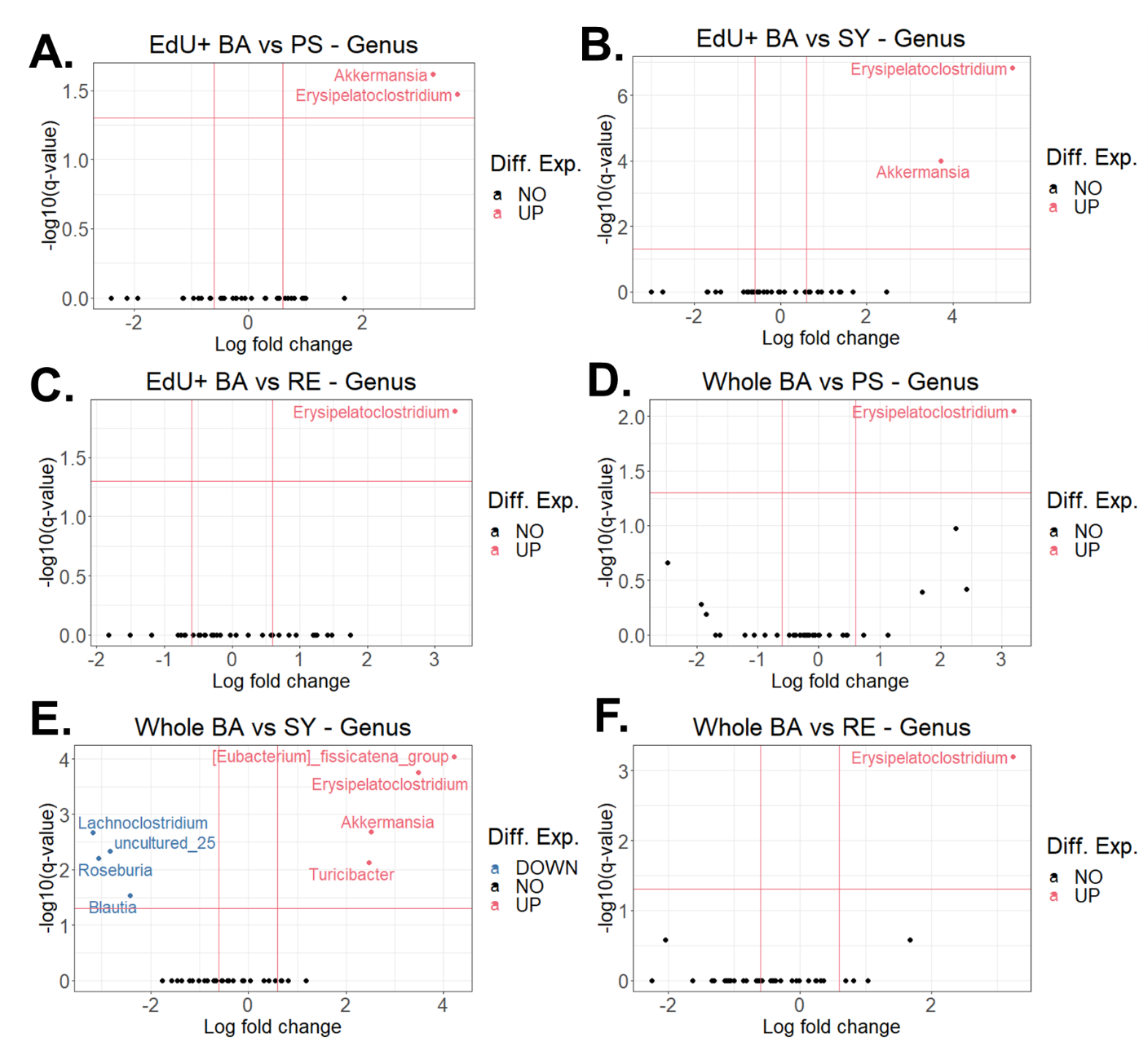

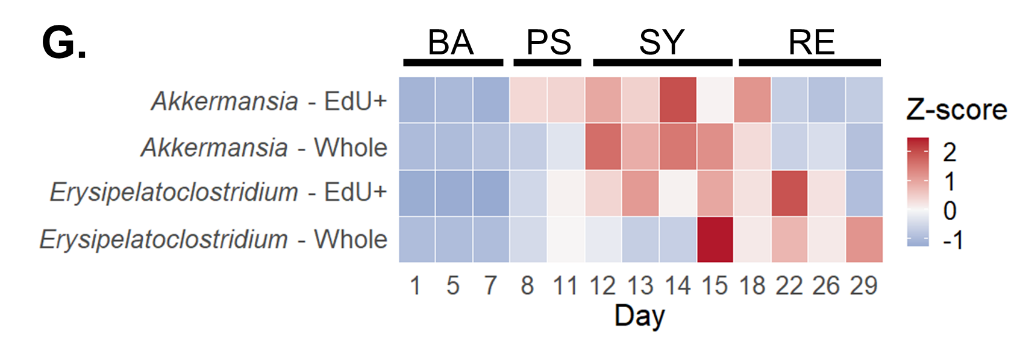

**Supplementary Figure 10**. Differentially abundant taxa for replicating (EdU^+^) and whole community of bacteria.

(A-C) Differentially abundant taxa as compared to the baseline state for replicating (EdU^+^) cells in the second DSS experiment during the (A) pre-symptomatic (PS) (B) symptomatic (SY) and (C) recovery (RE) state. (D-F) Differentially abundant taxa as compared to the baseline state for the whole community of bacterial cells cells during the (D) pre-symptomatic (PS) (E) symptomatic (SY) (F) recovery (RE) state. Taxa with increased abundances during the health state of interest (pre-symptomatic, symptomatic, recovery) versus baseline are in red/pink on the upper right of each plot, whereas taxa with increased abundances in the baseline state as compared to the health state of interest are in blue, on the upper left of each plot. (G) Heat map of standardized abundances of Akkermansia and Erysipelatoclostridium in both the replicating (EdU^+^) and whole community (Whole) fractions during each health state. BA = baseline; PS = pre-symptomatic; SY = symptomatic; RE = recovery.

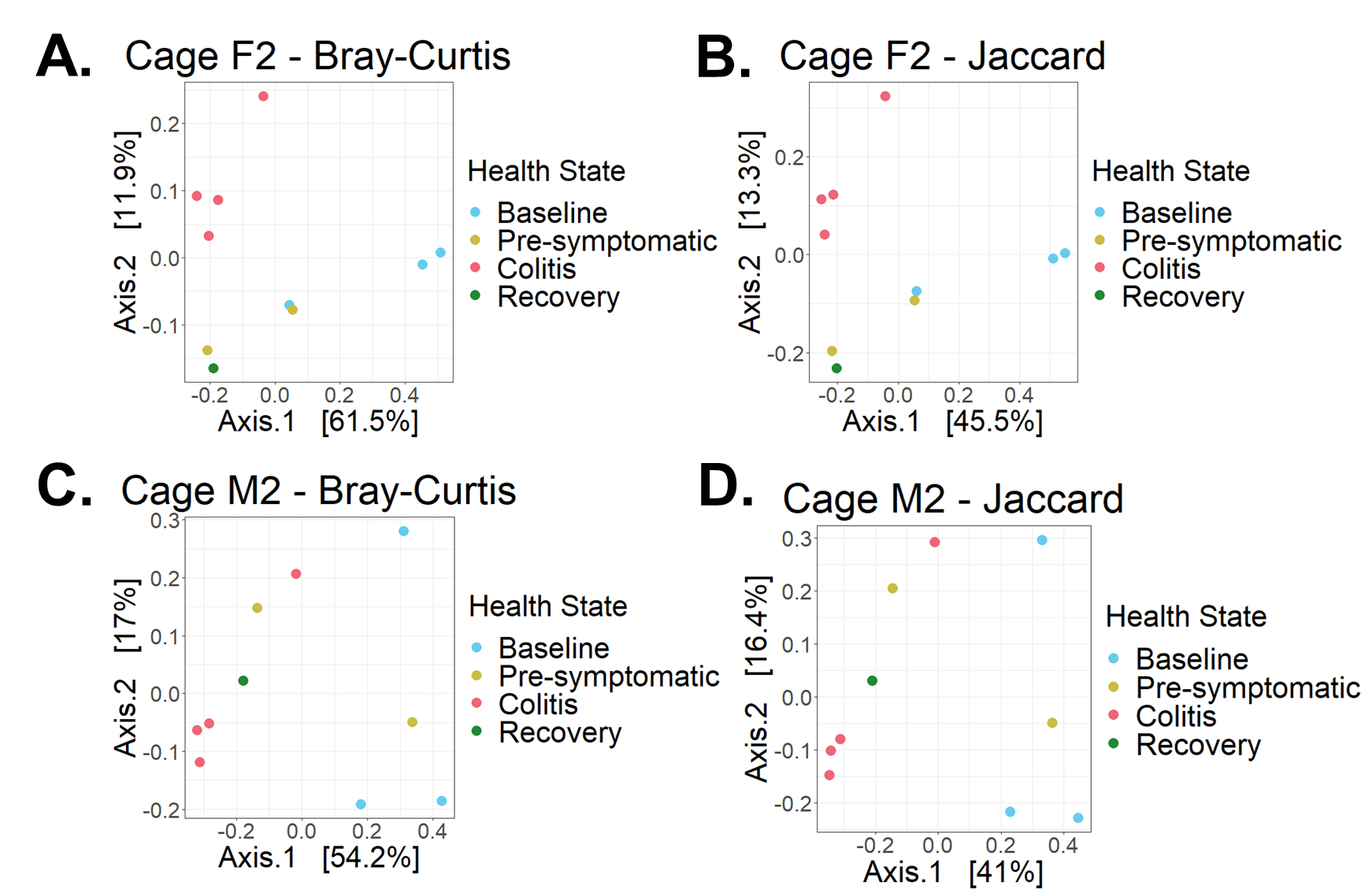

**Supplementary Figure 11**. Beta diversity of metagenome assembled genomes (MAGs).

(A-B) Beta diversity for the community of MAGs from cage F2 for Bray-Curtis dissimilarity (A) and Jaccard index (B). (C-D) Beta diversity for the community of MAGs from cage M2 for Bray-Curtis dissimilarity (C) and Jaccard index (D).

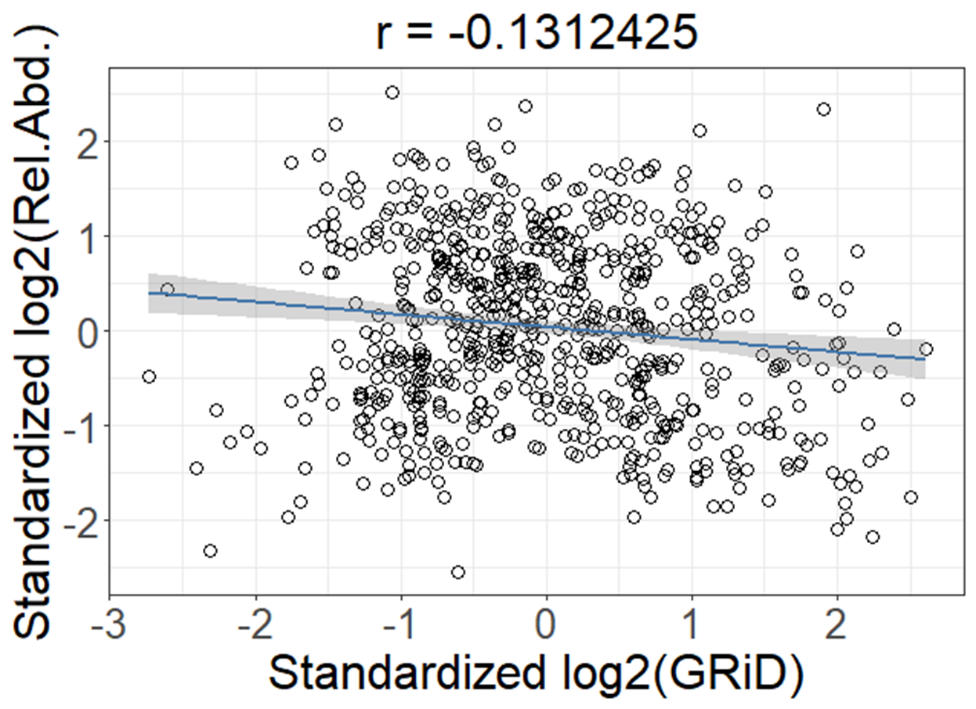

**Supplementary Figure 12**. Correlation between replication rates and relative abundances.

Standardized, log2-transformed values of GRiD-calculated replication rates were correlated using Pearson correlation with standardized log2-transformed values of corresponding metagenome assembled genomes.

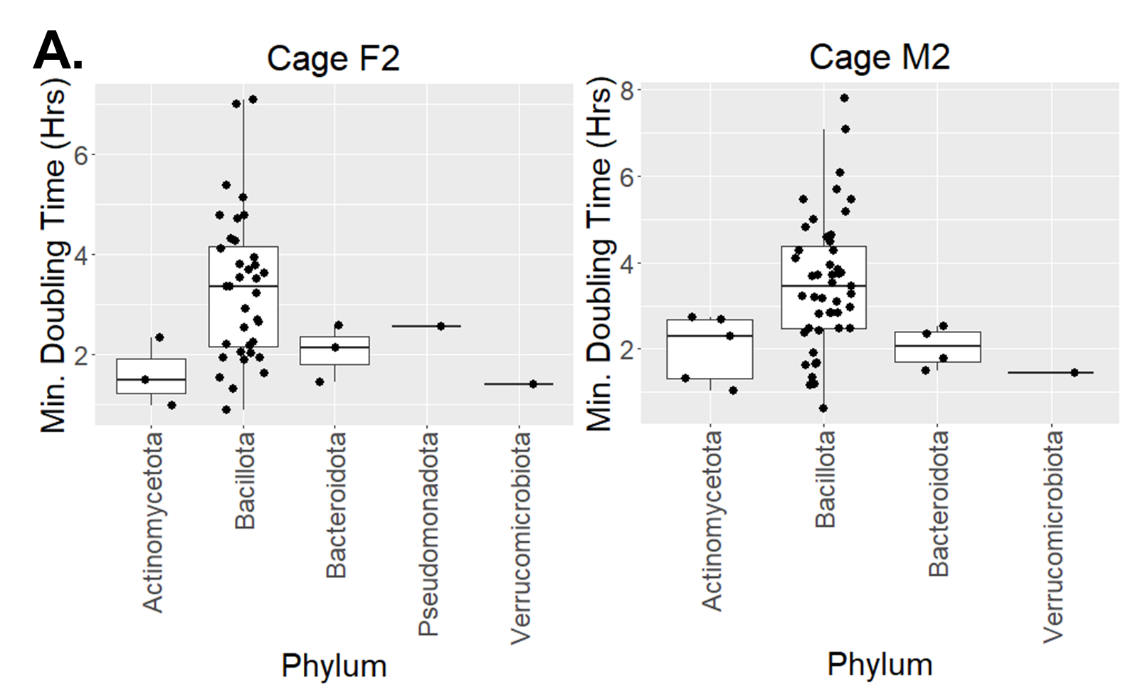

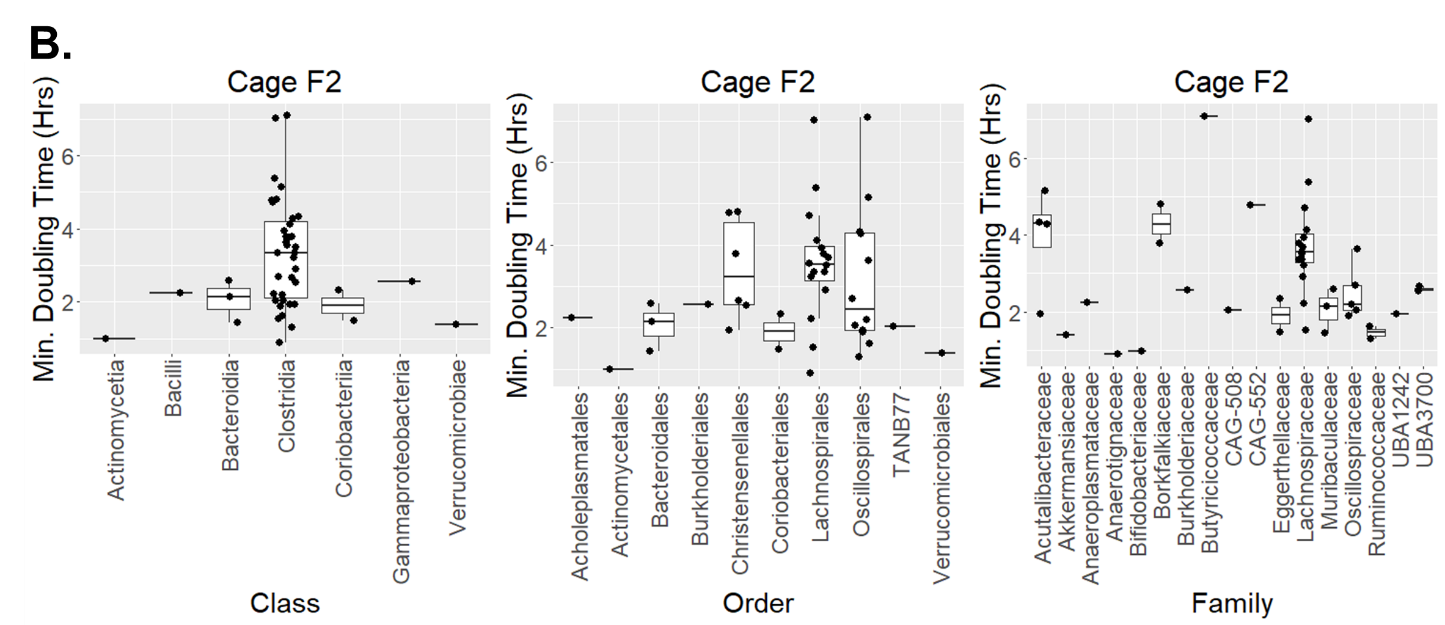

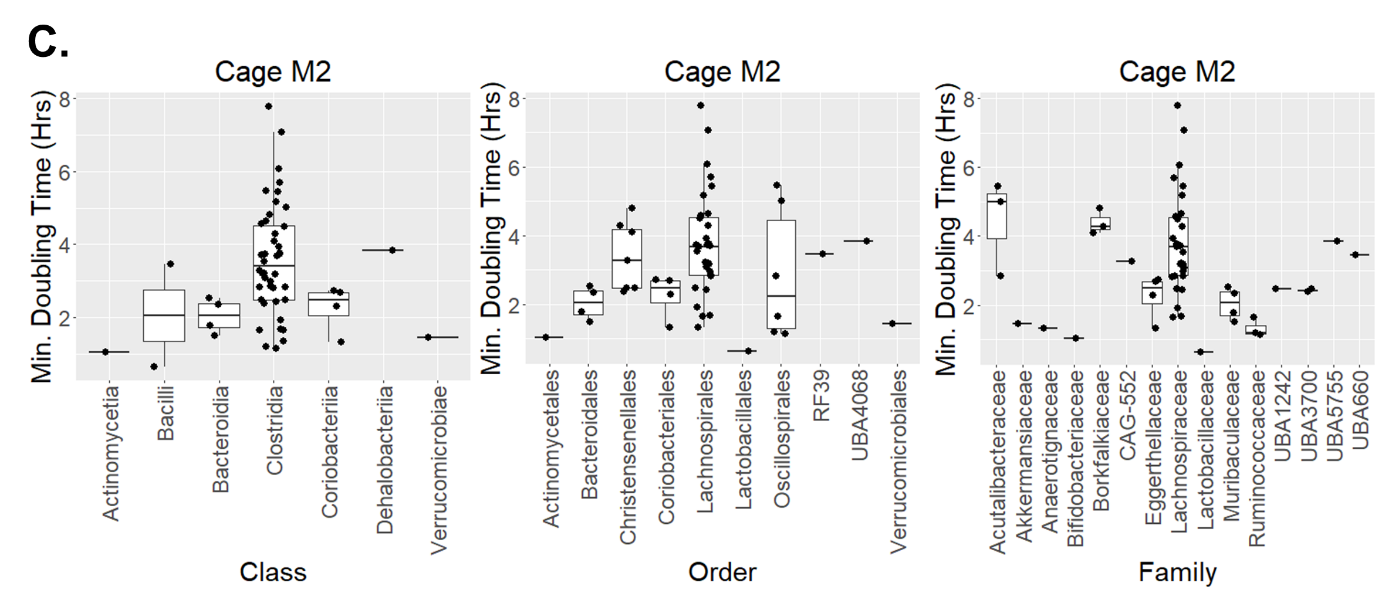

**Supplementary Figure 13**. Minimal doubling times for metagenome-assembled genomes (MAGs) at various taxonomic levels.

(A) Estimated minimal doubling times for each MAG, as calculated by gRodon, at the phylum level for cages F2 (left) and M2 (right). (B) Estimated minimal doubling times for each MAG, as calculated by gRodon, at the class (left), order (middle), and family (right) level for cage F2. (C) Estimated minimal doubling times for each MAG, as calculated by gRodon, at the class (left), order (middle), and family (right) level for cage M2.

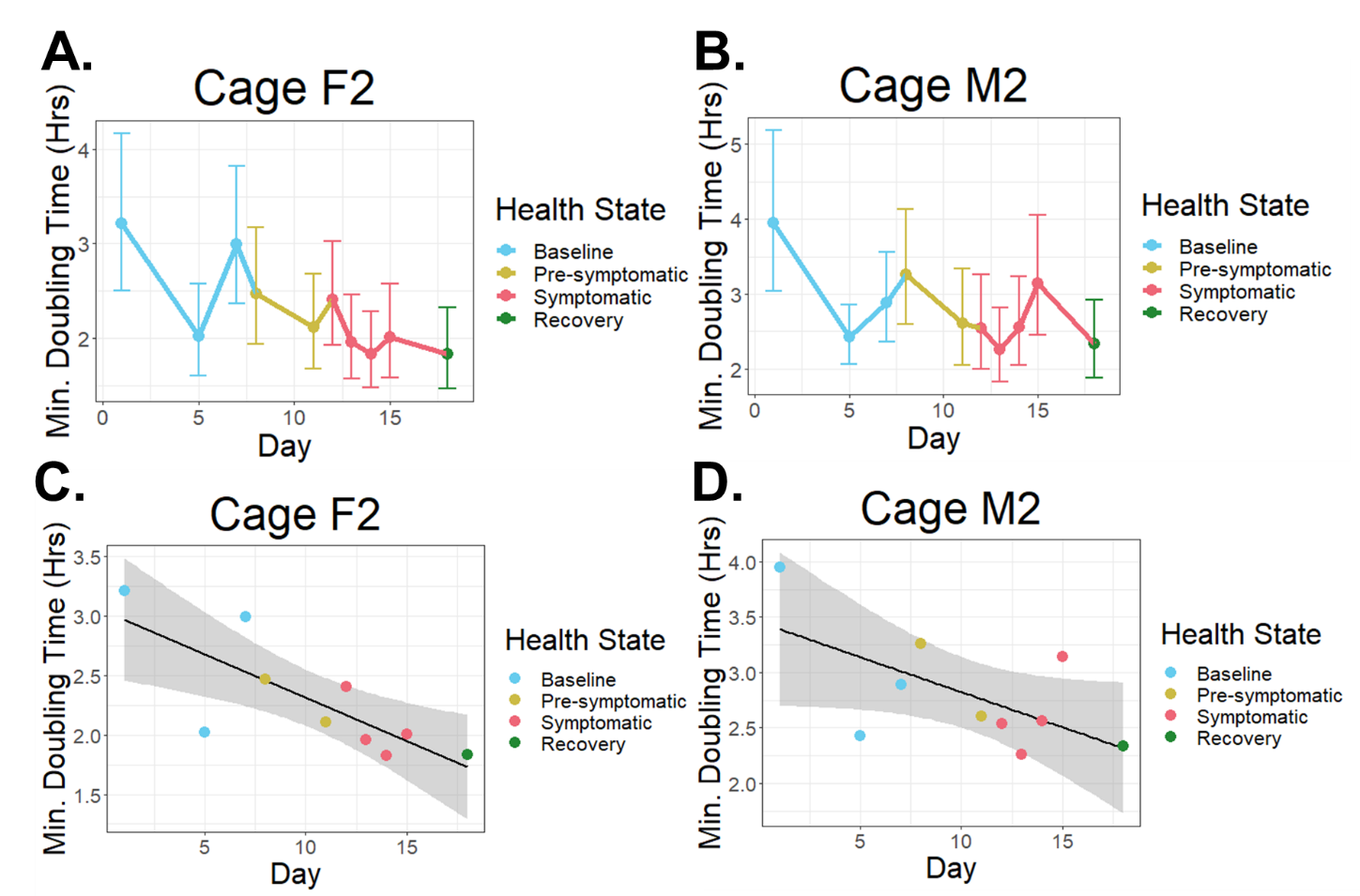

**Supplementary Figure 14**. Minimal doubling time for all bacterial reads from whole genome shotgun (WGS) sequencing data.

Line plots (A-B) and linear regressions (C-D) of the estimated minimal doubling time for the whole bacterial community from the baseline state to the beginning of the recovery state for cage F2 (A, C) and cage M2 (B, D).

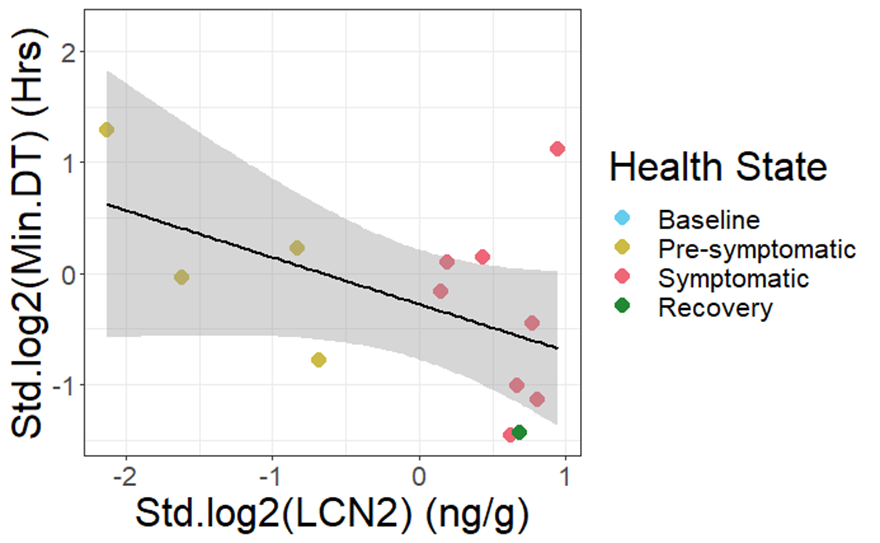

**Supplementary Figure 15**. Correlation between lipocalin-2 and minimal doubling times

Linear mixed effects model for the standardized, log2-transformed values of lipocalin-2 (LCN2) versus minimal doubling time (DT) as calculated with gRodon. Random intercept effects included Day (time effect) and cage (host origin of bacteria). Fixed effects included levels of LCN2.

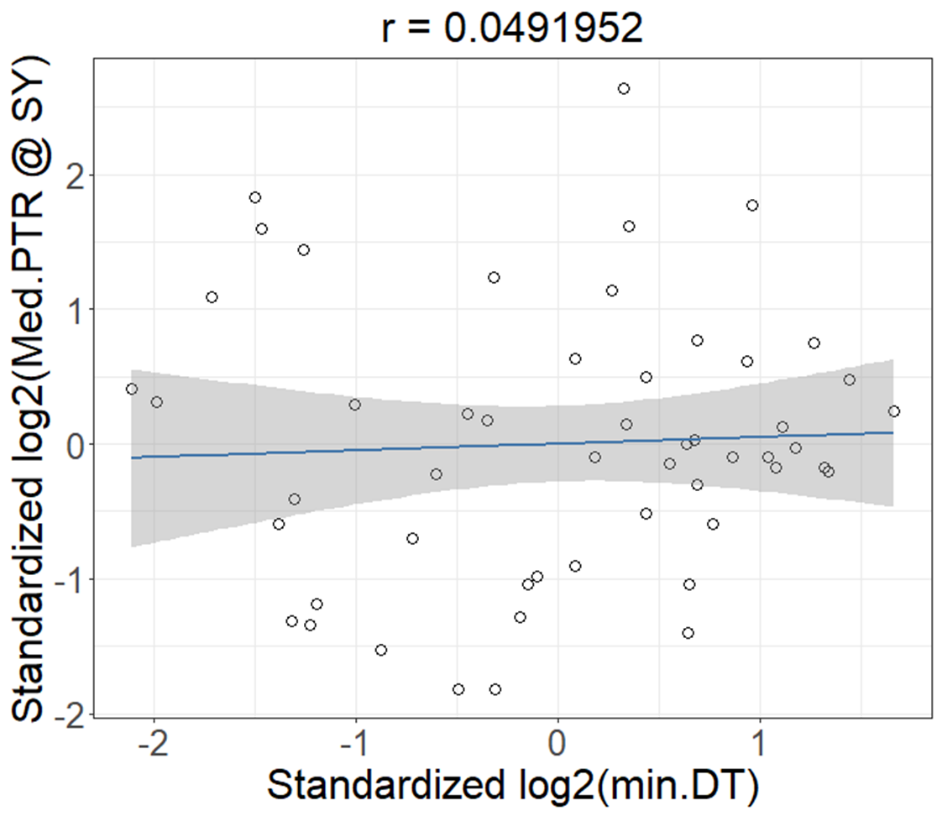

**Supplementary Figure 16**. Correlation between minimal doubling times and replication rates

Pearson correlation between the standardized, log2-transformed estimations of minimum doubling time (calculated from gRodon) and median replication rate (calculated from GRiD) of all recoverable metagnome-assembled genomes (MAGs) during the symptomatic phase.

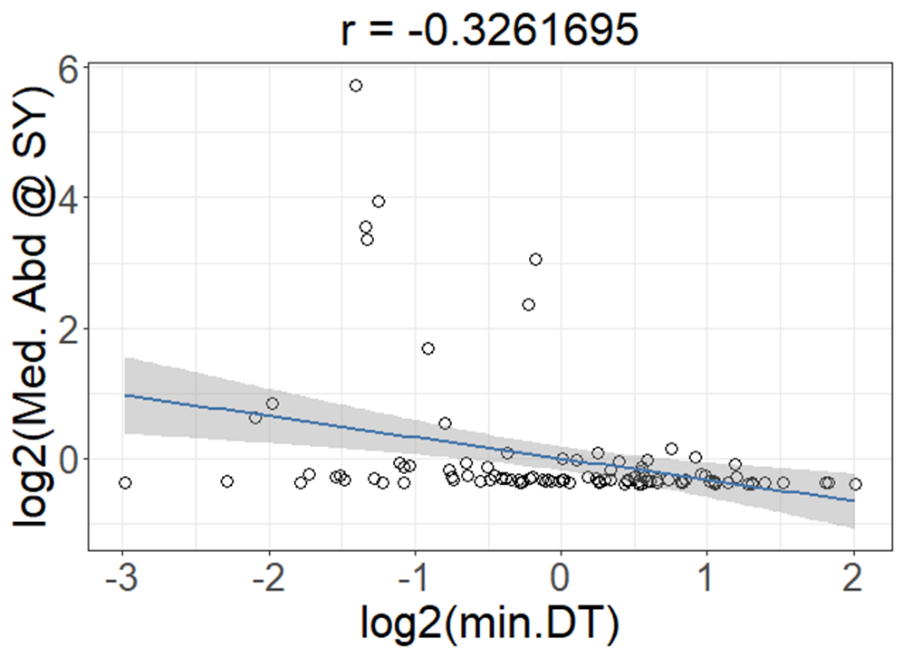

**Supplementary Figure 17**. Correlation between minimal doubling time and relative abundance

Pearson correlation between log2-transformed estimated minimal doubling time (calculated from gRodon) and log2-transformed median relative abundance during the symptomatic state
