## Supplementary Tables for "Dextran sodium sulfate-induced colitis alters the proportion and composition of replicating gut bacteria"

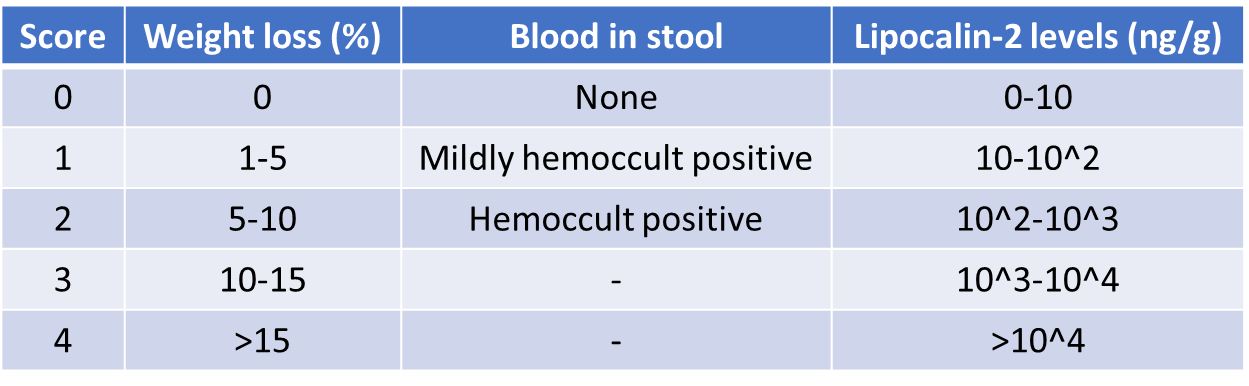

**Supplementary Table 1**. Colitis scoring method.

Scores for weight loss and blood in stool are based on those used by Kim JJ et al. (2012).^28^ Lipocalin-2 levels are expressed as nanograms per gram of feces (ng/g).

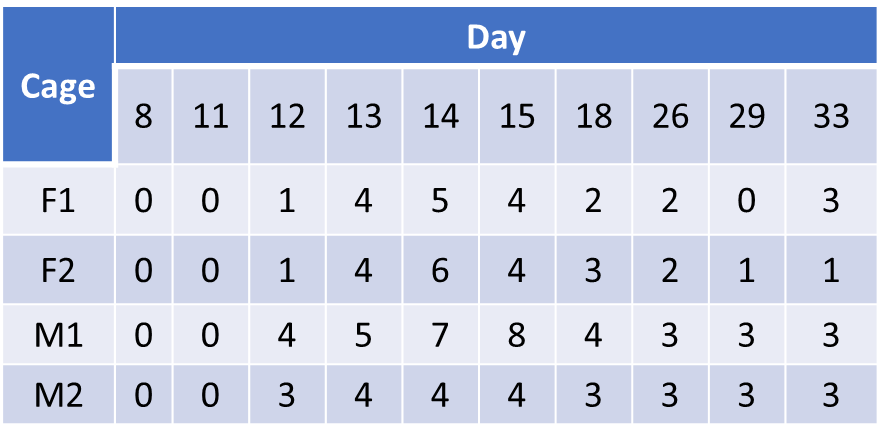

**Supplementary Table 2**. Colitis scores during first DSS experiment.

Colitis scores for each cage of mice during the first DSS experiment, starting one day before DSS administration (day 8) and ending 21 days (3 weeks) after termination of DSS administration (day 33).

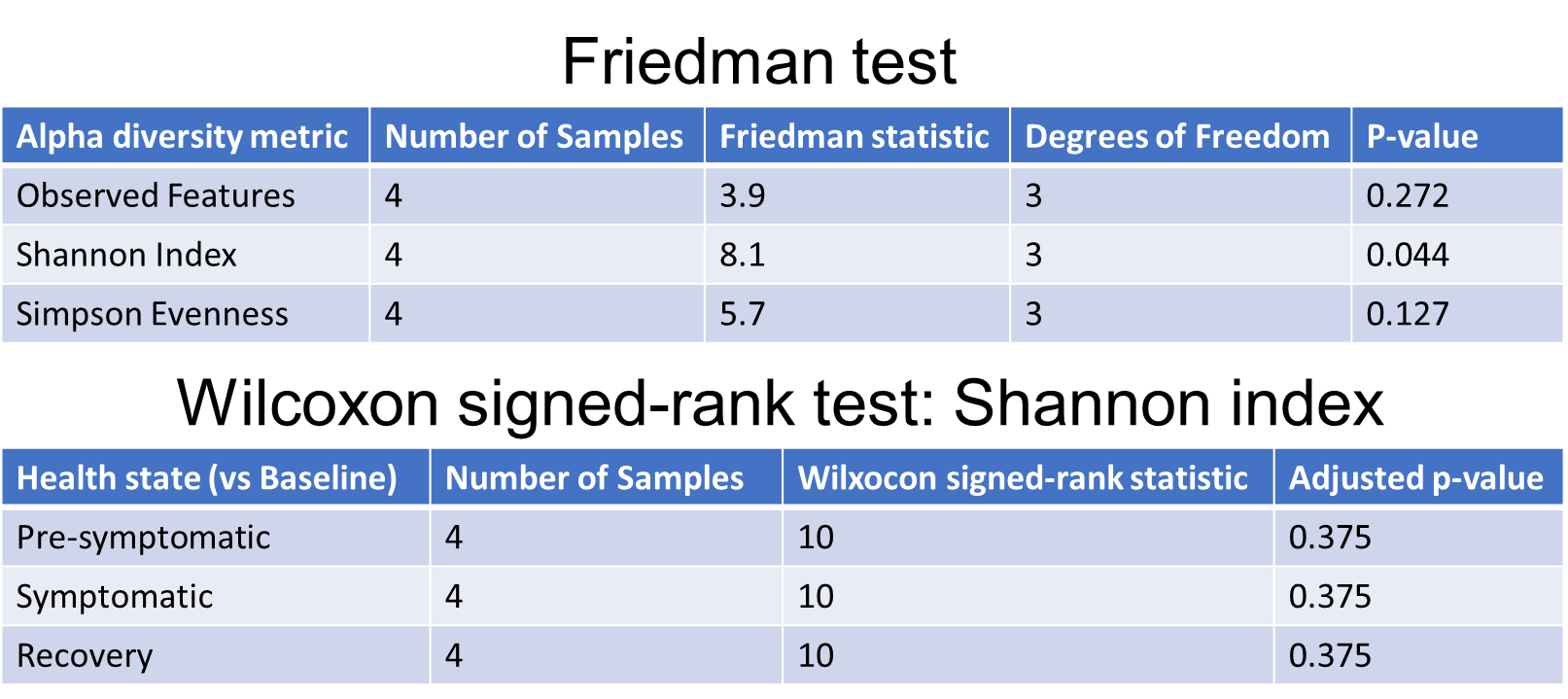

**Supplementary Table 3**. Alpha diversity metric statistics for first DSS mouse experiment.

The Friedman test was conducted by comparing the median alpha diversity values of each cage per health state to the baseline state, to determine whether alpha diversity significantly changed from the baseline state. The post-hoc Wilcoxon signed-rank test was performed using default settings on the Shannon index to determine which health state was responsible for the significant difference.

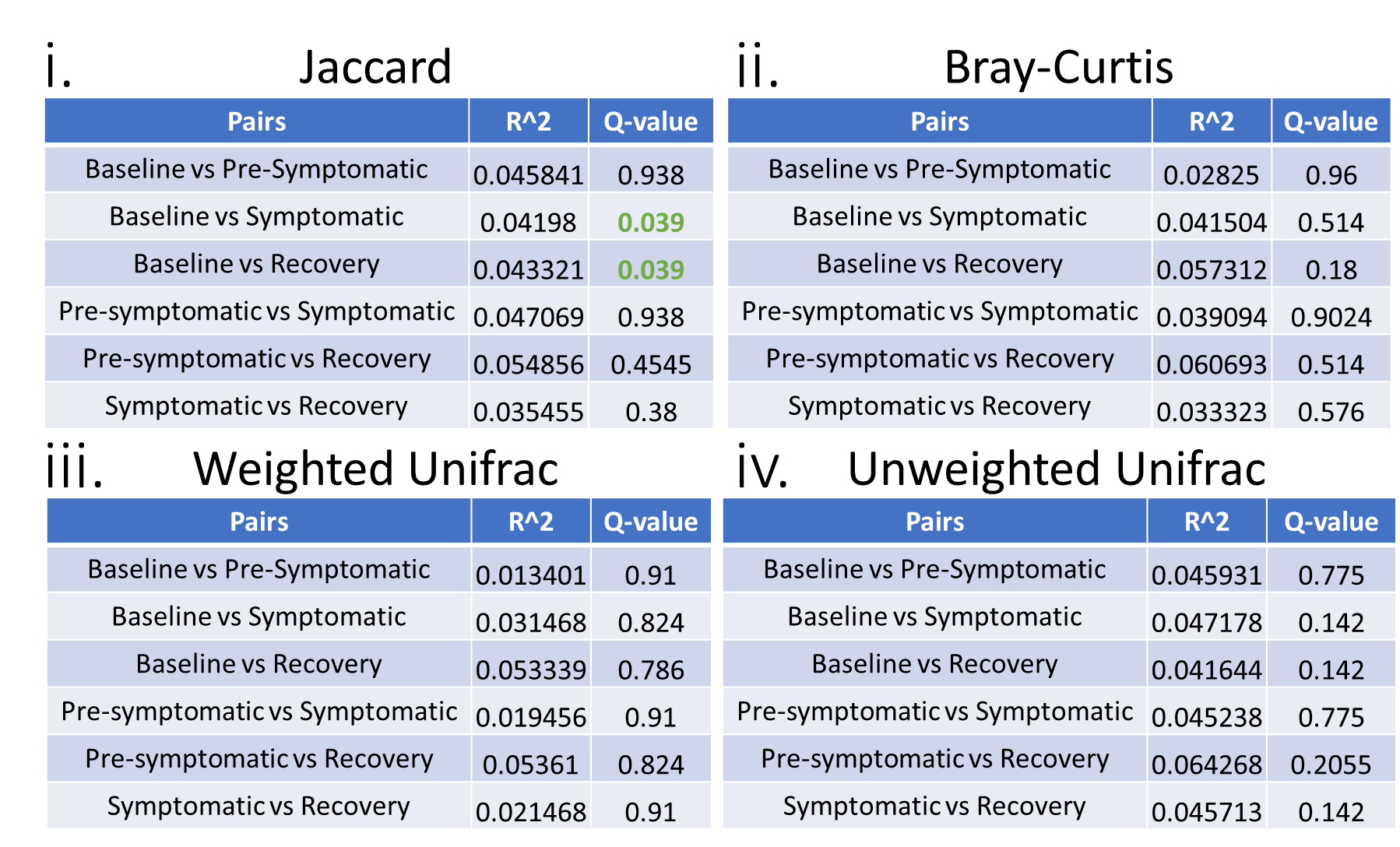

**Supplementary Table 4**. PERMANOVA for all cages during first DSS mouse experiment.

*Jaccard (i), Bray-Curtis (ii), Weighted Unifrac (iii), and Unweighted Unifrac (iv) Green Q-values indicate statistically significant results. .*

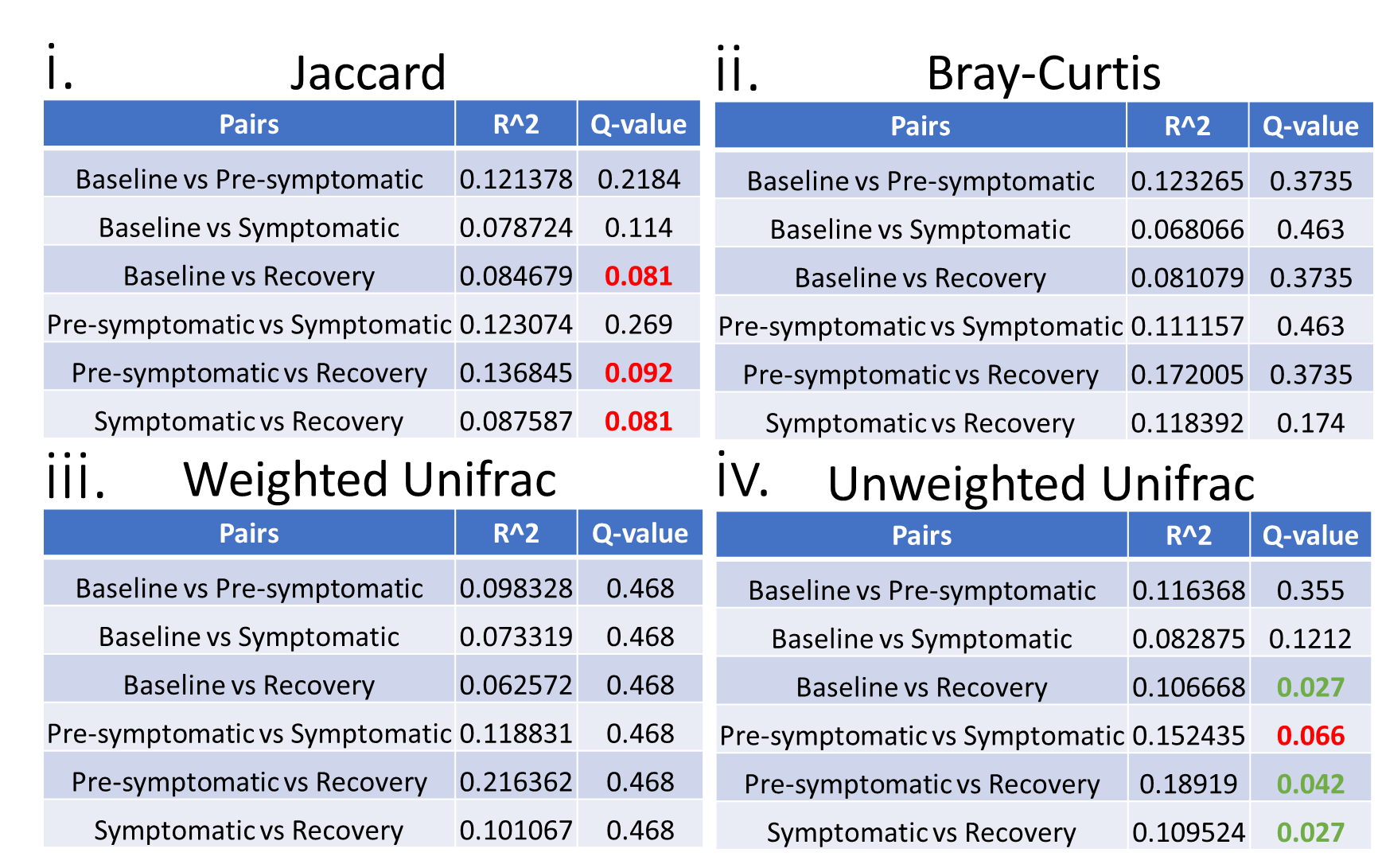

**Supplementary Table 5**. PERMANOVA for all female mice during first DSS mouse experiment.

Jaccard (i), Bray-Curtis (ii), Weighted Unifrac (iii), and Unweighted Unifrac (iv). Green Q-values indicate statistically significant results. Red Q-values indicate near statistically significant results (q >0.05 but <0.1)

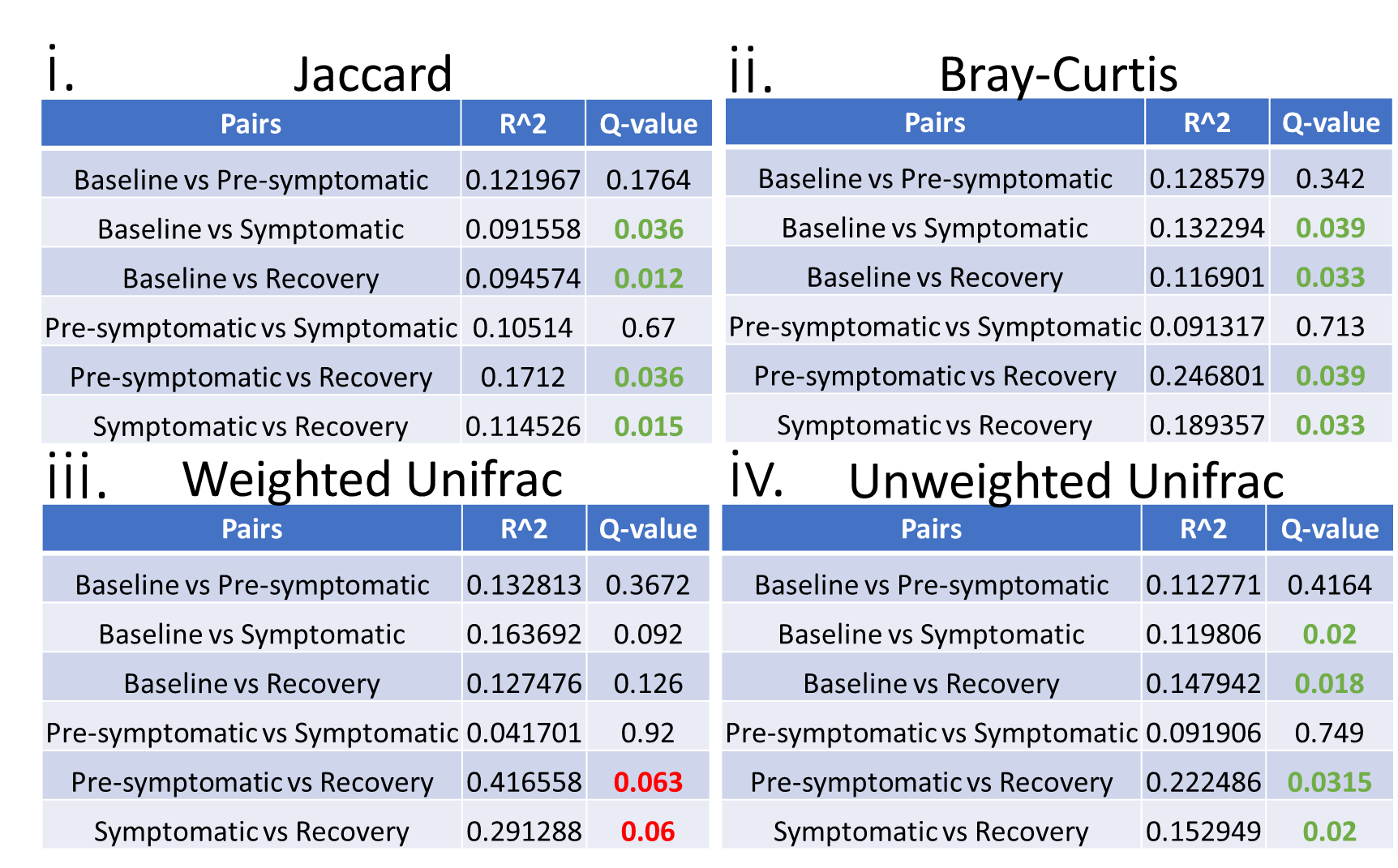

**Supplementary Table 6**. PERMANOVA for all male mice during first DSS mouse experiment.

Jaccard (i), Bray-Curtis (ii), Weighted Unifrac (iii), and Unweighted Unifrac (iv). Green Q-values indicate statistically significant results. Red Q-values indicate near statistically significant results (q >0.05 but <0.1)

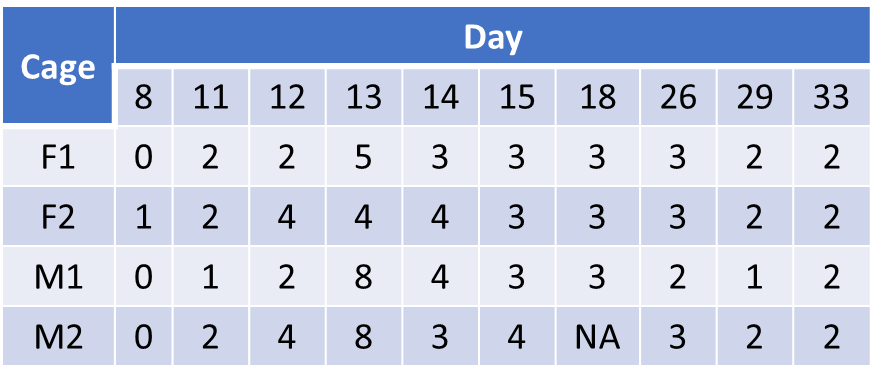

**Supplementary Table 7**. Colitis scores during second DSS experiment.

Colitis scores for each cage of mice during the second DSS experiment, starting one day before DSS administration (day 8) and ending 21 days (3 weeks) after termination of DSS administration (day 33). The NA is because there was not enough stool from this cage at this time point to perform a lipocalin-2 test.

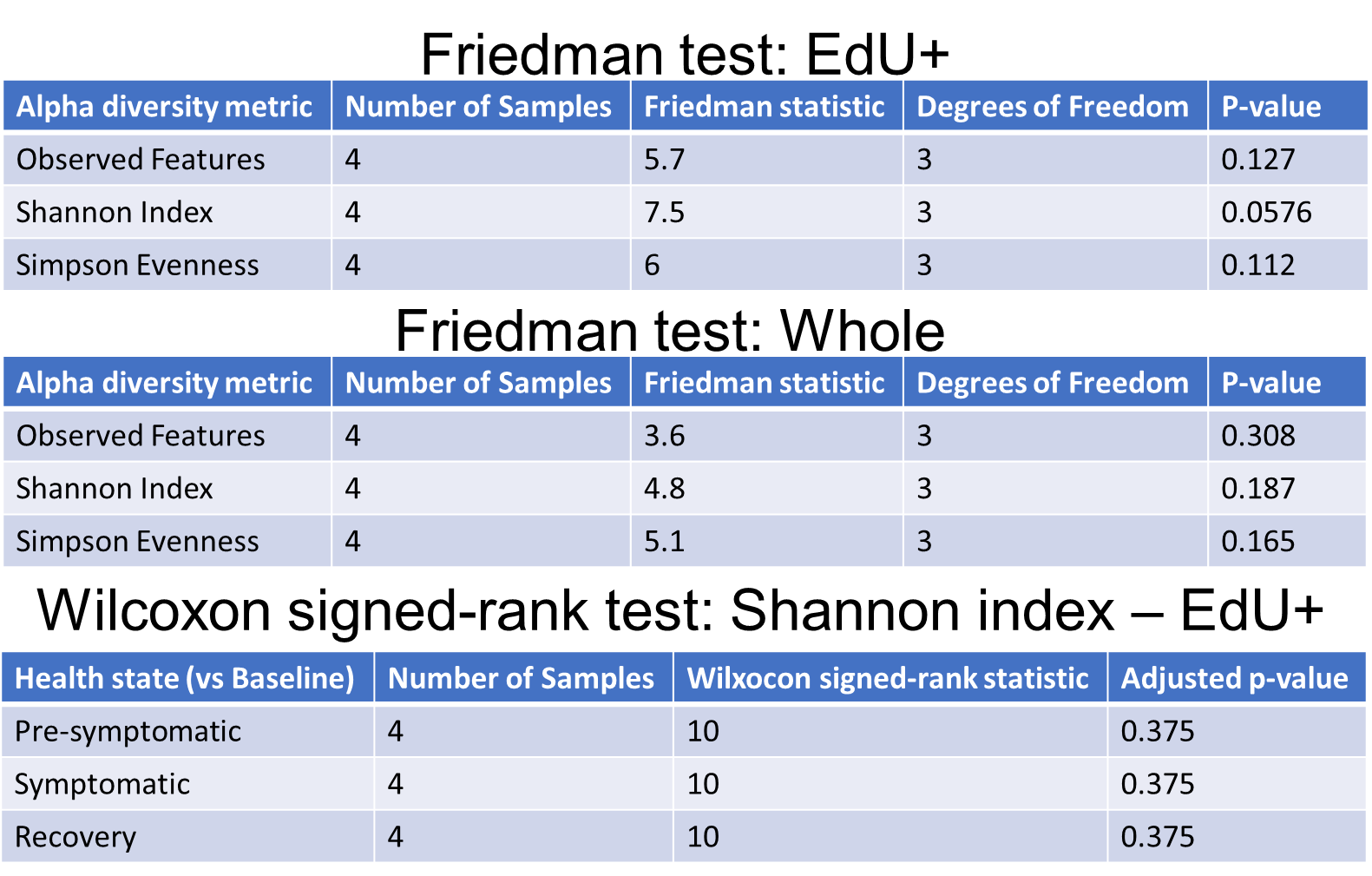

**Supplementary Table 8**. Alpha diversity metric statistics for second DSS mouse experiment.

Alpha diversity metric statistics for the second DSS mouse experiment, for both the replicating (EdU^+^) and whole community (Whole) of bacteria. The Friedman test was conducted by comparing the median alpha diversity values of each cage per health state and per sorted fraction to the baseline state, to determine whether alpha diversity significantly changed from baseline. The post-hoc Wilcoxon signed-rank test was performed using default settings on each alpha diversity metric (observed features, Shannon index, Simpson evenness) to determine which health state was responsible for the significant difference, where relevant.

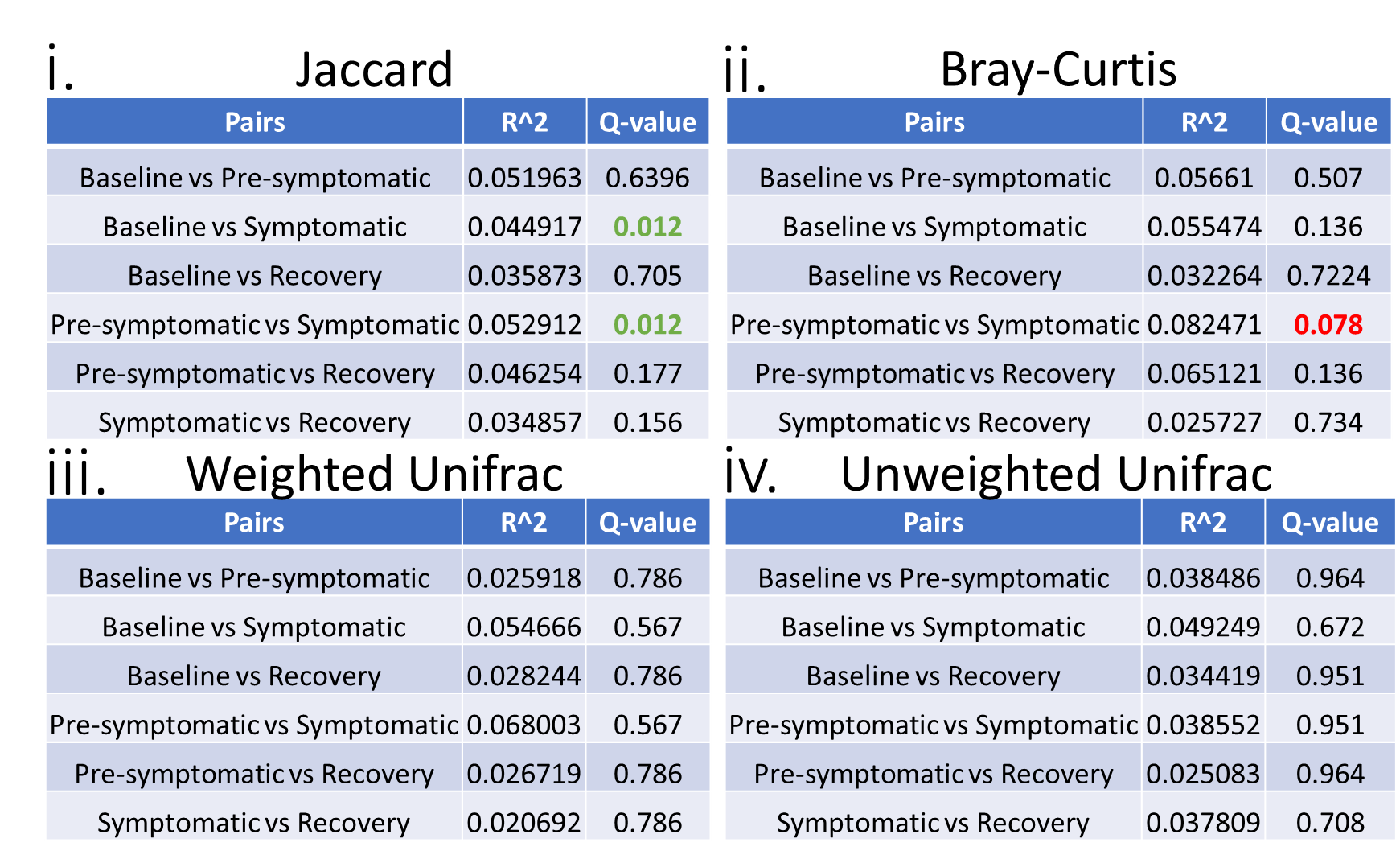

**Supplementary Table 9**. PERMANOVA for replicating (EdU^+^) bacteria.

PERMANOVA analyses for replicating (EdU^+^) bacterial cells from all cages combined during the second DSS mouse experiment. Jaccard (i), Bray-Curtis (ii), Weighted Unifrac (iii), and Unweighted Unifrac (iv). Green Q-values indicate statistically significant results. Red Q-values indicate near statistically significant results (q >0.05 but <0.1)

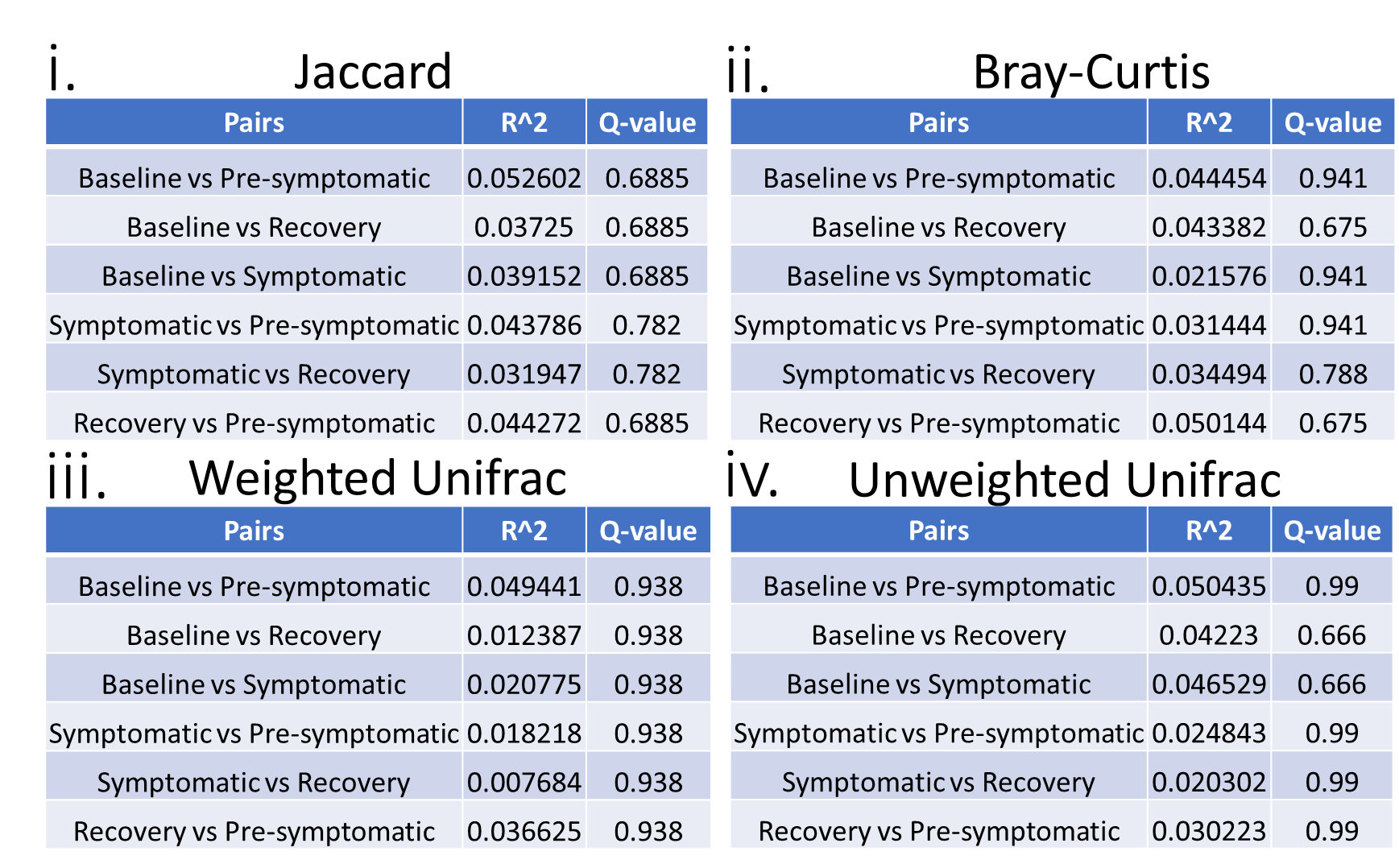

**Supplementary Table 10**. PERMANOVA for whole bacterial community

Results from PERMANOVA analysis for the whole community of bacterial cells from all cages combined during the second DSS mouse experiment. Jaccard (i), Bray-Curtis (ii), Weighted Unifrac (iii), and Unweighted Unifrac (iv).

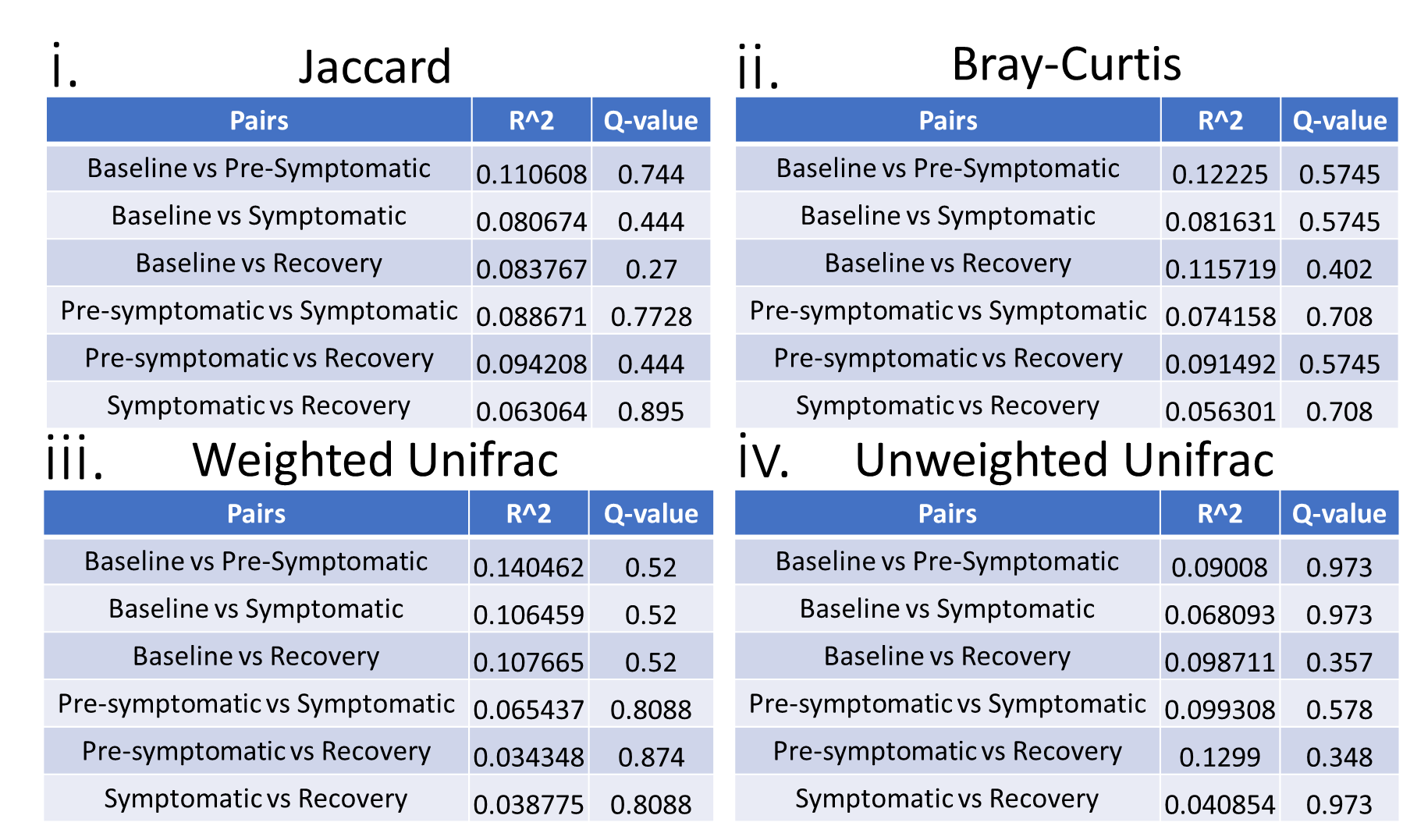

**Supplementary Table 11**. PERMANOVA for replicating (EdU^+^) bacteria from female mice

Results from PERMANOVA analysis for the replicating (EdU^+^) bacterial cells from cages of female mice during the second DSS mouse experiment. Jaccard (i), Bray-Curtis (ii), Weighted Unifrac (iii), and Unweighted Unifrac (iv).

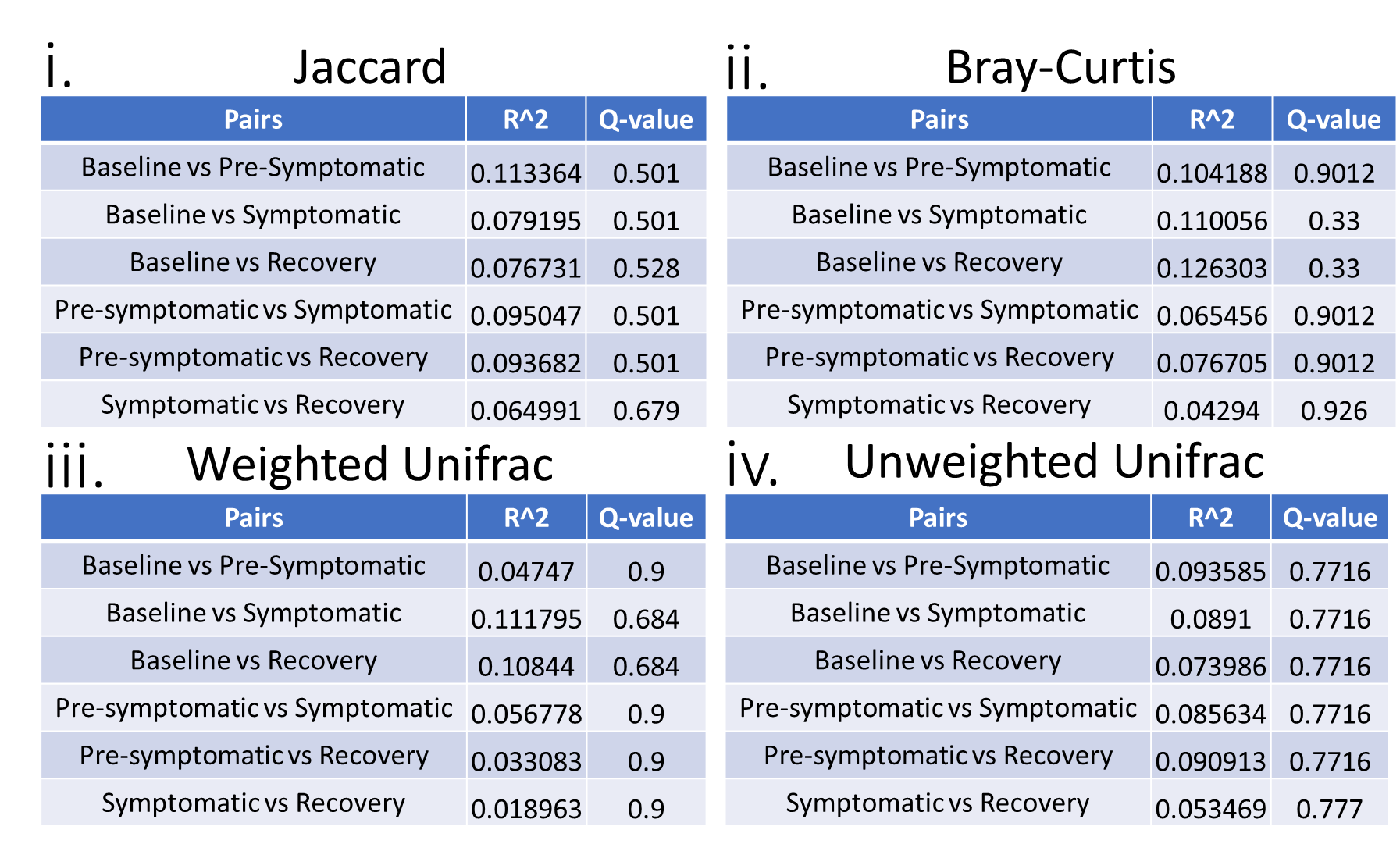

**Supplementary Table 12**. PERMANOVA for whole community of bacteria from female mice

Results from PERMANOVA analysis for the whole community of bacterial cells from cages of female mice during the second DSS mouse experiment. Jaccard (i), Bray-Curtis (ii), Weighted Unifrac (iii), and Unweighted Unifrac (iv).

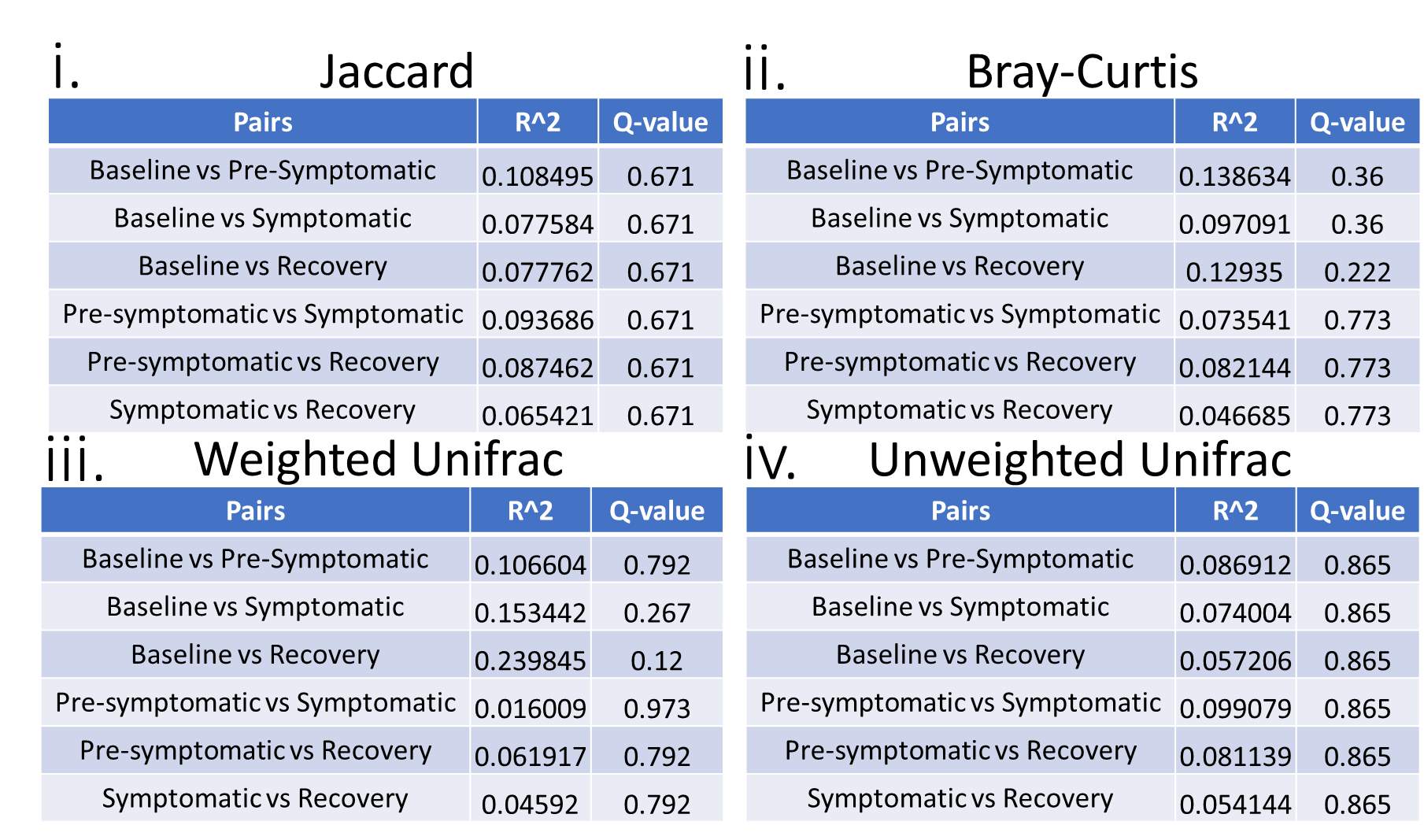

**Supplementary Table 13**. PERMANOVA for replicating (EdU^+^) bacteria from male mice

Results from PERMANOVA analysis for the replicating (EdU^+^) bacterial cells from cages of male mice during the second DSS mouse experiment. Jaccard (i), Bray-Curtis (ii), Weighted Unifrac (iii), and Unweighted Unifrac (iv).

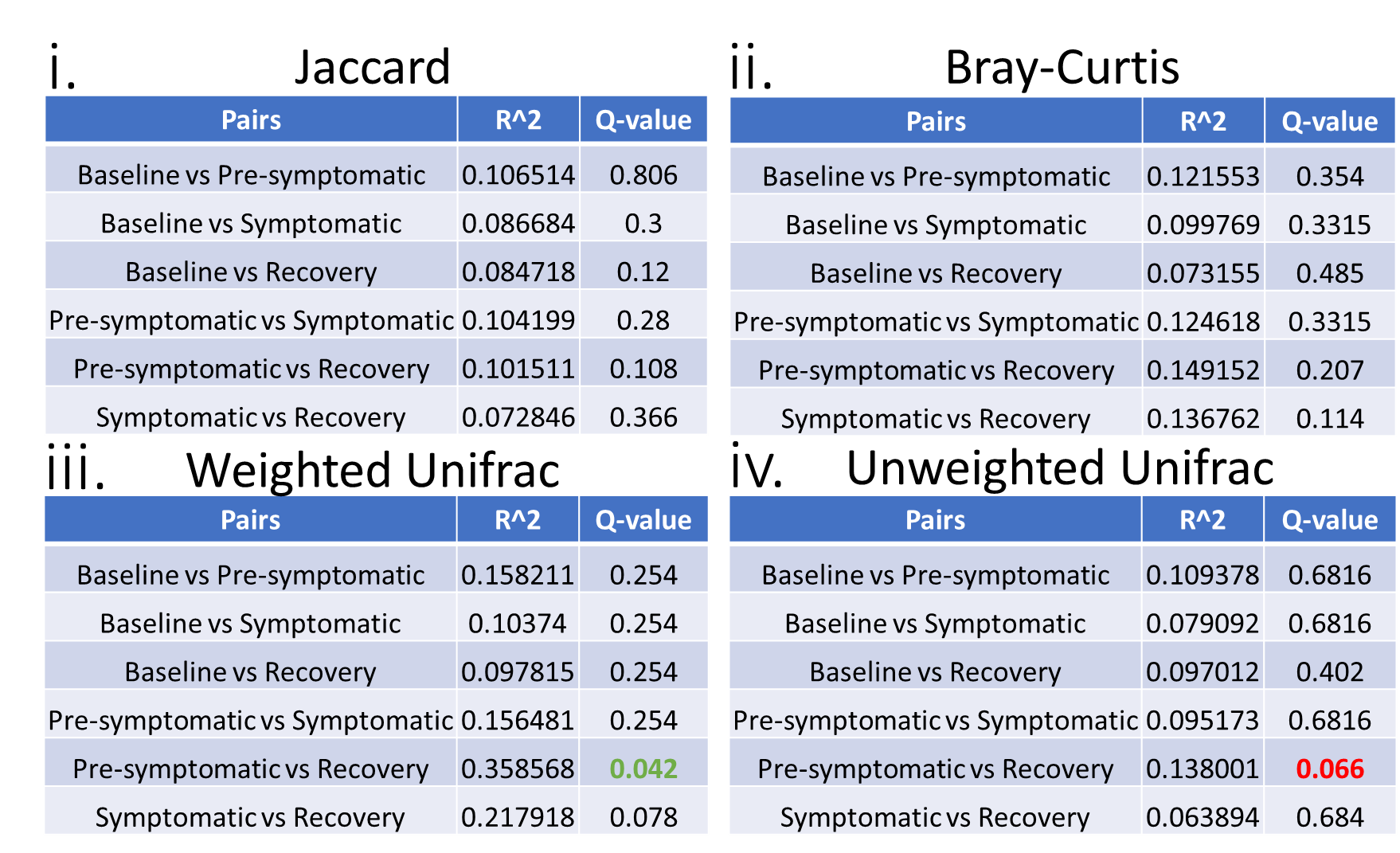

**Supplementary Table 14**. PERMANOVA for whole bacterial community from male mice

Results from PERMANOVA analysis for the whole community of bacterial cells from cages of male mice during the second DSS mouse experiment. Jaccard (i), Bray-Curtis (ii), Weighted Unifrac (iii), and Unweighted Unifrac (iv). Green Q-values indicate statistically significant results. Red Q-values indicate near statistically significant results (q >0.05 but <0.1)

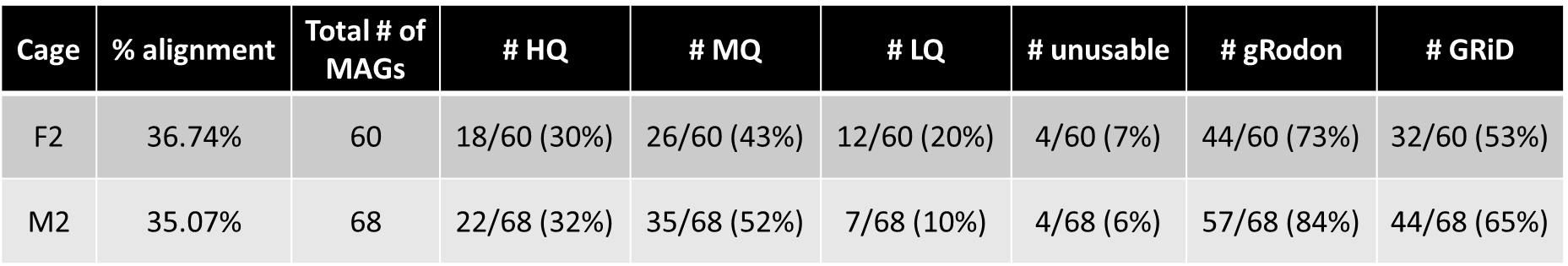

**Supplementary Table 15**. Metagenome-assembled genomes (MAGs) statistics from second DSS experiment.

Percent (%) alignment denotes the percent of all the DNA sequences from the experiment which were captured by assembling these bacterial genomes. HQ = high quality, MQ = medium quality, LQ = low quality, according to the MIMAGS definition of MAG quality (Bowers R.M. et al. 2017).^158^ The quantity and percentage of the MAGs which could be used for gRodon (# gRodon) and GRiD (# GRiD) are additionally quantified.

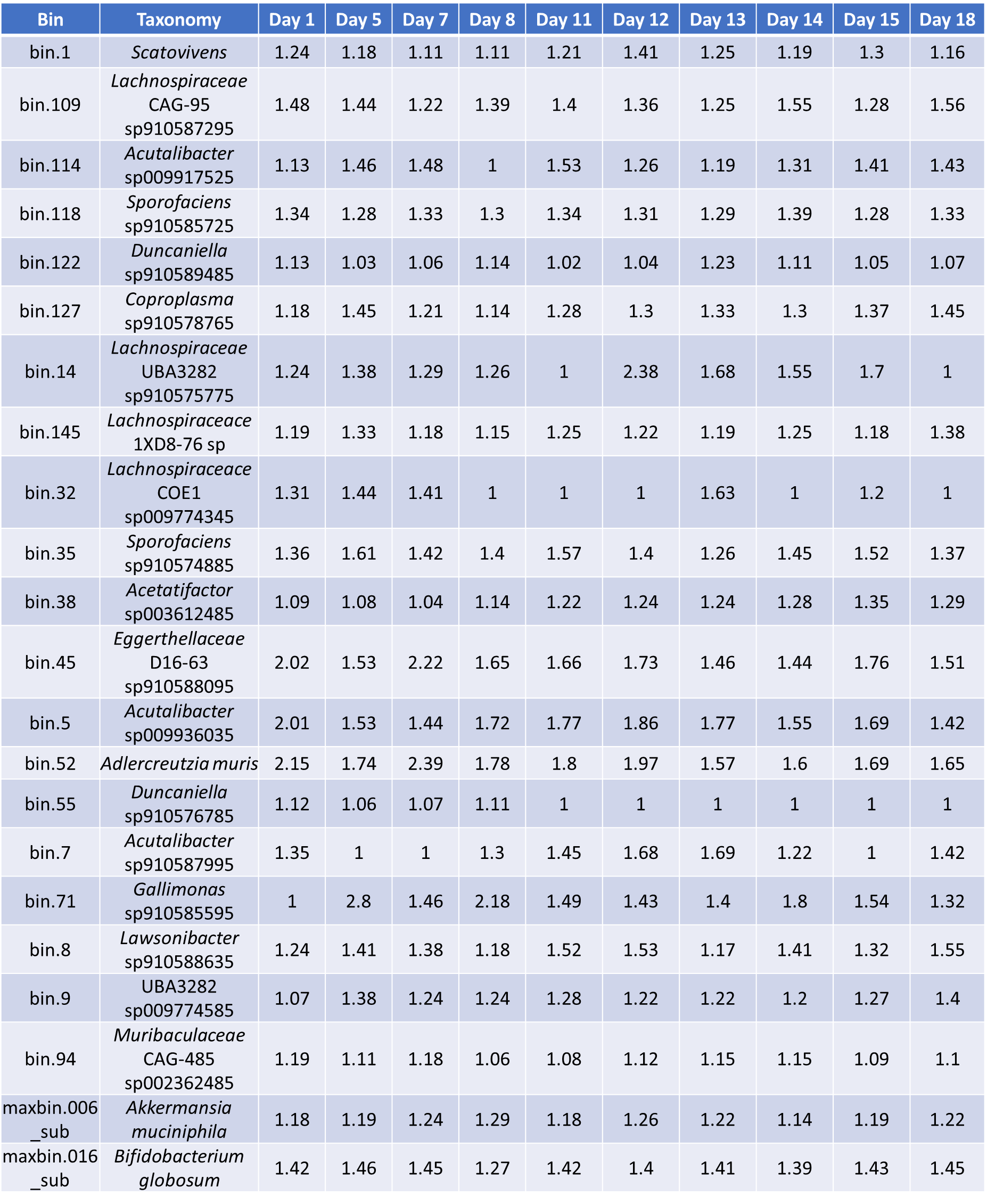

**Supplementary Table 16**. GRiD replication rates – cage F2.

Replication rate values from GRiD for all recoverable metagenome-assembled genomes (MAGs) at all measured time points from cage F2

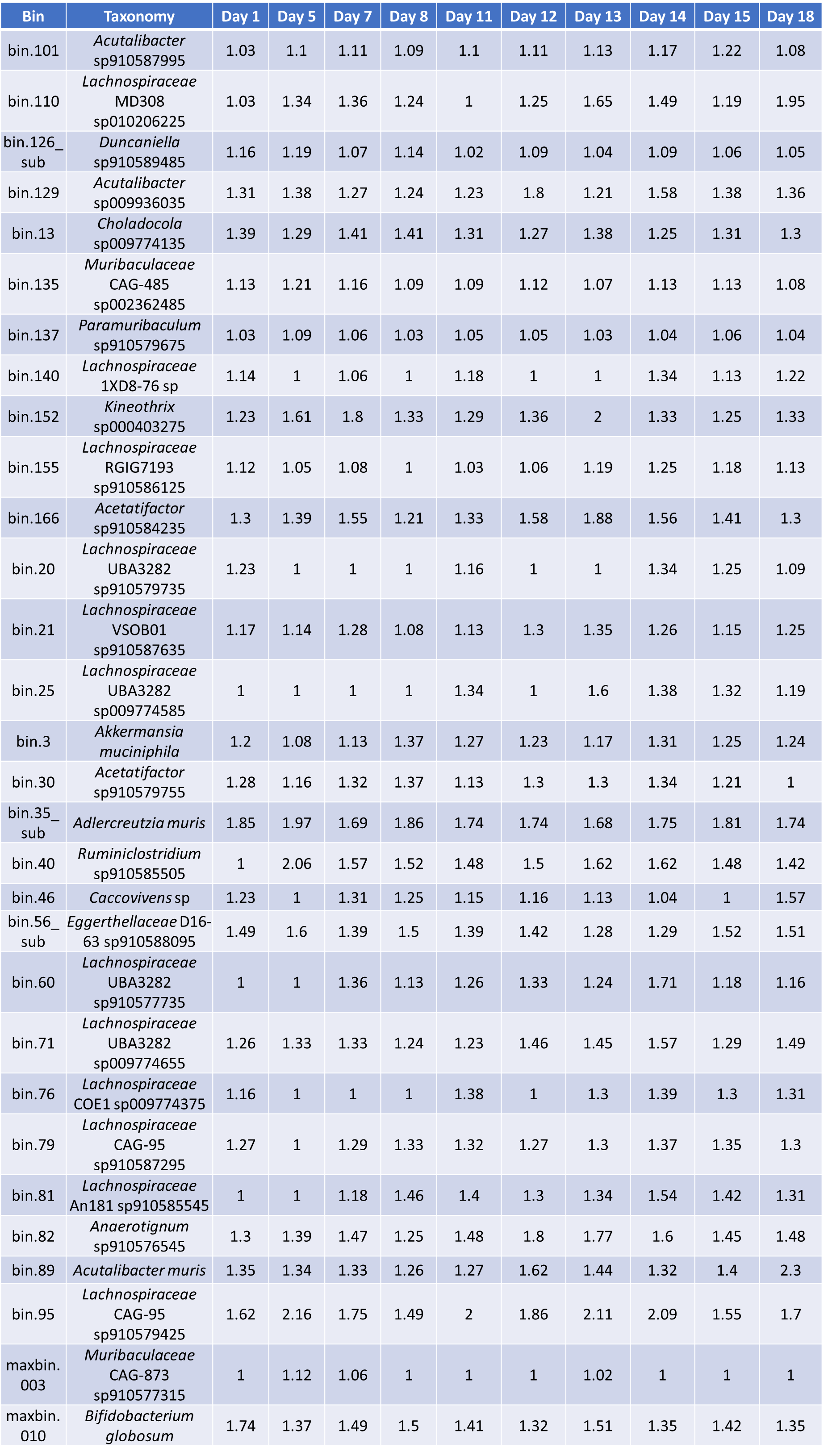

**Supplementary Table 17**. GRiD replication rates – cage M2.

Replication rate values from GRiD for all recoverable metagenome-assembled genomes (MAGs) at all measured time points from cage M2

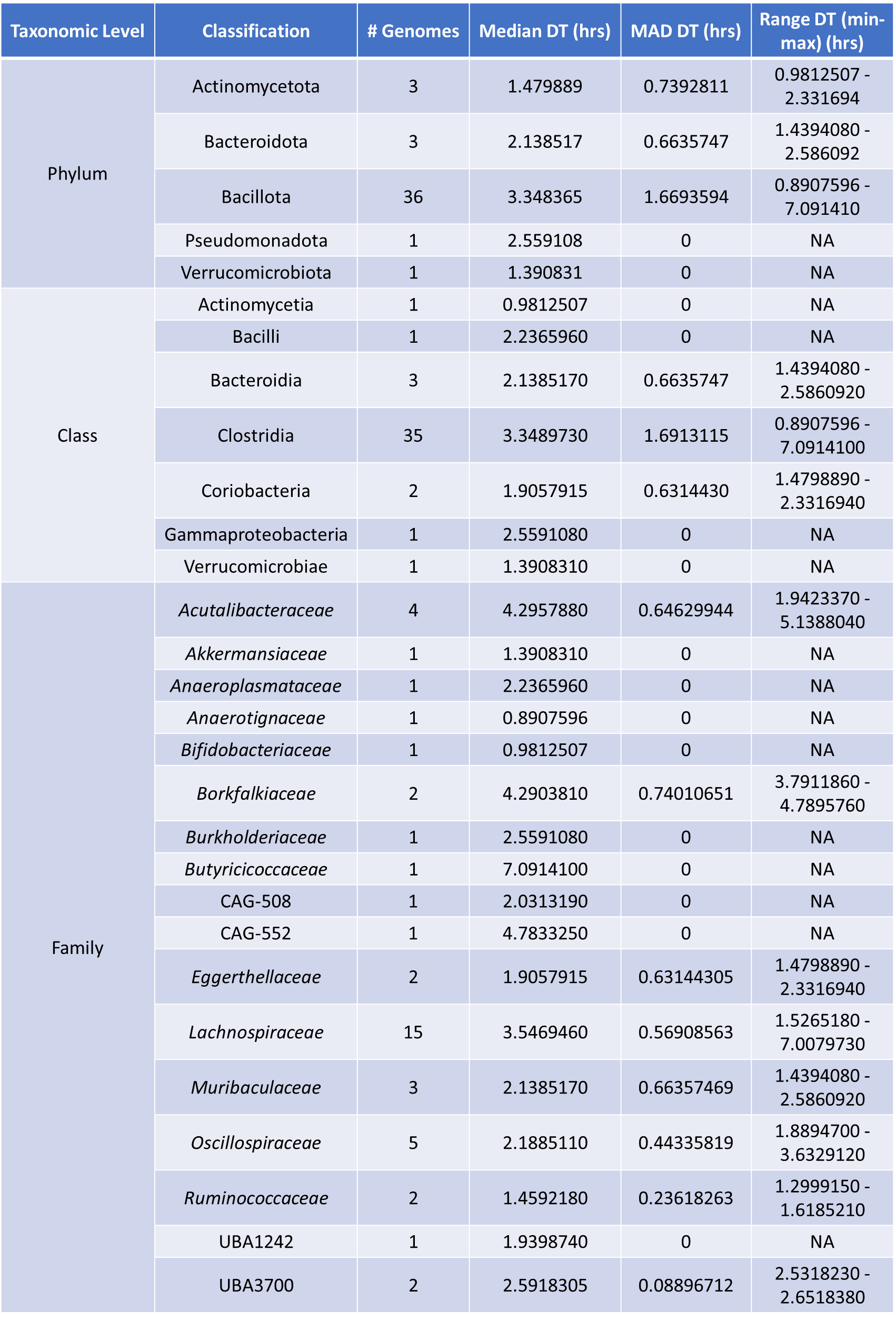

**Supplementary Table 18**. gRodon minimal doubling times – cage F2.

Statistics on metagenome-assembled genomes (MAGs) with calculated minimal doubling times for cage F2. Number of metagenome-assembled genomes (MAGs) (# Genomes) at a certain taxonomic level, median doubling time (DT) and median absolute deviation (MAD) of minimal DTs calculated by gRodon at that taxonomic level, and range of minimal DT values at that taxonomic level.

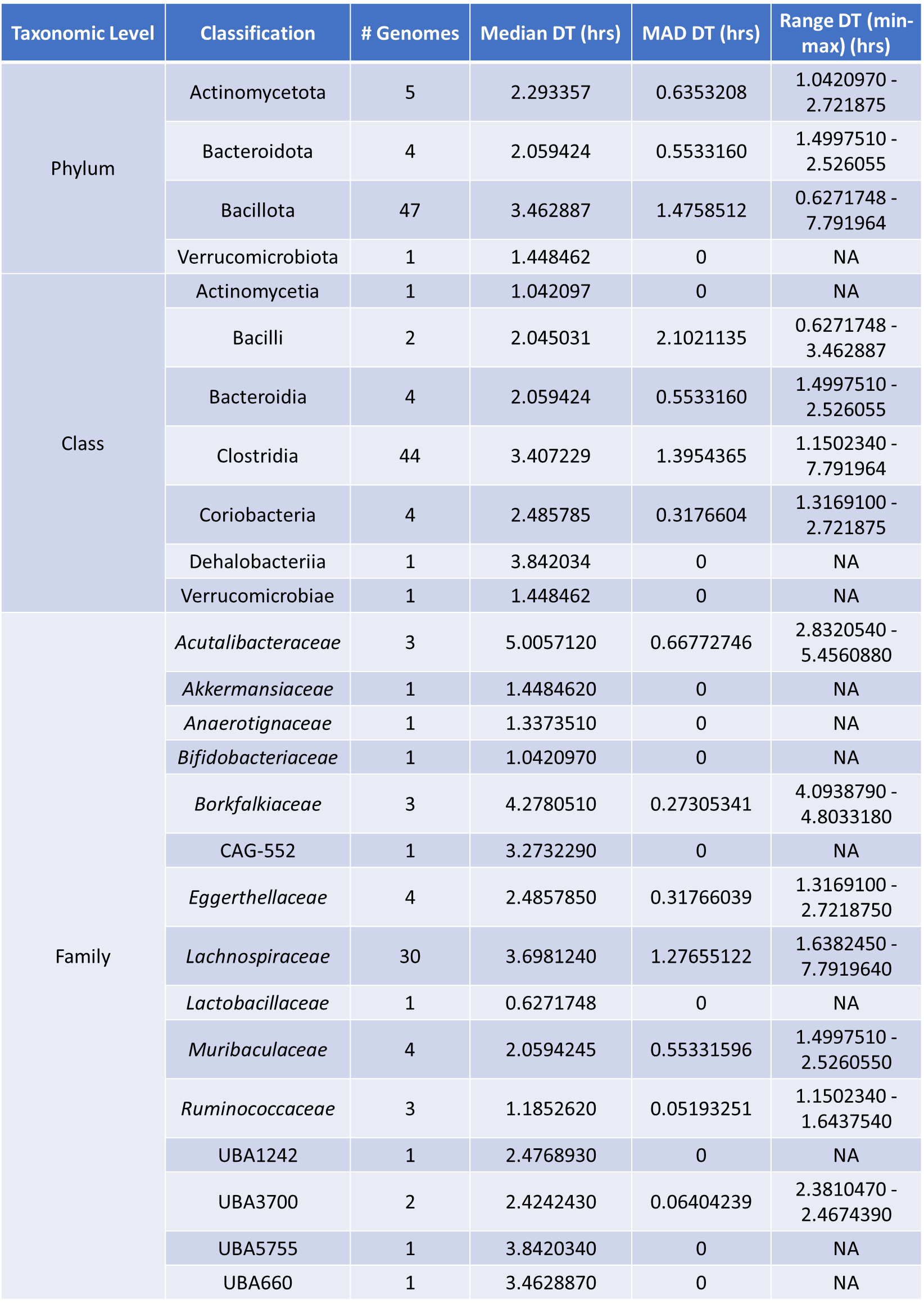

**Supplementary Table 19**. gRodon minimal doubling times – cage M2.

Statistics on metagenome-assembled genomes (MAGs) with calculated minimal doubling times for cage M2. Number of metagenome-assembled genomes (MAGs) (# Genomes) at a certain taxonomic level, median doubling time (DT) and median absolute deviation (MAD) of minimal DTs calculated by gRodon at that taxonomic level, and range of minimal DT values at that taxonomic level.

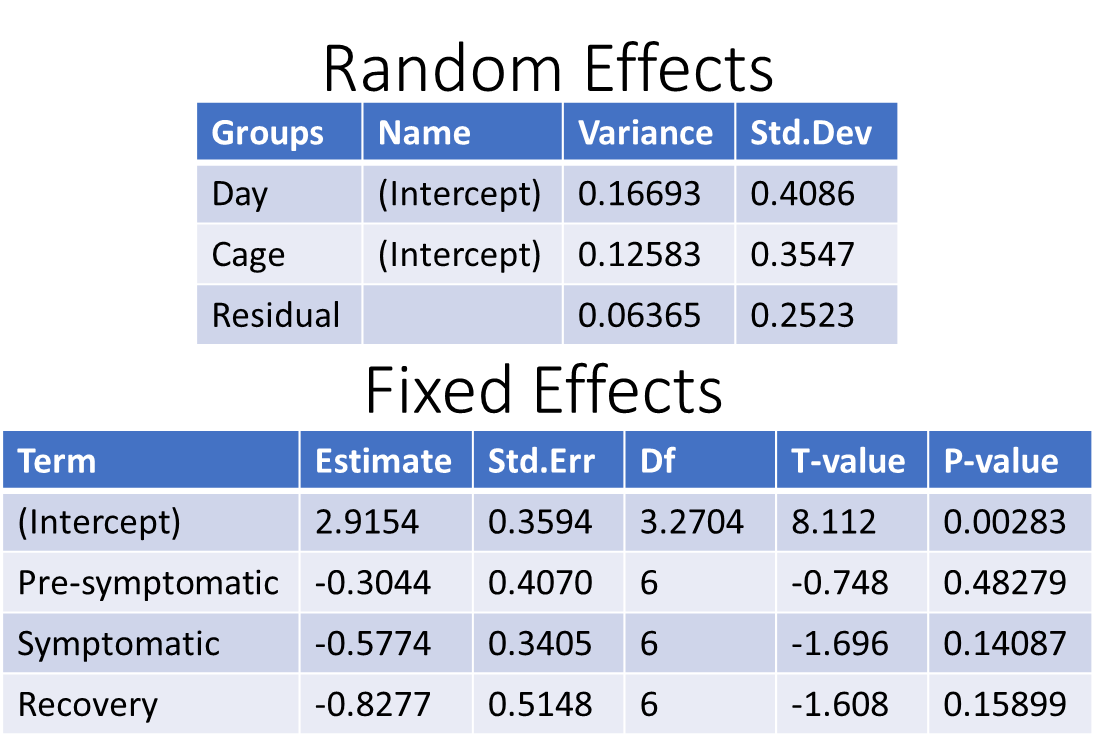

**Supplementary Table 20**. Linear regression statistics – Figure 3.5E.

Statistics on the random effects and fixed effects for the linear regression performed in Figure 3.5E

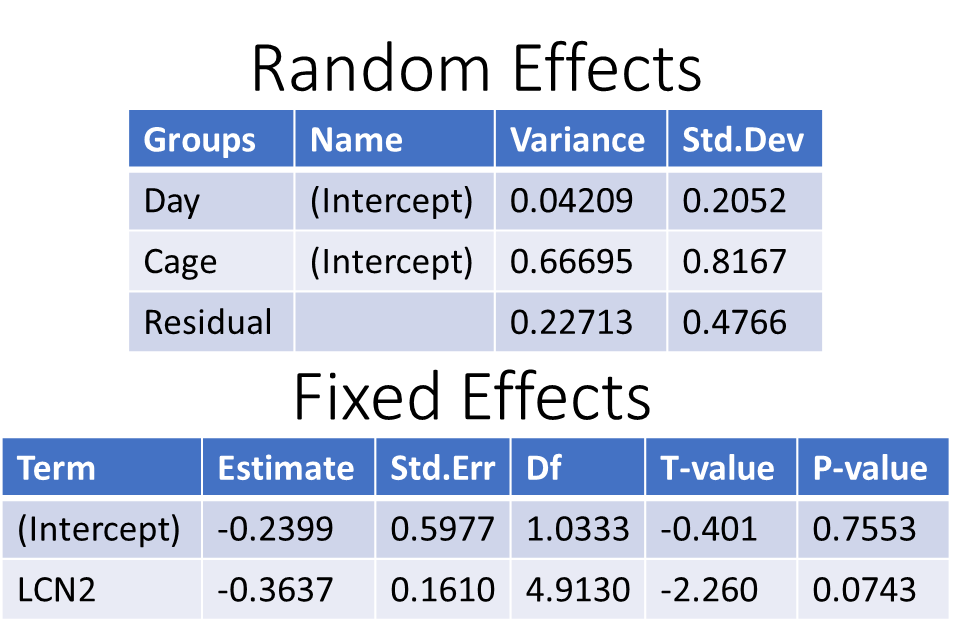

**Supplementary Table 21**. Linear regression statistics – Supplementary Figure 3.15.

Statistics on the random effects and fixed effects for the linear regression performed in Supplementary Figure 3.15.
